# Dynamics of diversified A-to-I editing in *Streptococcus pyogenes* is governed by changes in mRNA stability

**DOI:** 10.1101/2023.09.19.555891

**Authors:** Thomas F. Wulff, Karin Hahnke, Anne-Laure Lécrivain, Katja Schmidt, Rina Ahmed-Begrich, Knut Finstermeier, Emmanuelle Charpentier

**Affiliations:** Max Planck Unit for the Science of Pathogens, 10117 Berlin, Germany; Institute for Biology, Humboldt University Berlin, 10115 Berlin, Germany

**Author notes:** Anne-Laure Lécrivain, Berlin Institute for Medical Systems Biology (BIMSB), Max Delbrück Center for Molecular Medicine in the Helmholtz Association (MDC), 10115 Berlin, Germany.

## Abstract

Adenosine-to-inosine (A-to-I) RNA editing plays an important role in the post-transcriptional regulation of eukaryotic cell physiology. However, our understanding of the occurrence, function and regulation of A-to-I editing in bacteria remains limited. Bacterial mRNA editing is catalysed by the deaminase TadA, which was originally described to modify a single tRNA in *E. coli*. Intriguingly, several bacterial species appear to perform A-to-I editing on more than one tRNA. Here, we provide evidence that in the human pathogen *Streptococcus pyogenes*, tRNA editing has expanded to an additional tRNA substrate. Using RNA sequencing, we identified more than 27 editing sites in the transcriptome of *S. pyogenes* SF370 and demonstrate that the adaptation of *S. pyogenes* TadA to a second tRNA substrate has also diversified the sequence context and recoding scope of mRNA editing. Based on the observation that editing is dynamically regulated in response to several infection-relevant stimuli, such as oxidative stress, we further investigated the underlying determinants of editing dynamics and identified mRNA stability as a key modulator of A-to-I editing. Overall, our findings reveal the presence and diversification of A-to-I editing in *S. pyogenes* and provide novel insights into the plasticity of the editome and its regulation in bacteria.

**GRAPHICAL ABSTRACT:** 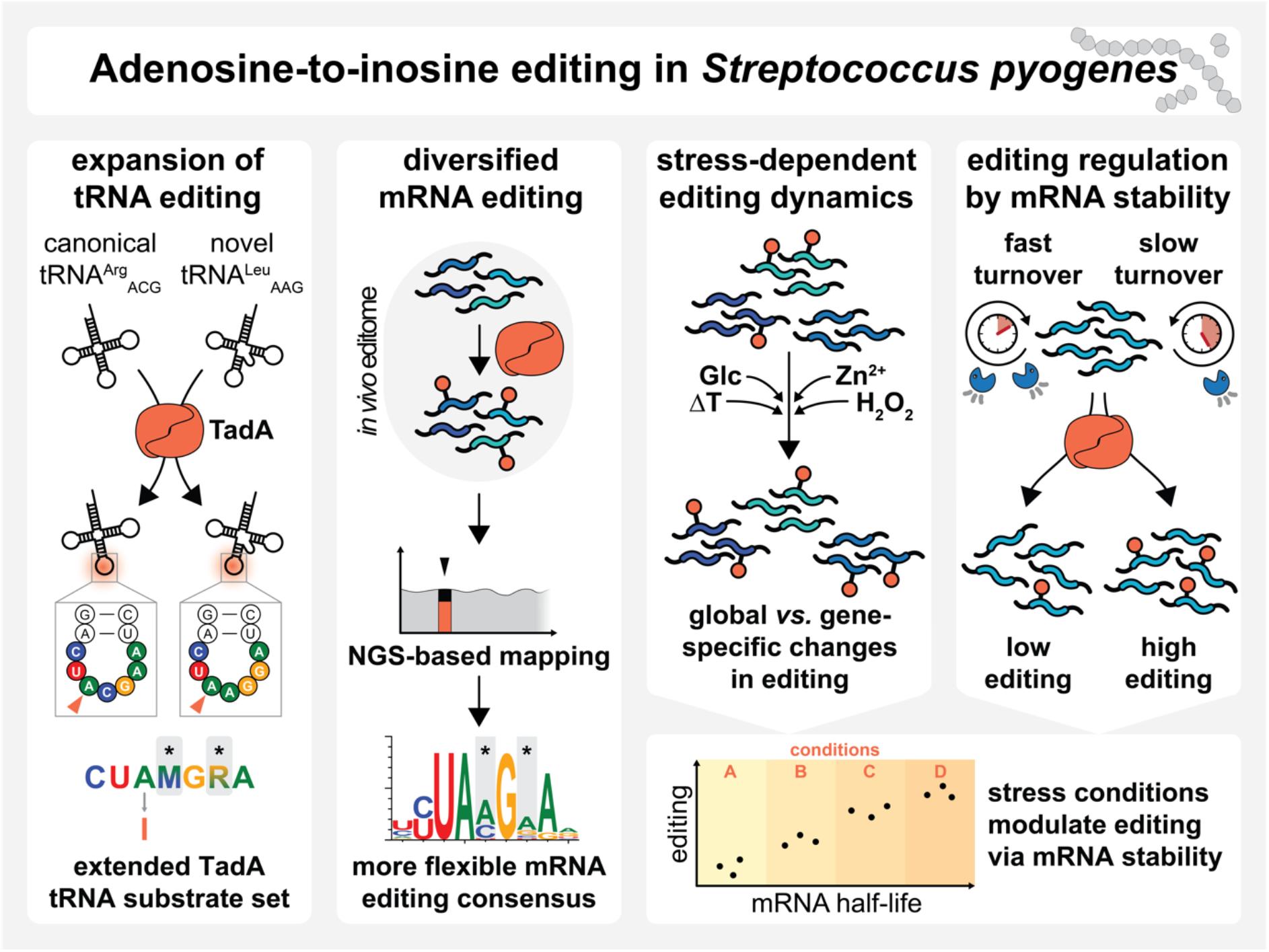

## INTRODUCTION

RNA modifications have emerged as important players in regulatory networks and potent modulators of diverse physiological processes (1). Inosine, a modification derived from adenosine, was first discovered in yeast tRNA^Ala^ and soon found to be present in tRNAs from all domains of life (2–4). In both bacteria and eukaryotes, the presence of inosine at the tRNA wobble position 34 enables modified tRNAs to read A-, C- and U-ending codons (4, 5). While seven to eight tRNAs undergo A34-to-I34 editing in eukaryotes, only a single tRNA (tRNA^Arg^_ACG_) is edited in the model bacterium *Escherichia coli* (4). The increase in the numbers of A34-to-I34 edited tRNAs in eukaryotes correlates with distinct changes in codon usage and tRNA gene content, whereas in bacteria, U34 modifications have been favoured instead (6). Although the selective pressure is far from being understood, positive effects on translation efficiency and fidelity have been discussed (6–9).

The modification of A34 to I34 is catalysed by dedicated tRNA adenosine deaminases. In eukaryotes, the heterodimeric adenosine deaminase acting on tRNA 2 and 3 (ADAT2/3) complex modifies full-length tRNAs with different anticodon loop sequences (4, 10–12). In contrast, the bacterial homologue, the homodimeric tRNA adenosine deaminase TadA, acts on a minimal anticodon arm-like structure, but displays strict sequence specificity in accordance with the anticodon loop sequence of its single target tRNA^Arg^_ACG_ (13, 14). Studies in *E. coli* suggest that bacterial A34-to-I34 editing is essential (4, 13), but some bacterial species, such as *Mycoplasma* spp., have evolved alternative decoding patterns that do not rely on tRNA^Arg^_ICG_ and TadA (15). Conversely, several Firmicutes have recently been shown to harbour additional A34-tRNAs, including *Streptococcus pyogenes* (8). Observations in *Oenococcus oeni* suggest that at least some of these additional A34-tRNAs are edited *in vivo* (9). However, the consequences of expanded A34-to-I34 editing in bacteria remain to be explored.

In addition to its function at the tRNA wobble position, inosine is also present in messenger RNA (mRNA) or other low-abundant non-coding RNA (16). In animals, A-to-I editing is catalysed by members of the adenosine deaminase acting on RNA (ADAR) family (17) and exerts a wealth of physiological functions, *e.g.* in development and immunity (18). Protein recoding, the most prominent effect of RNA editing, relies on the interpretation of inosine as guanosine by the translating ribosome and the subsequent alteration of the amino acid identity encoded in the genome (16). In addition to direct effects on mRNA translation, A-to-I editing also affects RNA maturation, localisation, and interference, and is a crucial modulator of the immune response against double-stranded RNA in animals (16, 19, 20).

Although A-to-I editing is widely recognised for its functions in animals, its occurrence in bacterial RNA species other than tRNA has only recently been established (21). Whereas animals employ dedicated enzyme sets for tRNA and mRNA editing (ADAT2/3 and ADARs, respectively), editing of both tRNA and mRNA depends on a single enzyme in bacteria, the tRNA deaminase TadA (21). Consistent with this, mRNA editing sites in bacteria resemble the sequence and structure of the anticodon loop of the *bona fide* TadA target, tRNA^Arg^_ACG_ (21). In *E. coli*, most editing sites were located in coding sequences and resulted in recoding of tyrosine to cysteine. By studying the toxin-encoding *hokB* mRNA in more detail, Bar-Yaacov *et al.* were able to show that *hokB* editing was growth phase-dependent and resulted in a more toxic recoded protein isoform (21, 22). In *Klebsiella pneumoniae*, recoding of a transcriptional regulator was similarly growth phase-dependent and affected quorum sensing and virulence (23). A study carried out in the plant pathogen *Xanthomonas oryzae* demonstrated that editing of an mRNA encoding a ferric siderophore receptor increased under iron deprivation, leading to enhanced iron uptake and an improved chemotactic response towards iron (24).

Despite the emerging role of a dynamic, stress-responsive editome in bacterial physiology and adaptation, the underlying regulatory mechanisms remain elusive. It is known that eukaryotic ADAR-mediated editing is regulated by a multi-faceted network involving ADAR expression, post-translational modifications, sub-cellular localisation and additional *trans*-acting factors interfering with ADAR activity or target RNA accessibility (reviewed in (25, 26)). Indeed, changes in TadA expression and activity as well as the competition between tRNA and mRNA target sites have been proposed to control editing levels in bacteria (22). However, none of these hypotheses have been tested experimentally to date, and we still lack a comprehensive understanding of the occurrence, plasticity and regulation of bacterial A-to-I editing.

Here, we have characterised A-to-I editing in the strictly human pathogen *S. pyogenes*, which causes a wide range of diseases ranging from mild superficial to severe invasive infections. We demonstrate that the expansion of A34-to-I34 editing to a second tRNA substrate, tRNA^Leu^_AAG_, also leads to more diverse mRNA editing. By studying in more detail the response to several editing-modulating stress conditions, we provide insight into the mechanisms that contribute to the dynamics of the editome in *S. pyogenes*, and identify mRNA stability as an important determinant of bacterial A-to-I editing.

## MATERIAL AND METHODS

### Bacterial culture

*S. pyogenes* and isogenic derivatives (Table S1) were cultured without agitation at 37°C and 5% CO_2_. Trypticase soy agar (TSA, BD Difco) plates supplemented with 3% defibrinated sheep blood (Oxoid) were routinely used for growth on solid medium. C medium (0.5% (w/v) Protease Peptone No. 3 (Difco), 1.5% (w/v) yeast extract (Servabacter) and 1 g/L NaCl), THY medium (Bacto Todd Hewitt Broth (Becton Dickinson) with 0.2% yeast extract) or chemically defined medium (CDM) were used as liquid media. In brief, CDM was prepared from powder (order 4393306, Alpha Biosciences) and supplemented to 1 mg/L Fe(NO_3_)_3_ · 9 H_2_O, 5 mg/L FeSO_4_ · 7 H_2_O, 3.4 mg/L MnSO_4_ · H_2_O and 1% glucose. 50x stocks of NaHCO_3_ and L-cysteine in H_2_O were added to final concentrations of 2.5 g/L and 708 mg/L, respectively, freshly before inoculation. *E. coli* was grown at 37°C with shaking in Luria–Bertani medium or on agar. When needed, antibiotics were supplemented at the following concentrations: 100 μg/ml carbenicillin, 50 μg/ml kanamycin and 20 µg/mL chloramphenicol for *E. coli*; 300 μg/ml kanamycin, 3 µg/mL erythromycin and 20 µg/mL chloramphenicol for *S. pyogenes*. Overnight (O/N) cultures of *S. pyogenes* were used to inoculate cultures at an optical density at 620 nm (OD_620_) of 0.02, and cell growth was monitored using a microplate reader (200 µL sample volume; BioTek Epoch 2 Microplate Spectrophotometer).

### Bacterial transformation

*S. pyogenes* competent cells for the integration of DNA into the genome were generated as previously described (27). Competent cells were adjusted to an OD_620_ of 2.0 in 100 µL and incubated with 10 to 15 µg of DNA on ice for 1 h before electroporation in a 0.1 cm electrode gap cuvette (1.5 kV, 400 Ω and 25 μF pulse; BioRad Gene Pulser Xcell electroporation system). After recovery in THY, mutants were selected on plates with the respective antibiotic. Electrocompetent cells for transformation of *S. pyogenes* with plasmids were prepared according to a previously described protocol (28) and transformed as above except that only 500 ng of plasmid DNA and a pulse of 1.8 kV, 400 Ω and 25 μF were used. *E. coli* strains used for cloning were transformed according to the standard heat shock procedure (29).

### DNA handling and cloning

Standard molecular biology techniques, such as PCR (Phusion High-Fidelity DNA Polymerase and recombinant Taq DNA Polymerase, Thermo Scientific), DNA digestion with restriction enzymes (Thermo Scientific), DNA ligation (T4 DNA ligase, Thermo Scientific), 5’ end phosphorylation (T4 polynucleotide kinase, Thermo Scientific), purification of PCR products (DNA Clean & Concentrator-5, Zymo Research), gel extraction (QIAquick Gel Extraction Kit), plasmid DNA preparation (QIAprep Spin MiniPrep Kit and Plasmid Midi Kit, Qiagen) and gel electrophoresis were performed according to the manufacturer’s instructions or using standard protocols (29). Site-directed mutagenesis of plasmids was performed following the two-stage PCR-based protocol (30). All oligonucleotides (Sigma-Aldrich) used in this study are listed in Table S2. All constructed plasmids were validated by Sanger sequencing (Microsynth Seqlab) and are listed in Table S3. Cloning procedures are detailed in Supplementary Methods, and schematic views of mutant loci are presented in Figure S1.

### Analysis of RNA editing and abundance

#### Isolation of total RNA

Samples of *S. pyogenes* were collected by mixing the bacterial culture with an equal volume of ice-cold acetone/ethanol (1:1, v/v). Cells were pelleted, washed and pre-treated with lysozyme and mutanolysin, before RNA was extracted using TRIzol (Invitrogen) / chloroform and precipitated with isopropanol. Total RNA was treated with TURBO DNase (Invitrogen) and recovered using RNA Clean & Concentrator-5 (Zymo Research).

#### Detection of editing by Sanger sequencing

500 ng of total RNA or 20 ng of *in vitro* transcribed RNA were reverse transcribed with random hexamer primers (Thermo Scientific) or tRNA-specific primers using SuperScript III Reverse Transcriptase (Invitrogen). 1 µL of the reverse transcription reaction was used to amplify regions of interest using gene-specific primers in a 50 µL PCR reaction using recombinant *Taq* DNA Polymerase (Invitrogen) (see Table S2). Amplicons were purified using DNA Clean & Concentrator-5 (Zymo Research) and analysed by Sanger sequencing (Microsynth Seqlab). Sequencing chromatograms were analysed using 4Peaks and SnapGene Viewer. Raw peak values were used to quantify editing levels.

#### Quantitative RT-PCR

qRT-PCR was performed using the Power SYBR Green RNA-to-CT 1-Step Kit (Applied Biosystems) in a QuantStudio 5 Real-Time PCR System (Applied Biosystems) using 10 ng of DNase-treated total RNA per 20 µL reaction. *gyrA*, *rpoB*, *era* and *secA* served as reference genes as indicated, and relative expression was calculated as previously described (31, 32). Primers used for qRT-PCR are listed in Table S2.

### Detection of inosine by RNase T1 digestion of glyoxal/borate-protected RNA

RNase T1 digestion of glyoxal/borate-protected RNA was performed as previously described (33, 34). 10 µg of total RNA were incubated in 10 mM sodium phosphate buffer (pH 7.0) with 50% DMSO and 2.4% deionised glyoxal (Sigma-Aldrich) for 45 min at 37°C and subsequently mixed with an equal volume of 1 M sodium borate. RNA was then precipitated with ethanol and resuspended in Tris-borate buffer (1 M sodium borate, 10 mM Tris-HCl at pH 7.5). Samples were split, incubated for 20 min at 37°C with RNase T1 (Thermo Scientific) or H_2_O, and treated with proteinase K for 20 min at 37°C. RNA was then extracted using TRIzol / chloroform and precipitated using isopropanol. For de-glyoxalation, RNA was resuspended in 10 mM sodium phosphate buffer (pH 7.0) with 50% DMSO and incubated for 3 h at 65°C at 300 rpm with additional flicking every 30 min. RNA was precipitated with ethanol in the presence of RNA grade glycogen (Thermo Scientific). 100 ng of each sample were then subjected to polyacrylamide Northern blot analysis.

### Denaturing gel electrophoresis and Northern blotting

RNA was denatured for 5 min at 95°C in 1x RNA Loading Dye (NEB) and separated on denaturing polyacrylamide gels (12%, 19:1 acrylamide/bisacrylamide, 8 M urea, 1x TBE) or agarose gels (1%, 20 mM MOPS (pH 7.0), 5 mM NaOAc, 1 mM EDTA, 6.6% formaldehyde). For direct visualisation of RNA, post-staining with SYBR Gold Nucleic Acid Gel Stain (Invitrogen) was performed. Alternatively, Northern blotting was performed as previously described with minor modifications (35, 36). RNA was transferred from gels onto Amersham Hybond-N+ membranes (Cytiva) by electroblotting or capillary transfer (in 20x SSC), respectively. After UV cross-linking, membranes were pre-hybridised in Rapid-hyb Buffer (Cytiva). Oligonucleotide probes were radiolabelled with γ-^32^P-ATP (Hartmann Analytic) using T4 polynucleotide kinase (Thermo Scientific), purified using illustra MicroSpin G-25 columns (Cytiva) and added to the pre-hybridisation buffer. Hybridisation was conducted O/N at 42°C under constant rotation. Membranes were washed twice and exposed to storage phosphor screens. Signals were visualised with a Typhoon FLA-9000 laser scanner. As a loading control, membranes were probed against 5S or 16S rRNA. Decade Markers System (Invitrogen) and RiboRuler High Range RNA Ladder (Thermo Scientific) were used as size marker.

### *In vitro* tRNA deamination assay

#### In vitro transcription

T7 promoter-containing plasmid DNA was digested with FastDigest MvaI (Thermo Scientific) and used as template for *in vitro* transcription of tRNA using the AmpliScribe T7-Flash Transcription Kit for 4 h at 42°C (see Table S3). After DNase treatment, RNA was separated on denaturing polyacrylamide gels, visualised by UV shadowing, excised and purified using the ‘crush-and-soak’ method (29). RNA purity was verified by gel electrophoresis and quantified using a NanoDrop spectrophotometer.

#### TadA protein purification

NiCo21(DE3) Competent *E. coli* (NEB) was transformed with overexpression plasmids encoding *E. coli* or *S. pyogenes tadA* fused to a C-terminal His-tag (see Table S3). Cells were grown to an OD_600_ of 0.8, and expression was induced with 0.4 mM IPTG. After growth for 4 h at 37°C, cells were pelleted, resuspended in lysis buffer (50 mM Tris-HCl (pH 8.0), 500 mM NaCl, 5 mM β-mercaptoethanol, 5% glycerol, 0.1% TWEEN 20, 10 µM ZnCl_2_) and lysed by sonication (6 × 30 s with 30 s on ice in between; SONOPULS HD4050 ultrasonic homogenizer). Cleared lysates were incubated with Ni-NTA Agarose (Qiagen) at 4°C for 1 h. After two washes with lysis buffer containing 20 mM imidazole, recombinant TadA was eluted with elution buffer (lysis buffer with 250 mM imidazole) and elution fractions were analysed by SDS-PAGE. Protein-containing fractions were pooled and incubated with Chitin Resin (NEB) for 30 min at 4°C. The flow-through was dialysed O/N at 4°C in a SnakeSkin™ Dialysis Tubing (3.5K MWCO, Thermo Scientific) against dialysis buffer (50 mM Tris-HCl (pH 8.0), 100 mM NaCl, 2 mM DTT, 5% glycerol, 10 µM ZnCl_2_). Protein concentration was determined using the Bio-Rad Protein Assay. Samples were mixed with an equal amount of 80% glycerol, flash-frozen in liquid nitrogen and stored at −80°C.

#### In vitro tRNA deamination

*In vitro* deamination was performed as described before (13). To refold tRNA, samples in 0.5x TE buffer (pH 8.0) were denatured for 5 min at 80°C and cooled to 60°C. After incubation for 5 min, MgCl_2_ was added to 10 mM and samples were cooled to 20°C at 0.1°C/s. 1 µg of tRNA was then mixed at an equimolar ratio with recombinant TadA in deamination buffer (50 mM Tris-HCl (pH 8.0), 25 mM NaCl, 2.5 mM MgCl_2_, 0.1 mM EDTA, 10% glycerol, 2 mM DTT, 20 µg/mL BSA). Samples were incubated for 30 min at 37°C and purified using the RNA Clean & Concentrator-5 kit (Zymo Research).

### Approaches for studying the dynamics and regulation of editing

#### Dynamics of editing in response to different culture conditions

Cultures were grown to mid-logarithmic growth phase at 37°C and 5% CO_2_ in C medium. For temperature shift experiments, cultures were diluted 1:10 into pre-conditioned C medium and incubated for 15 min at the respective temperatures (12°C, 30°C, 37°C, and 42°C; without CO_2_). To apply zinc or oxidative stress, cultures were split and compounds were added at the desired concentration (ZnSO_4_ at 500 µM, and H_2_O_2_ at up to 2 mM) with mock-treated cultures as controls. Cultures were incubated at 37°C and 5% CO_2_ for the indicated time. To study the effect of different sugars, cells were pelleted, resuspended in fresh, pre-warmed medium with sugars supplemented at 0.5% (w/v) and incubated for 30 min. Culture aliquots were harvested before splitting (t = 0 min) and after the indicated time for downstream applications.

#### Effect of tadA expression during stress

Strains SF370 P_tet_-TT*_tadA_* and P_tet_-*tadA* (EC3570 and EC3622, see Table S1) were grown O/N without AHT and used to inoculate cultures in C medium. At mid-logarithmic growth phase, cultures were split and exposed for 30 min to 1 mM H_2_O_2_, 0.5 mM ZnSO_4_ or 100 ng/mL AHT with H_2_O as mock-treated control. Culture aliquots were removed and total RNA was extracted. Editing levels of *pepN* and *fakB2* were analysed by Sanger sequencing, and expression of *tadA* was examined by qRT-PCR with *gyrA, rpoB* and *era* as reference genes. Samples were normalised to the mean editing and expression levels of the untreated control per strain.

#### Effect of gene expression levels

Strains SF370 P_tet_-*slo*, P_tet_-*amyA* and P_tet_-*fakB2* (EC3453, EC3456 and EC3459, see Table S1), grown O/N in C medium without AHT, were used to inoculate cultures in C medium with increasing concentrations of AHT (0, 0.1, 1, 10 and 100 ng/mL) and grown to mid-logarithmic growth phase. Editing levels were examined by Sanger sequencing and gene expression by qRT-PCR for *amyA* and *fakB2*. Samples from strain SF370 P_tet_-*slo* served as controls for native expression and editing of both genes. Relative gene expression was calculated with *gyrA* and *rpoB* as reference genes and normalised to the mean expression at 0 ng/mL AHT.

#### ermBL-based luminescence reporter assay

Strain SF370 was transformed with empty vector control pEC2812, the wildtype reporter pEC3045, and the reporter mutants pEC3046, pEC3047 and pEC3069. Cultures were grown to mid-logarithmic growth phase in C medium in the presence of kanamycin and exposed for 30 min to different concentrations of erythromycin. Culture aliquots (200 µL) were taken to measure OD_620_ and luminescence by adding 10 µL of 1 mg/mL beetle luciferin (Promega) in a BioTek Cytation 3 Microplate Reader, and samples were withdrawn for RNA extraction. Empty vector-corrected luminescence was normalised by OD_620_, and editing levels and *ffluc* expression were examined by Sanger sequencing and qRT-PCR with *gyrA* and *rpoB* as reference genes.

#### Analysis of RNA half-lives

Strain SF370 was grown to mid-logarithmic growth phase and exposed to different stimuli for 30 min (mock-treated, 1 mM H_2_O_2_, 0.5 mM ZnSO_4_ or 0.5% (w/v) glucose). Rifampicin (Sigma-Aldrich) was added to a final concentration of 250 µg/mL, and samples were collected right before rifampicin addition (t = 0 min) and after 1, 2, 4, 8, 16 and 32 min. mRNA abundances were examined by qRT-PCR with 16S rRNA as reference, and half-lives were calculated according to the “steepest slope” method using four consecutive timepoints (37). Editing levels at t = 0 min were analysed by Sanger sequencing. Strain SF370 P_tet_-*rnjA* and P_tet_-*tadA* were processed similarly but grown to mid-logarithmic growth phase in the presence or absence of 0.1 ng/mL or 100 ng/mL AHT, respectively, before adding rifampicin.

### Next Generation Sequencing

#### Culture conditions for NGS experiments

Cultures of *S. pyogenes* SF370 were grown at 37°C and 5% CO_2_ in C medium, and culture aliquots were removed for DNA and RNA extraction at early logarithmic, mid-logarithmic and early stationary growth phase in triplicates. In case of strains 5448 and 5448AP, experiments were performed in duplicate. To analyse the effect of *tadA* overexpression, *S. pyogenes* SF370 was transformed with the empty control vector pEC2812 and *tadA*-expressing pEC2813, and grown to mid-logarithmic growth phase in triplicate. To compare editing in different culture media, O/N cultures of *S. pyogenes* SF370 in C medium were washed in phosphate-buffered saline and used to inoculate cultures of C medium, THY and CDM, which were then grown to mid-logarithmic phase in triplicate. To study the response to H_2_O_2_, *S. pyogenes* SF370 was grown to mid-logarithmic growth phase and exposed to 0.5 mM or 1.0 mM H_2_O_2_ for 15 min and 30 min with mock-treated culture as control. Lastly, *S. pyogenes* SF370 P_tet_-*rnjA* was grown to mid-logarithmic growth phase in THY in the presence or absence of 0.1 ng/mL AHT.

#### Whole-genome sequencing

Genomic DNA was prepared with the NucleoSpin Microbial DNA Kit (Macherey-Nagel) and treated with 1 µL of RNase Cocktail™ Enzyme Mix (Invitrogen). Genomic DNA libraries were prepared and sequenced by the Sequencing Core Facility of the Max Planck Institute for Molecular Genetics (Berlin, Germany) using the Nextera XT DNA Library Preparation Kit or the KAPA HyperPrep DNA Kit.

#### RNA sequencing

Ribosomal RNA was depleted from TURBO DNase-treated total RNA using the MICROBExpress Bacterial mRNA Enrichment Kit (Invitrogen), Pan-Prokaryote riboPOOL (siTOOLs Biotech) or Ribo-Zero rRNA Removal Kit for Bacteria (Illumina). rRNA-depleted RNA was treated with RppH (NEB), purified using phenol extraction, precipitated with ethanol and dissolved in H_2_O. RNA samples were then subjected to T4 PNK (NEB) treatment and purified using RNA Clean & Concentrator-5 (Zymo Research). In case of SF370 P_tet_-*rnjA*, only the large RNA fraction (> 200 nt) was purified and further processed. Next, RNA was sheared in a microTUBE AFA Fiber Pre-Slit Snap-Cap for 140 s in a Covaris M220 Focused-ultrasonicator. Alternatively, samples were T4 PNK-treated after sonication. Afterwards, cDNA libraries were prepared using the NEXTflex Small RNA-Seq Kit v3 (Perkin Elmer) according to the manufacturer’s protocol. Library quality was analysed using the Agilent High Sensitivity DNA Kit and the Qubit dsDNA HS Assay Kit. Sequencing was performed at the Sequencing Core Facility of the Max Planck Institute for Molecular Genetics (Berlin, Germany).

### NGS data analysis and conservation of editing site

#### Identification of A-to-I editing events in NGS datasets

A detailed description of the bioinformatic analysis is provided in Supplementary Methods. In brief, adapter sequences were removed using Cutadapt (v2.10) (38) and reads were mapped using BWA-MEM (v.0.7.17) (39). After PCR de-duplication using UMI Tools (v1.0.1) and read merging, single nucleotide polymorphisms were identified according to the following criteria: coverage ≥ 20x, quality score ≥ 30, minimum distance of 4 nt from read ends, ≥ 2 supporting reads per direction and a frequency ≥ 0.01. SNPs present in RNA but not in matched DNA datasets were further filtered for A-to-G transitions, and the identified sites were manually curated and reported with their respective modification level. For visualisation of changes in editing levels, editing z scores were calculated for each target site with enough coverage support in at least two out of three replicates and a mean editing level of at least 3%. Publicly available RNA-seq datasets from other strains of *S. pyogenes* (40–42) were processed as described but with whole-genome sequencing data simulated using Mason (v0.1.2; 1 million reads in paired-end mode, 150 bp) (43) due to the lack of matched experimental data. Editing sites with a frequency of at least 0.01 were then compared to sites experimentally identified in *S. pyogenes* SF370.

#### Sequence and structure consensus of A-to-I editing sites

The local sequence context consensus of the identified mRNA editing sites (± 5 nt, tRNA genes not considered) was generated using Weblogo (44) with position 0 representing the editing position. Secondary structure predictions for RNA were performed using RNAfold from the ViennaRNA Package 2.0 (45). The minimum free energy (MFE) was calculated using a sliding window approach around each target site with a window size of 17 nt (length of tRNA anticodon arm), and MFE average and standard deviation were calculated over all target sites. Control positions within coding sequences of the *S. pyogenes* SF370 genome were retrieved using the position-specific scoring matrix-based approach of the “motifs” package of the Python library Biopython (for positions −2 to +4 relative to editing position), and the sliding window approach was applied to the control position set as before.

#### Differential expression analysis

Reads were filtered for a minimum quality score of 10 and a length of at least 22 nt, trimmed using Cutadapt (v1.11) (38) and mapped against the reference genome NC_002737.2 using STAR (v2.7.3a) (46) in ‘random best’ and ‘end-to-end’ modes. BAM files were sorted and indexed using Samtools (v1.9) (47), and PCR de-duplication was performed using UMI Tools (v0.4.1). Gene counts were determined with featureCounts (v2.0.0) (48) and differentially expressed genes were identified using DESeq2 (v1.26.0) (49).

#### Phylogenetic analysis of editing site conservation in Streptococcaceae species

Taxonomy tree and genome assemblies for all *Streptococcaceae* species were retrieved from NCBI and filtered for the most recent and complete assembly versions of each available strain (as of 2019-12-08). For each editing site, coding and amino acid sequence of the parent gene was extracted from the *S. pyogenes* SF370 reference genome (NC_002737.2). All annotated CDS sequences of selected *Streptococcaceae* genomes were extracted, and homolog candidates for each edited gene in *S. pyogenes* SF370 were identified based on the similarity of the amino acid reference sequence. Homolog candidate genes as well as similar non-related sequences were included in a multisequence alignment calculated with ClustalO (v1.2.4). After a first round of scoring (pairwise Hamming distance via Python library Biopython) to establish a threshold in a multimodal model, false-positive hits were removed. Multi-sequence alignment using ClustalO was repeated using the identified homologs, and the amino acid and codon identity of the homologs at the editing position of the reference gene was reported.

## RESULTS

### TadA from *S. pyogenes* modifies two tRNAs

The genome of *S. pyogenes* encodes a second A34-tRNA other than the canonical tRNA^Arg^_ACG_. To explore the potential expansion of A-to-I editing to this additional tRNA^Leu^_AAG_, we examined the expression and modification status of both A34-tRNAs, exploiting the inosine-specific cleavage activity of RNase T1 (33). In the absence of RNase T1, full-length tRNA^Arg^_ACG_ and tRNA^Leu^_AAG_ could be detected by Northern blotting, confirming the expression of both A34-tRNAs *in vivo* (Figure 1A). Treatment with RNase T1 resulted in full cleavage of both tRNA^Arg^_ACG_ and tRNA^Leu^_AAG_, suggesting that both tRNAs are fully edited to I34 during all growth phases, whereas a control tRNA containing U34 was not cleaved by RNase T1 (Figure 1A). Since we observed an additional cleavage fragment for the non-canonical tRNA^Leu^_AAG_, we also performed Sanger sequencing, exploiting the interpretation of inosine as guanosine by reverse transcriptases, and observed complete modification of A34 to I34 in both tRNAs as well (Figure 1B). To investigate whether the tRNA deaminase TadA from *S. pyogenes* has adapted to accept a second tRNA substrate, we performed *in vitro* deamination reactions using recombinant TadA from *E. coli* and *S. pyogenes* and the *in vitro* transcribed TadA target tRNAs from both species. TadA from both *E. coli* and *S. pyogenes* efficiently modified the canonical tRNA^Arg^_ACG_ from either species (Figure 1C). In contrast, only *S. pyogenes* TadA modified the *S. pyogenes*-specific, non-canonical tRNA^Leu^_AAG_ (Figure 1C). Our findings thus demonstrate the expansion of A34-to-I34 editing in *S. pyogenes* and suggest that the streptococcal TadA enzyme has evolved to accept an extended substrate range.

**Figure 1.**
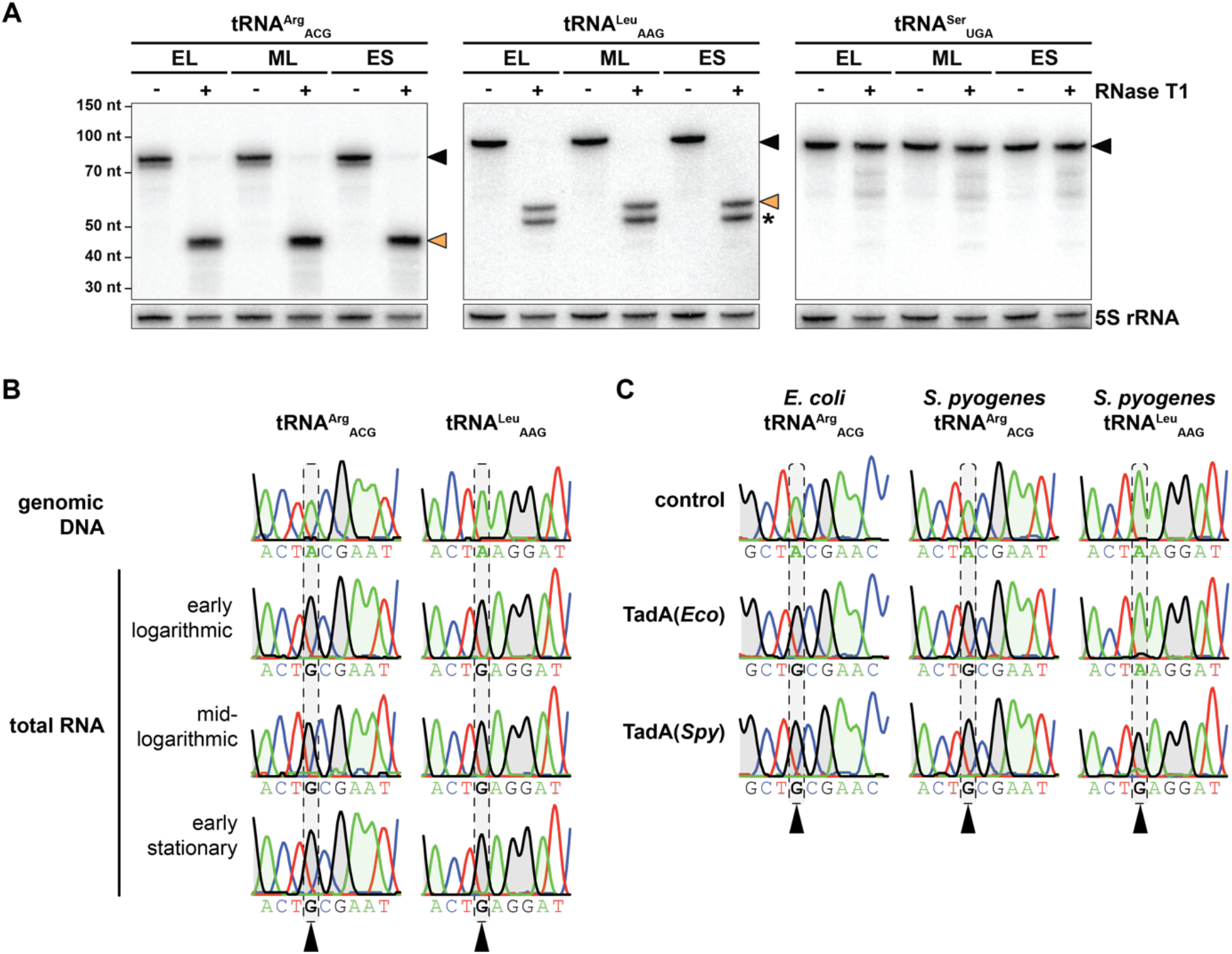
A34-to-I34 editing of *S. pyogenes* tRNAs *in vivo* and *in vitro*. (A) Expression and editing status of the two A34-tRNAs of *S. pyogenes* and of a non-A34 control tRNA were analysed by RNase T1 treatment of glyoxal/borate-protected total RNA and subsequent Northern blotting against tRNA 3’ ends. Full-length tRNAs are marked with a black arrow and inosine-dependent cleavage products with an orange arrow. Expected sizes are 77 nt *vs*. 42 nt for tRNA^Arg^_ACG_, 89 nt *vs*. 54 nt for tRNA^Leu^_AAG_, and 93 nt for the negative control tRNA^Ser^_UGA_ (no I34 modification). An additional cleavage fragment for tRNA^Leu^_AAG_, likely due to the presence of another tRNA modification that inhibits the formation of an RNase T1-resistant guanosine adduct, is marked by an asterisk. 5S rRNA was used as loading control. EL: early logarithmic; ML: mid-logarithmic; ES: early stationary. (B) The editing status of the two A34-tRNAs of *S. pyogenes* was further examined by Sanger sequencing at three different growth phases (three lower panels) and compared with the genomic DNA control (upper panel). (C) *In vitro* transcribed A34-tRNAs from *E. coli* (tRNA^Arg^_ACG_ only) and *S. pyogenes* (tRNA^Arg^_ACG_ and tRNA^Leu^_AAG_) were deaminated *in vitro* with recombinant TadA from both species or with buffer only as control. The Sanger chromatograms in (B) and (C) correspond to tRNA positions 31 to 39 with the editing position 34 highlighted by grey dashed boxes. Nucleotide sequences are shown below each chromatogram. Sequencing traces of adenosine and guanosine are shaded in green and grey, respectively, for better visualisation.

### mRNA editing events are more diverse in *S. pyogenes* than in *E. coli*

Given the expanded tRNA editing capacity of *S. pyogenes* TadA, we investigated the potential consequences for A-to-I editing at the transcriptome scale. To this end, we sequenced total RNA and genomic DNA from *S. pyogenes* SF370 grown in C medium at three different growth phases, and searched for A-to-I editing sites as A-to-G transitions present in RNA sequences but absent in genomic DNA sequences (see Material and Methods). Our transcriptome-wide analysis identified 27 editing events beyond the known positions in tRNA^Arg^_ACG_ and tRNA^Leu^_AAG_ (Table 1 and Table S4). Interestingly, all editing positions were located in protein coding sequences. Editing levels at several target sites reached 30% to 40%, although most sites were edited at less than 20%. Among the 18 sites predicted to recode the affected gene, three different types of recoding events were observed: (i) Lys-to-Glu (<u>A</u>AG codons, 10/18 sites), (ii) Tyr-to-Cys (U<u>A</u>U and U<u>A</u>C codons, 5/18 sites), and (iii) Thr-to-Ala recoding (<u>A</u>CG codons, 3/18 sites). The remaining 9 editing events were predicted to be synonymous and exclusively found for Leu codons (UU<u>A</u> and CU<u>A</u> codons, 9/9 sites) (Figure 2A). The identified A-to-I editing target genes were involved in various cellular processes, such as metabolic pathways (*e.g.*, *hpt, pepN*, *aroK*, and *amyA*), transport processes (*e.g.*, *adcC* and *opuC*), DNA integrity (*e.g.*, *ftsK* and *dinP*), tRNA biogenesis (*e.g.*, *trhP1* and *proS*) and virulence (*speC* and *speI*) (Table 1). We validated several A-to-I editing events using Sanger sequencing (Figure S2) and confirmed TadA-dependent mRNA editing by ectopic overexpression of *tadA* (Figure S3 and Table S5). While *tadA* is essential in *E. coli* (13), a deletion mutant in *Bacillus subtilis* is viable, albeit at compromised growth and translation fidelity (50). However, we were unable to obtain a clean *tadA* deletion strain in *S. pyogenes* using a Cre-lox-based approach (data not shown) (27). Although other deaminases may contribute to A-to-I editing, the overall increase in editing levels in response to *tadA* overexpression suggests that TadA is the primary mRNA editing enzyme in *S. pyogenes* (Figure S3).

**Table 1.**
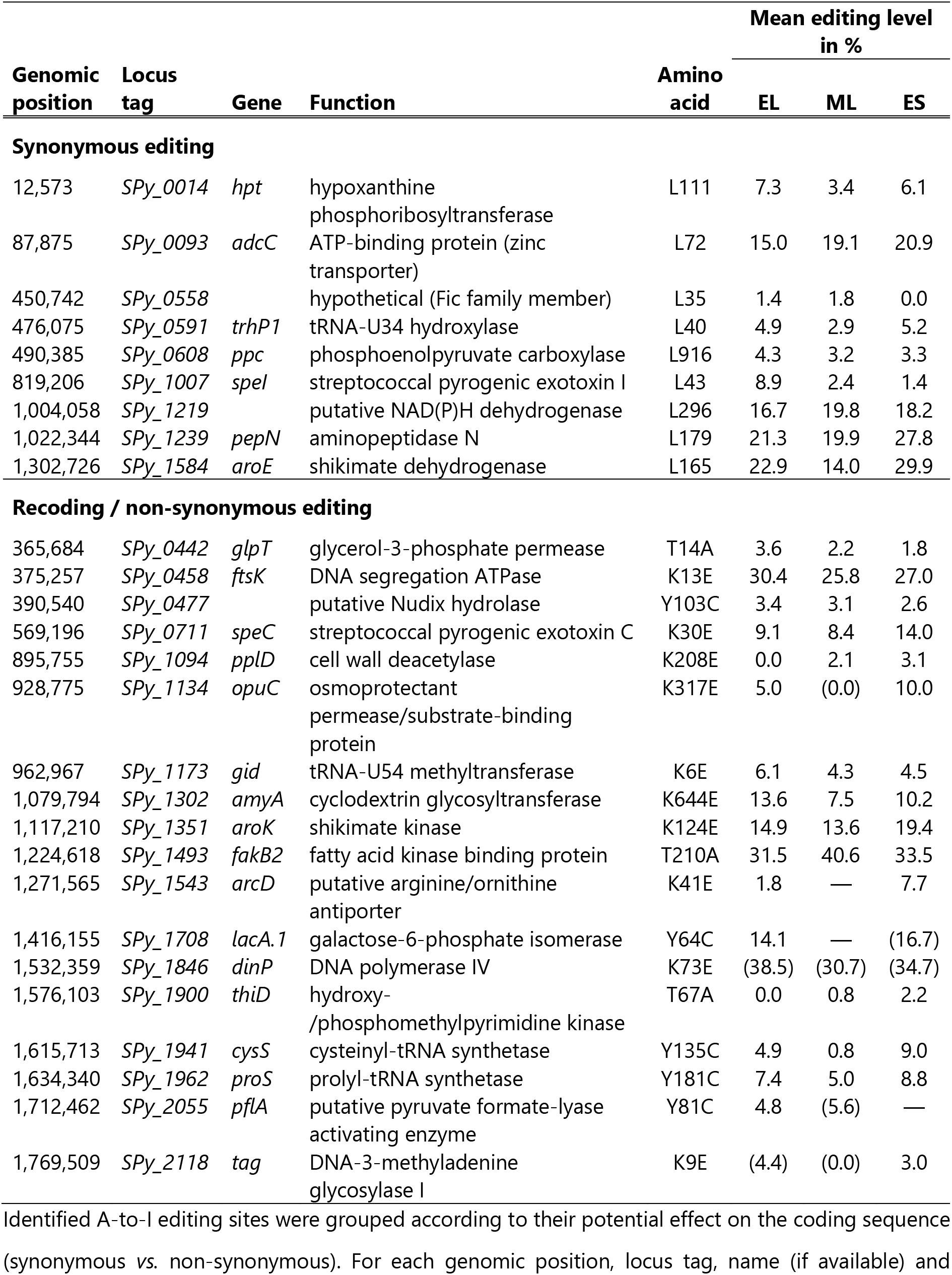
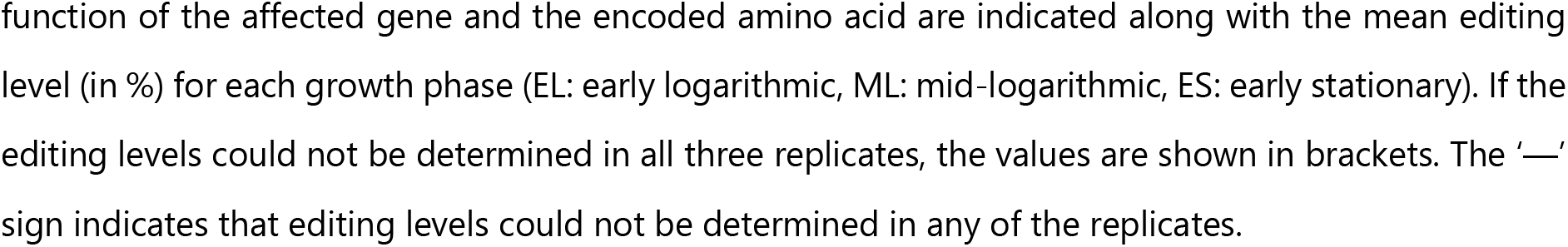
A-to-I editing in the transcriptome of *S. pyogenes* SF370.

| Genomic position | Locus tag | Gene | Function | Amino acid | Mean editing level in % |  |  |
| --- | --- | --- | --- | --- | --- | --- | --- |
|  |  |  |  |  | EL | ML | ES |
| Synonymous editing |  |  |  |  |  |  |  |
| 12,573 | SPy_0014 | *hpt* | hypoxanthine phosphoribosyltransferase | L111 | 7.3 | 3.4 | 6.1 |
| 87,875 | SPy_0093 | *adcC* | ATP-binding protein (zinc transporter) | L72 | 15.0 | 19.1 | 20.9 |
| 450,742 | SPy_0558 |  | hypothetical (Fic family member) | L35 | 1.4 | 1.8 | 0.0 |
| 476,075 | SPy_0591 | *trhP1* | tRNA-U34 hydroxylase | L40 | 4.9 | 2.9 | 5.2 |
| 490,385 | SPy_0608 | *ppc* | phosphoenolpyruvate carboxylase | L916 | 4.3 | 3.2 | 3.3 |
| 819,206 | SPy_1007 | *speI* | streptococcal pyrogenic exotoxin I | L43 | 8.9 | 2.4 | 1.4 |
| 1,004,058 | SPy_1219 |  | putative NAD(P)H dehydrogenase | L296 | 16.7 | 19.8 | 18.2 |
| 1,022,344 | SPy_1239 | *pepN* | aminopeptidase N | L179 | 21.3 | 19.9 | 27.8 |
| 1,302,726 | SPy_1584 | *aroE* | shikimate dehydrogenase | L165 | 22.9 | 14.0 | 29.9 |
| Recoding / non-synonymous editing |  |  |  |  |  |  |  |
| 365,684 | SPy_0442 | *glpT* | glycerol-3-phosphate permease | T14A | 3.6 | 2.2 | 1.8 |
| 375,257 | SPy_0458 | *ftsK* | DNA segregation ATPase | K13E | 30.4 | 25.8 | 27.0 |
| 390,540 | SPy_0477 |  | putative Nudix hydrolase | Y103C | 3.4 | 3.1 | 2.6 |
| 569,196 | SPy_0711 | *speC* | streptococcal pyrogenic exotoxin C | K30E | 9.1 | 8.4 | 14.0 |
| 895,755 | SPy_1094 | *pplD* | cell wall deacetylase | K208E | 0.0 | 2.1 | 3.1 |
| 928,775 | SPy_1134 | *opuC* | osmoprotectant permease/substrate-binding protein | K317E | 5.0 | (0.0) | 10.0 |
| 962,967 | SPy_1173 | *gid* | tRNA-U54 methyltransferase | K6E | 6.1 | 4.3 | 4.5 |
| 1,079,794 | SPy_1302 | *amyA* | cyclodextrin glycosyltransferase | K644E | 13.6 | 7.5 | 10.2 |
| 1,117,210 | SPy_1351 | *aroK* | shikimate kinase | K124E | 14.9 | 13.6 | 19.4 |
| 1,224,618 | SPy_1493 | *fakB2* | fatty acid kinase binding protein | T210A | 31.5 | 40.6 | 33.5 |
| 1,271,565 | SPy_1543 | *arcD* | putative arginine/ornithine antiporter | K41E | 1.8 | — | 7.7 |
| 1,416,155 | SPy_1708 | *lacA.1* | galactose-6-phosphate isomerase | Y64C | 14.1 | — | (16.7) |
| 1,532,359 | SPy_1846 | *dinP* | DNA polymerase IV | K73E | (38.5) | (30.7) | (34.7) |
| 1,576,103 | SPy_1900 | *thiD* | hydroxy-/phosphomethylpyrimidine kinase | T67A | 0.0 | 0.8 | 2.2 |
| 1,615,713 | SPy_1941 | *cysS* | cysteinyl-tRNA synthetase | Y135C | 4.9 | 0.8 | 9.0 |
| 1,634,340 | SPy_1962 | *proS* | prolyl-tRNA synthetase | Y181C | 7.4 | 5.0 | 8.8 |
| 1,712,462 | SPy_2055 | *pflA* | putative pyruvate formate-lyase activating enzyme | Y81C | 4.8 | (5.6) | — |
| 1,769,509 | SPy_2118 | *tag* | DNA-3-methyladenine glycosylase I | K9E | (4.4) | (0.0) | 3.0 |

Identified A-to-I editing sites were grouped according to their potential effect on the coding sequence (synonymous vs. non-synonymous). For each genomic position, locus tag, name (if available) and
function of the affected gene and the encoded amino acid are indicated along with the mean editing level (in %) for each growth phase (EL: early logarithmic, ML: mid-logarithmic, ES: early stationary). If the editing levels could not be determined in all three replicates, the values are shown in brackets. The '—' sign indicates that editing levels could not be determined in any of the replicates.

**Figure 2.**
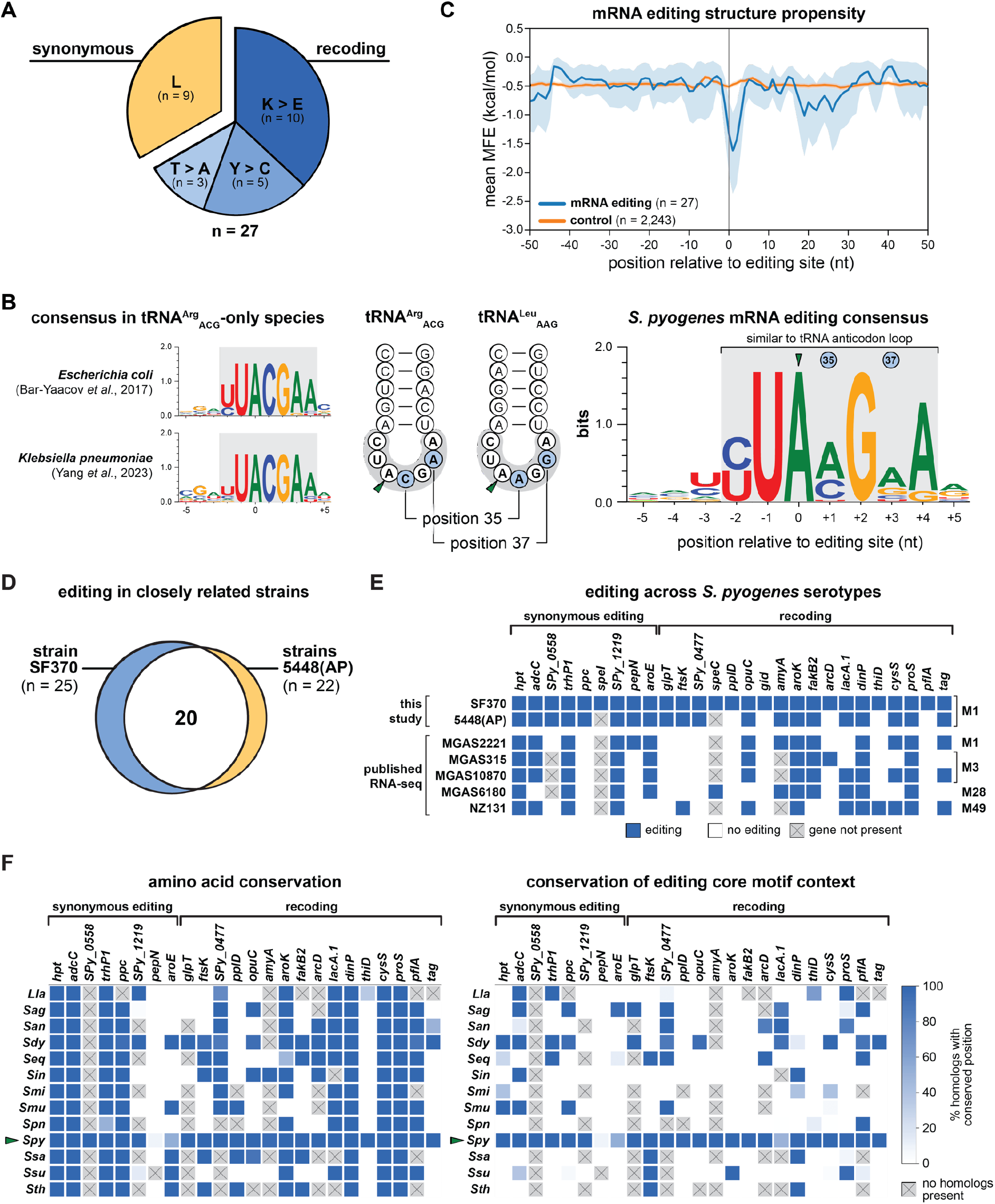
mRNA editing in *S. pyogenes*. (A) The A-to-I editing events identified in *S. pyogenes* SF370 (n = 27) were classified as synonymous or non-synonymous (*i.e.*, recoding), and further sorted according to the effect on the amino acid level. (B) Left: mRNA editing consensus motifs in *E. coli* and *K. pneumoniae*. Middle: Schematic representation of the tRNA anticodon arms of the two TadA substrate tRNAs in *S. pyogenes* with the loop sequence shaded in grey and diverging nucleotides at position 35 and 37 highlighted in blue. Right: Consensus of the sequence context of the editing sites identified in *S. pyogenes* (± 5 nt) with sequence conservation shown in bits. (C) Secondary structure propensity around A-to-I editing target sites in *S. pyogenes* SF370 (blue) and control positions in coding sequences harbouring the YUAMGRA consensus sequence (orange, n = 2, 243). (D) Venn diagram of editing target genes in *S. pyogenes* strain 5448 and hypervirulent strain 5448AP (referred to as 5448(AP)) compared with strain SF370. (E) Editing signatures in editing target genes in RNA-seq datasets from this study (upper rows) and publicly available RNAseq datasets (lower rows) of various *S. pyogenes* strains (names on the left) and M serotypes (indicated on the right). (F) Conservation of the amino acid (left) and TAMG core consensus context (right) in homologs of *S. pyogenes* SF370 editing target genes in selected *Streptococcaceae* family members. The fraction of homologs within each species (left) with identical amino acid and editable codons, respectively, is depicted for each editing target gene (top) as a heatmap. Synonymously edited and recoded target genes are grouped accordingly. *S. pyogenes* is highlighted by a green arrow. For homologs of *SPy_0477*, editing was also considered possible in the case of the TATG sequence, given the native editing context of the gene in *S. pyogenes* SF370. Species: *Lactococcus lactis* (*Lla*), *Streptococcus agalactiae* (*Sag*), *S. anginosus* (*San*), *S. dysgalactiae* (*Sdy*), *S. equi* (*Seq*), *S. iniae* (*Sin*), *S. mitis* (*Smi*), *S. mutans* (*Smu*), *S. pneumoniae* (*Spn*), *S. pyogenes* (*Spy*), *S. salivarius* (*Ssa*), *S. suis* (*Ssu*), and *S. thermophilus* (*Sth*).

In *E. coli* and *K. pneumoniae*, all editing events occur within the core consensus sequence U<u>A</u>CG and the extended motif YU<u>A</u>CGAA, resembling the anticodon loop sequence of the canonical TadA target tRNA tRNA^Arg^_ACG_ (position 32 to 38; Figure 2B, left) (21, 23). Consistent with the expanded range of *S. pyogenes* TadA tRNA substrates, we identified the more flexible core consensus UAMG (M = A or C) and the extended consensus motif YU<u>A</u>MGRA in *S. pyogenes* (Figure 2B, right). Importantly, lower nucleotide identity in the mRNA editing consensus was observed at positions corresponding to the divergent tRNA positions 35 and 37 of the TadA substrates tRNA^Arg^_ACG_ and tRNA^Leu^_AAG_. In addition to the consensus editing sequence, A-to-I editing sites in *E. coli* were found within tRNA anticodon arm-like secondary structures. To examine the secondary structure requirements in *S. pyogenes*, we used a sliding window approach to calculate the overall secondary structure propensity across all mRNA editing sites. As a control, we used positions within coding sequences harbouring the identified editing consensus sequence. Notably, we observed a significant decrease in the minimal free energy (MFE) around the editing site compared with the control (Figure 2C), indicating the presence of a secondary structure at the identified editing positions. In summary, A-to-I editing events in *S. pyogenes* mRNAs are more diverse than in species that only encode the canonical tRNA^Arg^_ACG_, and occur predominantly in tRNA anticodon arm-like structures with a core UAMG consensus sequence.

### RNA editing events are well-conserved across strains and serotypes of *S. pyogenes*

To gain insight into the conservation of the detected A-to-I editing events, we experimentally identified editing events in the closely related strain 5448 of the same M1 serotype. We also included the hypervirulent isogenic strain 5448AP, which harbours a mutation in the CovRS two-component system known to regulate approximately 15% of the *S. pyogenes* transcriptome (51). We identified a total of 22 editing positions in coding sequences of strains 5448(AP), 20 of which were previously detected in strain SF370 (Figure 2D and Table S6). When comparing editing levels among the three strains, we observed only few differences (Tables S4 and S6), most of which were due to high inter-replicate variability (*e.g.*, for *fakB2* or *dinP*). Similarly, the levels of editing in strains 5448 and 5448AP were highly concordant, with only a few sites showing considerable differences (*e.g.*, *pepN*). Thus, the CovRS system does not globally affect A-to-I editing levels.

We extended our analysis to other strains and M serotypes of *S. pyogenes* by analysing publicly available RNA-seq datasets for the presence of editing signatures at positions known to be modified in strain SF370. We detected six out of nine synonymous editing events and 13 out of 18 recoding events (Figure 2E). Notably, six target genes were edited in all datasets studied (*hpt*, *trhP1*, *SPy_1219*, *aroK*, *dinP*, and *proS*) and four additional genes in four out of five datasets (*adcC*, *aroE*, *opuC*, and *fakB2*). Therefore, a core set of A-to-I editing events is well-conserved across different serotypes and strains of *S. pyogenes*.

Lastly, we addressed the conservation of editing events within the *Streptococcaceae* family. We identified homologs of *S. pyogenes* SF370 editing target genes in selected *Streptococcaceae* species (Figure S4) and analysed the amino acid conservation at corresponding positions. In addition, we examined the presence of the core consensus sequence TAMG at the corresponding codon position to evaluate whether these sites are potentially amenable to A-to-I editing by TadA (hereafter termed ‘editability’). Consistent with our analysis of publicly available RNA-seq datasets, most editing target genes found in *S. pyogenes* SF370 exhibited high conservation of both amino acid identity and codon editability in other *S. pyogenes* strains (Figure 2F). In contrast, we observed only moderate conservation at the amino acid level and low conservation of codon editability across the *Streptococcaceae* species studied. In particular, the emerging pathogen *S. dysgalactiae*, which shares the highest gene content similarity with *S. pyogenes* (52), showed the highest conservation of codon editability among the *Streptococcaceae* species studied. In conclusion, A-to-I editing is generally highly conserved among different strains and serotypes of *S. pyogenes*, but is only poorly conserved in other *Streptococcaceae* species.

### mRNA editing is dynamic and responds to infection-relevant stimuli

Several mRNA modifications in eukaryotes, including inosine, are dynamically regulated in response to defined stimuli and conditions (53, 54). To gain insights into the dynamics of A-to-I editing in *S. pyogenes* mRNAs, we examined the effect of various environmental conditions and stresses. First, we compared A-to-I editing in three commonly used *S. pyogenes* culture media, namely chemically defined medium (CDM), THY, and C medium, but did not observe significant differences under standard growth conditions (Table S7).

During infection, *S. pyogenes* colonizes niches with varying temperature (*e.g.*, superficial skin infection *vs* invasive tissue infection), prompting us to examine the effect of different temperature shifts on A-to-I editing. Editing levels generally decreased with increasing temperatures and were highest for cultures exposed to a cold shock at 12°C (Figure 3A). Importantly, editing levels also changed in response to more infection-relevant temperature changes, with decreased editing at 42°C and increased editing at 30°C compared to 37°C.

**Figure 3.**
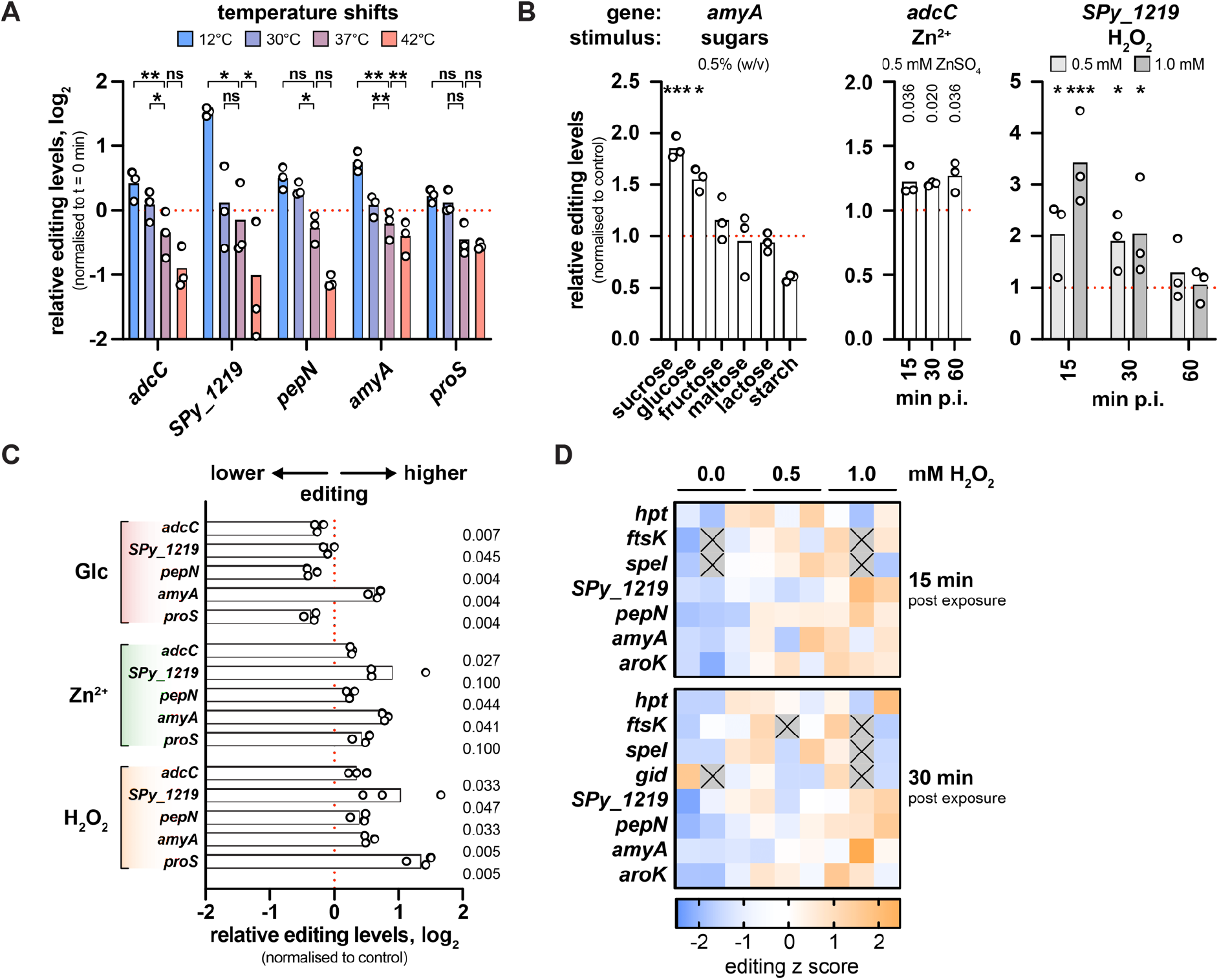
Dynamics of mRNA editing in response to environmental stresses. (A) A-to-I editing levels of selected target genes 15 min after dilution in pre-conditioned culture medium at 12°C, 30°C, 37°C and 42°C. Editing levels were normalised to t = 0 min for each replicate and gene. Statistical analysis was performed using replicate-matched one-way ANOVA with Dunnett’s post-hoc test. (B) Changes in A-to-I editing levels of target genes in response to functionally related conditions. Left: effect of carbohydrate supplementation (0.5% w/v) on *amyA* editing after 30 min, statistical analysis using replicate-matched one-way ANOVA and Dunnett’s post-hoc test. Middle: time-dependent effect of 0.5 mM ZnSO_4_ on *adcC* editing, statistical analysis using paired t-tests with *P*-values adjusted for false discovery rate of 5% (procedure of Benjamini, Krieger and Yekutieli; q values shown for each comparison). Right: time-dependent effect of different doses of H_2_O_2_ (0.5 mM in light grey, 1.0 mM in dark grey) on *SPy_1219* editing, statistical analysis using two-way ANOVA with Dunnett’s post-hoc test. Editing levels were normalised to the mock-treated control for each replicate. p.i. = post induction. (C) A-to-I editing levels of selected target genes 30 min after exposure to the conditions described in (B) (Glc: 0.5% glucose; zinc: 0.5 mM ZnSO_4_; H_2_O_2_: 1 mM H_2_O_2_). Editing levels were normalised to the mock-treated control for each replicate and gene. Statistical analysis was performed using paired t-tests, and *P*-values were adjusted for a false discovery rate of 5% using the procedure of Benjamini, Krieger and Yekutieli. (D) *S. pyogenes* SF370 was treated with 0.5 mM and 1.0 mM H_2_O_2_ at mid-logarithmic growth phase for 15 min and 30 min, and A-to-I editing levels were examined by RNA sequencing. The absolute editing levels of each replicate were transformed into editing z scores and are represented as a heatmap. For replicates depicted as grey tiles with a black cross, editing could not be determined due to filtering steps. *** P < 0.001; ** P < 0.01; * P < 0.05; ns, not significant.

To identify other conditions that modulate A-to-I editing, we tested a range of stimuli related to infection and to the functional category of identified editing target genes, such as carbohydrate availability, zinc stress and oxidative stress. The availability and utilisation of differentially preferred carbohydrates are important regulators of virulence gene expression in *S. pyogenes* (40, 55). The editing target gene *amyA* encodes a secreted starch-degrading enzyme involved in virulence (56), so we examined *amyA* editing in the presence of various carbohydrates. Interestingly, levels of *amyA* editing increased in the presence of sucrose and glucose, while the addition of other mono- and disaccharides (*i.e.*, fructose, maltose, or lactose) did not affect *amyA* editing (Figure 3B, left). In contrast, supplementation with starch resulted in decreased *amyA* editing levels.

Zinc serves as an essential cofactor but is toxic in excess, and the host immune system exploits this ambivalence towards zinc to fight infections (57). Consequently, zinc transport is tightly regulated in *S. pyogenes* (58). Since *adcC*, a component of the AdcABC zinc transporter, is synonymously edited, we analysed the levels of *adcC* editing in response to excess zinc. Compared with the mock-treated control, editing of *adcC* was mildly but consistently increased after exposure to ZnSO_4_ (Figure 3B, middle). Notably, the editing-modulatory effect of zinc excess was still visible 60 min post exposure.

The host immune system employs reactive oxygen species (ROS) to combat bacterial infections, and *S. pyogenes* has developed an arsenal of secreted and cytoplasmic factors to limit ROS toxicity (59). The editing target gene *SPy_1219* has been implicated in redox defence (60), so we studied editing under mild and strong oxidative stress conditions (0.5 mM and 1.0 mM H_2_O_2_, respectively; see Figure S5A). Editing levels of *SPy_1219* increased sharply 15 min and 30 min after H_2_O_2_ challenge, but were largely unaffected 60 min after exposure (Figure 3B, right). A higher concentration of H_2_O_2_ caused a faster and more pronounced increase in *SPy_1219* editing 15 min post exposure, whereas no difference between 0.5 mM and 1.0 mM H_2_O_2_ was observed 30 min after exposure.

Next, we tested whether the observed editing dynamics were exclusive to the selected genes or whether this was a general trend for all editing sites. Although a strong modulator of *amyA* editing, supplementation with glucose did not increase but rather decreased the editing levels of other target genes examined, in particular *pepN* and *proS* (Figure 3C). In contrast, the observed increases in editing upon zinc and oxidative stress were also found for several other editing sites examined, such as *pepN* and *amyA* (Figure 3C), indicating that stress-induced changes in editing are not specific for the mRNAs linked to these stresses.

Given the marked rise in mRNA editing under oxidative stress, we aimed to gain more detailed insights into the dynamics of editing under oxidative stress and thus performed RNA-seq on *S. pyogenes* SF370 exposed to 0.5 mM and 1.0 mM H_2_O_2_ for 15 min and 30 min. Differential expression analysis revealed a major remodelling of the transcriptome upon H_2_O_2_-mediated oxidative stress (Figure S6 and Table S9) and validated the induction of a panel of oxidative stress response genes based on previous publications (Table S10) (59). To examine the effects of oxidative stress on editing levels (Table S8), we focused on editing events that were reliably detected in at least two replicates per condition per timepoint. As observed by Sanger sequencing, editing levels were generally increased both 15 min and 30 min after exposure to H_2_O_2_ compared with the untreated control (Figure 3D). Interestingly, the editing levels of several genes were higher at 1.0 mM than at 0.5 mM H_2_O_2_ (*e.g., SPy_1219* and *aroK* at 15 min and *pepN* at 30 min post exposure), suggesting that editing dynamics vary at least partially with the intensity of stresses.

In conclusion, we found that A-to-I editing is dynamically modulated upon different environmental conditions. While no major differences in the *S. pyogenes* editome were observed across commonly used culture media, several infection-relevant conditions led to global or gene-specific alterations in editing levels.

### Editing dynamics are not governed by *tadA* or target gene expression

Although previous studies have reported condition-dependent changes in bacterial A-to-I editing in response to growth phase and iron availability (21, 23, 24), the mechanisms underlying these dynamic changes remain unexplored. Given the global impact of oxidative stress on A-to-I editing in *S. pyogenes*, we sought to investigate the underlying causes of the observed editing dynamics.

ADAR expression has been found to correlate with A-to-I editing levels in eukaryotes (25), and ectopic overexpression of bacterial *tadA* also resulted in significant increases in editing (see Figure S3B). Therefore, we examined whether changes in *tadA* expression could explain the overall increases in A-to-I editing under stresses. Upon exposure to H_2_O_2_, we indeed observed an increase in *tadA* abundance by qRT-PCR (Figure 4A), Northern blotting (Figure S5C) and in the RNA-seq dataset (Table S9). Interestingly, *tadA* expression also increased in response to zinc exposure, similarly resulting in generally elevated levels of editing (compare Figure 3C and Figure S7A). To determine whether the increased editing levels during oxidative and zinc stress were a direct result of *tadA* induction, we placed *tadA* under the control of an anhydrotetracycline (AHT)-inducible promoter (Figure 4B) and evaluated the editing dynamics upon exposure to H_2_O_2_, ZnSO_4_ or AHT under both native and AHT-inducible *tadA* expression conditions. Under native expression conditions, *tadA* expression increased 2.5-fold when exposed to 1 mM H_2_O_2_ and 1.8-fold when exposed to 0.5 mM ZnSO_4_, whereas no difference was observed in the presence of 100 ng/mL AHT (Figure 4C, left panel). Consistent with these changes in *tadA* abundance, the editing levels of *pepN* increased in the presence of H_2_O_2_ and ZnSO_4_, but remained unaffected in the presence of AHT (Figure 4C, right panel). Under AHT-inducible expression conditions, *tadA* expression was strongly induced by 3.8-fold after addition of AHT, and accordingly, the editing levels of *pepN* also increased (Figure 4D). Strikingly, despite the disruption of the native regulation of *tadA* expression, *pepN* editing levels still markedly rose in the presence of H_2_O_2_ but not zinc (Figure 4D, right panel). A similar behaviour was also observed for a second editing target gene, *fakB2* (Figure S7B). Although *tadA* was under the control of P_tet_, we still observed a moderate increase in *tadA* expression by 1.7-fold in the presence of H_2_O_2_, whereas *tadA* expression remained essentially unaltered when exposed to ZnSO_4_ (Figure 4D, left panel). We cannot completely rule out that the increase in *tadA* expression under oxidative stress contributes to the observed changes in A-to-I editing. However, given that the H_2_O_2_-induced increase in *pepN* editing exceeded the effect of AHT in the AHT-inducible *tadA* expression background, factors other than *tadA* expression appear to govern the dynamics of editing. Consistently, *tadA* expression did not correlate with editing in response to glucose supplementation or temperature shifts (compare Figure S7A).

**Figure 4.**
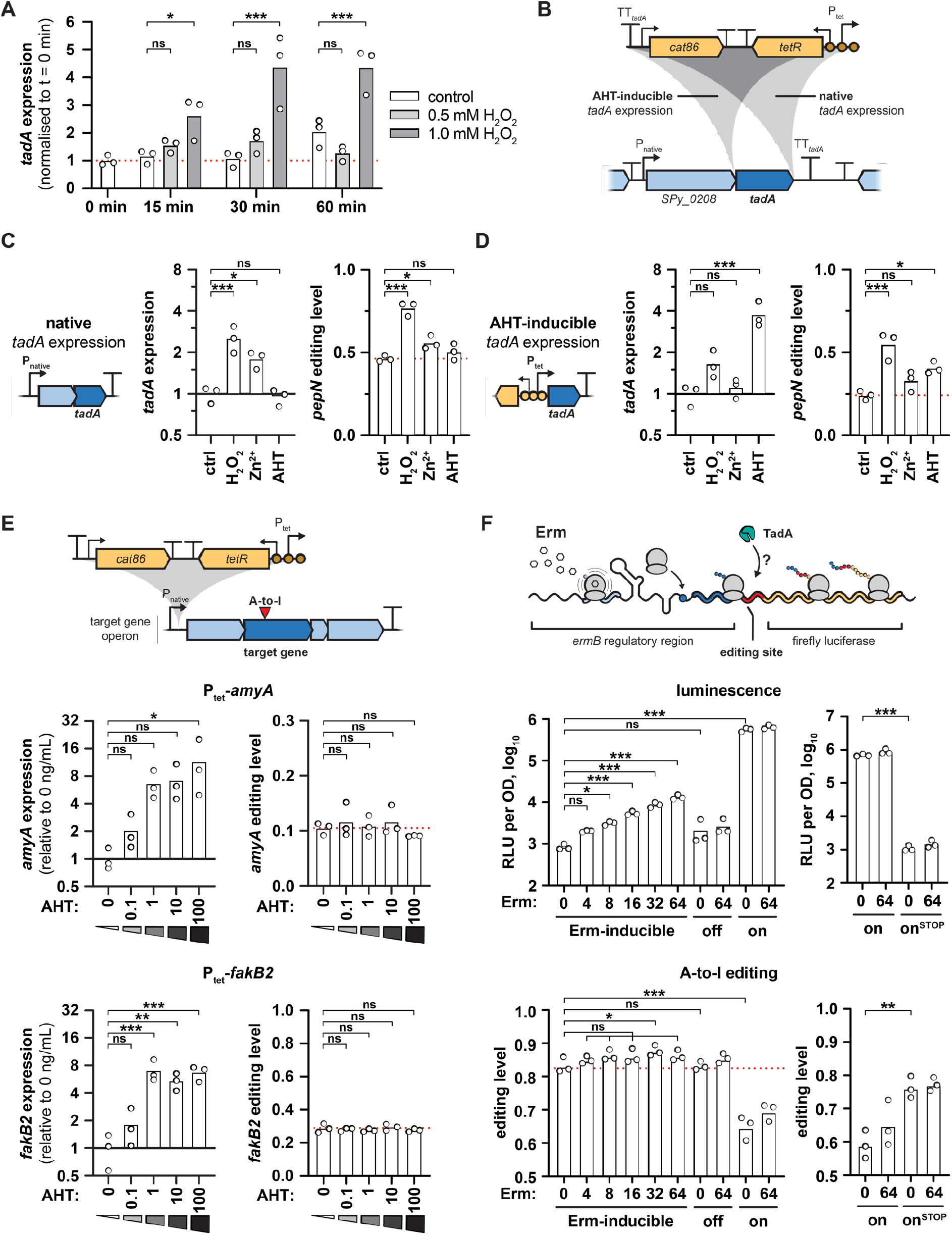
*tadA* expression as well as mRNA expression and translation do not explain the dynamics of stress-dependent mRNA editing. (A) Changes in *tadA* abundance in response to 0.5 mM or 1.0 mM H_2_O_2_ at different time points post exposure in *S. pyogenes* SF370. *gyrA*, *rpoB* and *era* served as reference genes for qRT-PCR, and *tadA* expression was normalised to the mean expression at t = 0 min. Statistical analysis was performed using two-way ANOVA and Dunnett’s post-hoc test. (B) Schematic overview of the *tadA* locus and the integration of the AHT-inducible P_tet_ cassette. The cassette harbouring the antibiotic marker *cat86*, the repressor *tetR* and the promoter with three operator sites (P_tet_) was inserted downstream of the *SPy_0208*-*tadA* operon (‘native’ *tadA* expression, EC3570) or upstream of *tadA* (‘AHT-inducible’ *tadA* expression, EC3622). (C, D) Effect of H_2_O_2_, zinc and AHT exposure on *tadA* abundance (left panel) and *pepN* editing (right panel) under native (C) and inducible *tadA* expression conditions (D). Strains in exponential phase were treated for 30 min with water (control, ‘ctrl’), 1 mM H_2_O_2_, 0.5 mM ZnSO_4_ or 100 ng/mL AHT. *tadA* abundance was measured by qRT-PCR with *gyrA*, *rpoB* and *era* as reference genes, and *pepN* editing levels were examined by Sanger sequencing. Expression and editing levels were normalised relative to the control condition for each strain. Statistical analysis was performed using one-way ANOVA and Dunnett’s post-hoc test. (E) Effect of mRNA abundance on mRNA editing of *amyA* (upper panels) and *fakB2* (lower panels). The AHT-inducible P_tet_ promoter cassette was integrated upstream of two target gene operons (upper scheme), *malX-amyBA-malCDA* (EC3456, upper panels) and *fakB2*-*SPy_1492*-*SPy_1491* operons (EC3459, lower panels), respectively. Strains were grown in the presence of AHT (0 to 100 ng/mL) until mid-logarithmic phase. mRNA expression levels were measured by qRT-PCR with *gyrA* and *ropB* as reference genes (left panels), and editing levels were examined by Sanger sequencing (right panels). Editing and expression were normalised to the respective mean at 0 ng/mL AHT for each strain. Statistical analysis was performed using one-way ANOVA and Dunnett’s post-hoc test. (F) Effect of translation on mRNA editing. The *ermB* regulatory region including the first 30 nt of *ermB* (shades of blue) was fused to an artificial, in-frame A-to-I editing site (red) and the firefly luciferase *ffluc* (yellow). Addition of erythromycin (Erm) causes ribosome stalling on the *ermB* leader peptide but induces translation of the *ermB*’-*ffluc* fusion (upper scheme). *S. pyogenes* SF370 was transformed with the Erm-inducible wildtype reporter (pEC3045), an erythromycin insensitive (off, pEC3046) and a constitutively high translation control (on, pEC3047), and cells were exposed to different concentrations of erythromycin for 30 min at mid-logarithmic growth phase. Luminescence was measured and expressed as log_10_-transformed relative luminescence units (RLU) per OD (upper left panel), and editing levels were examined by Sanger sequencing (lower left panel). As an additional control, *S. pyogenes* SF370 was transformed with the constitutively high translation control reporter (on, pEC3047) and its start codon mutant version (on^STOP^, pEC3069), and cells were treated and analysed as before (right panels). Statistical analysis was performed using one-way ANOVA and Dunnett’s post-hoc test (left panels), or unpaired two-sided t-test (right panels). (A, C-F) *** P < 0.001; ** P < 0.01; * P < 0.05; ns, not significant.

In *E. coli*, oxidative stress slows down translation through the rapid enzymatic degradation of tRNAs (61). We hypothesised that reduced tRNA levels might consequently shift TadA activity towards mRNAs. However, we did not observe a decrease in bulk tRNA abundance nor changes in the abundance of the two TadA targets tRNA^Arg^_ICG_ and tRNA^Leu^_IAG_ in response to oxidative stress (Figure S5B).

Since the abundance of *tadA* and tRNA failed to explain the condition-dependent editing dynamics, we investigated whether changes in target mRNA expression were responsible for changes in their editing levels. To do this, we introduced the AHT-inducible P_tet_ cassette upstream of the *malX*-*amyBA*-*malCDA* and the *fakB2*-*Spy_1492*-*SPy_1491* operons as well as of the non-edited control operon *nga*-*ifs*-*slo* (Figure 4E, upper scheme). In the control strain, the addition of AHT did not affect expression and editing of *amyA* and *fakB2* (Figure S7C). When we adjusted target gene abundance with increasing amounts of AHT in the inducible strains, mRNA levels increased by up to 11.5-fold for *amyA* and by 7.1-fold for *fakB2* in the presence of 100 ng/mL AHT (Figure 4E, left panels). However, editing levels remained unchanged for both *amyA* and *fakB2* at all concentrations of AHT tested (Figure 4E, right panels), indicating that mRNA abundance does not modulate editing. Consistent with this, the A-to-I editing target genes did not show a common response in terms of their gene expression upon H_2_O_2_-mediated oxidative stress in our RNA-seq experiment (Figure S6B).

In bacteria, translation of most mRNAs is globally decreased under oxidative stress (61, 62), leading us to hypothesize that a reduced translation rate might make mRNAs more accessible to TadA. To confirm a decreased translation rate in H_2_O_2_-stressed *S. pyogenes*, we first performed a puromycin incorporation assay and found a consistent reduction in translation after exposure to 1 mM H_2_O_2_ (Figure S7D). To further investigate the role of translation, we constructed a luminescence reporter assay using the translation attenuation mechanism of the leader peptide of *ermB* (63). Addition of erythromycin would result in ribosome stalling on the leader peptide *ermBL*, but increased translation of the *ermB*’-*ffluc* fusion protein harbouring an artificial, in-frame A-to-I editing site (Figure 4F, upper scheme, and Figure S8). We included two mutant reporters that would lead to either erythromycin insensitivity or constitutively high translation. Although luminescence signals – as an indicator of translation rate – increased with increasing concentrations of erythromycin, no change in A-to-I editing was observed for the wild-type reporter and the erythromycin-insensitive control (Figure 4F, left panels). Interestingly, the constitutive translation control displayed reduced editing. To exclude that the deletion of large parts of the *ermB* regulatory region caused structural changes in the mRNA that decreased editing, we mutated the start codon of the fusion protein in this background. Abolishment of translation in this mutant reverted A-to-I editing back to the levels previously observed for the wildtype and the erythromycin-insensitive reporter (Figure 4F, right panels). Our results thus suggest that editing is only affected by strong changes in translation.

### RNA stability is a major determinant of A-to-I editing dynamics

Various stress conditions have been reported to globally stabilize the transcriptome, while nutrient-rich conditions typically increase RNA turnover (64). We therefore examined the stability of selected editing target mRNAs under different stress conditions using the transcription inhibitor rifampicin. RNA half-lives increased significantly under oxidative stress and moderately after zinc exposure, while the addition of glucose resulted in faster turnover of all tested genes, with the exception of *amyA* (Figure S9A). We then compared the calculated RNA half-lives to the editing levels prior to rifampicin addition under the different conditions. RNA half-life and editing were strongly correlated for all tested genes (Figure 5A), suggesting that RNA stability is indeed a major modulator of A-to-I editing.

**Figure 5.**
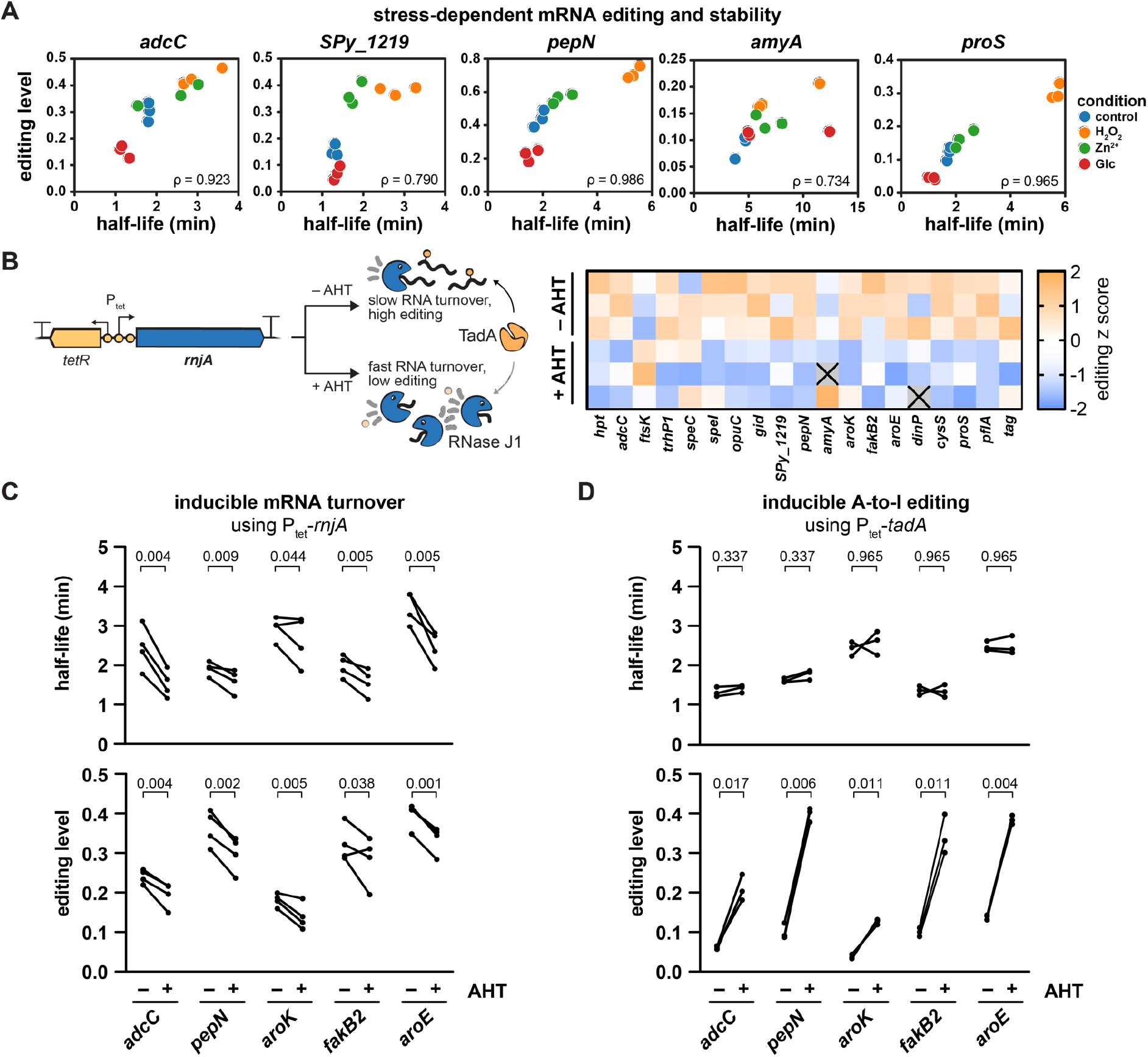
Changes in mRNA stability cause A-to-I editing dynamics. (A) Correlation of A-to-I editing and mRNA half-life in response to H_2_O_2_, zinc and glucose. *S. pyogenes* SF370 was exposed to 1.0 mM H_2_O_2_, 0.5 mM ZnSO_4_ or 0.5% (w/v) glucose at mid-logarithmic phase for 30 min, and rifampicin was added to determine half-lives by qRT-PCR. Half-lives, editing levels and Spearman’s ρ are shown. (B) *S. pyogenes* P_tet_-*rnjA*, harbouring the AHT-inducible P_tet_ promoter to control the expression of the RNase J1-encoding *rnjA*, was grown in the presence or absence of 0.1 ng/mL AHT in THY to mid-logarithmic growth phase, and A-to-I editing was analysed by RNA sequencing. Absolute editing levels were transformed to editing z scores and are shown as a heatmap. (C) *S. pyogenes* P_tet_-*rnjA* was grown in the presence or absence of 0.1 ng/mL AHT in C medium to mid-logarithmic growth phase and treated with rifampicin. Editing levels and mRNA half-lives were analysed as described in (A), and paired replicate values are shown by lines. (D) Rifampicin assay and editing analysis were performed as in (C), but with *S. pyogenes* P_tet_-*tadA* grown in the presence or absence of 100 ng/mL AHT. (C, D) Statistical analysis was performed using paired t-tests, and *P*-values were adjusted for a false discovery rate of 5% using the procedure of Benjamini, Krieger and Yekutieli.

To further validate the effect of RNA stability on editing, we constructed a conditional mutant of the major 5’-to-3’ exoribonuclease RNase J1 (*rnjA*) using the AHT-inducible P_tet_ cassette (65, 66). RNase J is the only bacterial RNase with 5’-to-3’ exonucleolytic and endonucleolytic activity and plays a crucial role in mRNA decay of Gram-positive bacteria (67). In *S. pyogenes*, the paralogous RNases J1 and J2 are essential, and their depletion has previously been shown to prolong mRNA half-lives (66). We envisioned that, in the P_tet_-*rnjA* strain, expression of RNase J1 would increase upon induction with AHT and consequently stimulate global RNA turnover, ultimately leading to a decrease in mRNA half-life and a reduction in editing levels (Figure 5B, left). To obtain a comprehensive picture of the modulatory effect of RNase J1-mediated RNA turnover on editing, we first conducted RNA sequencing and indeed observed reduced editing levels for the majority of target genes upon induction of *rnjA* (Figure 5B and Table S11). To establish the causal link between RNA stability and editing, we validated the interconnection between mRNA half-lives and editing levels using rifampicin. As hypothesised, induction of *rnjA* led to a reduction in mRNA half-lives and a concomitant reduction in editing levels for selected target genes (Figure 5C and Figure S9B).

To prove that the observed changes in mRNA turnover are the cause but not the effect of altered A-to-I editing, we took advantage of the inducible P_tet_-*tadA* strain and compared the half-lives of selected target genes in the presence and absence of AHT. Consistent with our previous results, the levels of editing strongly increased upon induction of *tadA* expression (Figure 5D, lower panel). In contrast, the half-lives of all genes tested remained unaffected by the induction of *tadA* and the associated increase in editing levels (Figure 5D, upper panel, and Figure S9C), thus demonstrating that changes in editing levels are caused by alterations in transcript turnover but not vice versa.

In conclusion, we provide an initial insight into the molecular causes of the dynamics of A-to-I editing. Although *tadA* expression is a potent modulator of A-to-I editing, physiological dynamics of editing are primarily governed by stress-dependent changes in mRNA stability. Our findings highlight an unexpected impact of mRNA stability on A-to-I editing, revealing novel avenues for understanding and potentially manipulating RNA editing in bacteria.

## DISCUSSION

### Expansion of A34-to-I34 editing in *S. pyogenes*

The discovery of additional A34-tRNAs in several bacterial genomes has suggested the expansion of A34-to-I34 editing to tRNAs other than the canonical tRNA^Arg^_ACG_ (8), which was first confirmed for tRNA^Leu^_AAG_ in *O. oeni* (9). In this study, we provide evidence for expanded A34-to-I34 editing in another bacterial species, *S. pyogenes*, and further demonstrate that both A34-tRNAs are fully modified under standard growth conditions (Figure 1) and that editing exclusively relies on the tRNA deaminase TadA. Notably, we found that *S. pyogenes* TadA, but not *E. coli* TadA, was able to efficiently modify the additional tRNA^Leu^_AAG_ (Figure 1C).

In eukaryotes, the expanded set of tRNA substrates is recognised and modified by the TadA-derived ADAT2/3 heterodimer. Recent structural studies suggest that the catalytically inactive ADAT3 subunit initially binds and positions target tRNAs for deamination by the ADAT2 subunit with relatively relaxed sequence specificity (10–12). In contrast, the bacterial homodimer TadA does not require full-length tRNAs for efficient deamination but exhibits very strict sequence specificity (4), raising the question of how TadA has adapted to an extended substrate range. The anticodon loop sequences of the two A34-tRNAs of *S. pyogenes* differ at positions 35 and 37 (compare Figure 2B), thus requiring an increased flexibility of *S. pyogenes* TadA to accommodate both tRNAs. Based on the crystal structure of *S. aureus* TadA in complex with a tRNA-like substrate, bases C35 and G37 of tRNA^Arg^_ACG_ are splayed outward upon binding to TadA and the relatively solvent-exposed base C35 only forms a single hydrogen bond to TadA (14), potentially representing a region more accessible to TadA:tRNA co-evolution. Furthermore, *S. pyogenes* TadA harbours a C-terminal extension absent from *E. coli* and *S. aureus* TadA (68), which might play an additional role in tRNA binding or positioning, thereby contributing to the recognition of the expanded set of tRNA substrates.

The evolutionary processes that led to the expansion of A34-to-I34 editing in eukaryotes remain elusive, but I34-tRNAs have been implicated in proper cell cycle progression (69), codon usage-biased translation (7) and translation of low-complexity proteins (70). Investigating the (co-)evolution of TadA and its substrate tRNAs as well as the physiological functions of expanded A34-to-I34 editing in bacteria will provide valuable information on the biological significance of I34.

### Functional implications of diversified mRNA editing in *S. pyogenes*

Although A-to-I editing of mRNA plays well-documented roles in eukaryotic physiology, our knowledge of the occurrence and function of bacterial mRNA editing remains limited (21, 23, 24). In this study, we identified A-to-I editing sites in *S. pyogenes* and provided insights into the biological plasticity of RNA editing. Consistent with our observation of expanded A34-to-I34 editing in *S. pyogenes*, the adaptation of TadA to a second tRNA substrate also led to a diversification of mRNA editing events compared to *E. coli* and other tRNA^Arg^_ACG_-only bacteria (Table 2). The relaxed sequence specificity of TadA enhances the scope of editing in *S. pyogenes* by Lys-to-Glu recoding and synonymous stop codon editing. Notably, Lys-to-Glu recoding was the most frequent recoding event observed in our study, and the associated change in amino acid charge offers an attractive means of adjusting protein functionality. In contrast, synonymous stop codon editing (UAA-to-UIA) was only observed for a single gene in the oxidative stress dataset (Table S8). No evidence for translational read-through at UIA codons was found in a eukaryote-derived *in vitro* translation assay (71). However, the significance of stop codon editing in bacteria remains unclear.

**Table 2.** Diversification of A-to-I editing in *S. pyogenes*.

| codon position<br>of inosine | tRNA<sup>Arg</sup><sub>ACG</sub>-only species<br>(UACG) |  | *S. pyogenes*<br>(UACG and UAAG) |  |
| --- | --- | --- | --- | --- |
|  | amino acid | codon | amino acid | codon |
| C1 | Thr-to-Ala | ACG-to-ICG | Thr-to-Ala<br>Lys-to-Glu | ACG-to-ICG<br>AAG-to-IAG |
| C2 | Tyr-to-Cys | UAC-to-UIC | Tyr-to-Cys<br>stop codon | UAC-to-UIC<br>UAA-to-UIA |
| C3 | Leu<br>[ Ile-to-Met ]<br>[ Val ] | YUA-to-YUI<br>[ AUA-to-AUI ]<br>[ GUA-to-GUI ] | Leu<br>[ Ile-to-Met ]<br>[ Val ] | YUA-to-YUI<br>[ AUA-to-AUI ]<br>[ GUA-to-GUI ] |

The possible outcomes of A-to-I editing in tRNA<sup>Arg</sup><sub>ACG</sub>-only species, such as *E. coli* or *K. pneumoniae* (left), and *S. pyogenes* (right) are based on the minimum core consensus sequences of each species (in brackets, as reported in (21, 23) and here) and listed according to the presence of inosine at the respective codon position (C1 to C3). Effects are shown at the amino acid and codon level. The editing events in square brackets contain the core consensus sequence but have no pyrimidine at position -2 relative to inosine and are therefore unfavoured.

Interestingly, synonymous editing of Leu codons constituted a substantial part of the *S. pyogenes* editome, whereas it was not observed under standard growth conditions in *E. coli* or *K. pneumoniae* (21, 23). Although differences in codon usage might account for this observation, it is interesting to ask whether the emergence of tRNA^Leu^_IAG_ in *S. pyogenes* has altered the physiological relevance of synonymously edited Leu codons. Indeed, significant peptide truncation was observed at UUI Leu codons in the aforementioned *in vitro* translation assay (71), and the direct modulation of the decoding process at synonymously edited sites may fine-tune translation efficiency and mRNA stability (72–74). The high conservation across different *S. pyogenes* serotypes thus makes synonymously edited Leu codons promising targets for future research.

To date, the physiological consequences of inosine-dependent recoding have only begun to be explored in bacteria. Previous reports suggest that recoding does indeed fine-tune bacterial physiology, ranging from increased toxin toxicity (21) to altered transcriptional regulons (23) to improved iron uptake (24). In addition, A-to-I editing has been proposed to increase transcriptome diversity, potentially contributing to phenotypic variation and adaptation in eukaryotes (75). However, the physiological consequences of individual editing events, particularly those occurring at low frequency, have yet to be determined. Editing events may only become functionally relevant under gene-specific conditions, which are currently unknown. It is also possible that only a few editing events have a defined physiological function, while the majority of sites represents tolerated accessory modifications with no major implications. A global analysis of the contribution of mRNA editing to bacterial physiology is further complicated by the dual function of bacterial TadA in both tRNA and mRNA modification. Although single recoding events can be studied by targeted mutagenesis, more sophisticated approaches are hence required to dissect the RNA type-specific contributions of A-to-I editing to bacterial physiology.

### A-to-I editing dynamics are primarily modulated by mRNA turnover

The dynamic nature of RNA modifications has attracted considerable interest as a means of stress-dependent regulation of gene expression in eukaryotes (76). In this study, we identified several infection-relevant stress conditions that caused substantial alterations in the editing levels of *S. pyogenes*, adding to the growing body of evidence that bacterial mRNA modifications are similarly dynamic (77–79). Although condition-dependent A-to-I editing has previously been reported for bacteria, most work has focused on a single selected target gene and has not provided a more comprehensive view of the dynamics of the editome (21, 23, 24). Among the editing-modulating conditions identified here, oxidative, zinc and temperature stress resulted in an overall change in editing levels across a wide range of target genes. In contrast, elevated glucose levels resulted in decreased editing of most target genes, with the exception of *amyA* (compare Figure 3C), suggesting that the bacterial editome responds non-uniformly to different environmental conditions.

Based on these observations, we sought to investigate the unexplored determinants of bacterial editome dynamics. Bar-Yaacov *et al.* proposed that changes in *tadA* expression and activity, as well as the competition between tRNA and mRNA substrates, are responsible for editing dynamics (22). Although manipulation of *tadA* expression did have profound effects on the *S. pyogenes* editome (Figure 5D, lower panel, and Figure S3B), we identified mRNA stability as the primary determinant of A-to-I editing under physiological conditions. The half-lives of editing target mRNAs positively correlated with their A-to-I editing levels, suggesting that a prolonged transcript lifespan generally increases the probability of the TadA-mRNA interaction and consequently of mRNA modification. We observed a general stabilisation of mRNAs and increased editing under oxidative and zinc stress, while the addition of glucose resulted in destabilisation and reduced editing for all target mRNAs except for *amyA* (Figure 5A).

To demonstrate the effect of transcript stability on A-to-I editing, we used a conditional mutant of the essential RNase J1, the major 5’-to-3’ exoribonuclease in *S. pyogenes* (66, 67), and showed that target mRNA half-life and editing decrease concomitantly upon induction of RNase J1 (Figure 5B and C). Similarly, editing levels increased for selected target sites in a deletion strain of the endoribonuclease RNase Y (Figure S10), which has previously been shown to stabilise the *S. pyogenes* transcriptome by two-fold (37). Notably, increasing editing levels using an inducible *tadA* strain did not result in significant changes in mRNA turnover (Figure 5D). Consequently, mRNA editing levels are largely modulated by mRNA stability but not the opposite way.

The adaptation of RNA stability in response to environmental changes is a well-documented phenomenon. Transcriptome stabilisation is commonly observed under unfavourable conditions, such as nutrient shortage or stress, in various bacterial species (64, 80–83). In *S. pyogenes*, we accordingly observed transcript stabilisation in response to oxidative and zinc stress, but destabilisation in the presence of excess glucose (Figure S9A). The stability of transcripts is intricately regulated by the activity of a diverse set of endo- and exoribonucleases as well as the accessibility of transcript (regions) to these RNases (reviewed in (84, 85)). While the regulation of RNase activity by mechanisms such as post-translational modifications or cellular localisation affects transcriptome stability more globally (86) (compare Figure 5C), transcript-specific features and interactions add an additional layer of regulation to mRNA stability. Access to cleavage sites can be changed by the formation of restrictive secondary structures (87), ribosome occupancy (88), sRNA interactions (89) or RNA modifications (90, 91). Interestingly, the effect of the cellular transcript abundance on mRNA turnover is still a subject of debate, with studies showing negative, positive, or no correlation (64). At least for the transcripts studied here, mRNA abundance and translation did not significantly affect editing in *S. pyogenes* (Figures 4E and F). It will be interesting to explore the extent to which the changes in mRNA editing previously reported by other researchers are also caused by altered mRNA turnover.

Despite the prominent impact of RNA stability, additional features might contribute to editing dynamics. The regulation of A-to-I editing could conceptually follow similar considerations as for the regulation of transcript stability, *i.e.,* global modulation of enzyme activity and transcript-specific changes in recognition site accessibility. Manipulation of *tadA* expression has been shown to be a powerful means to globally modifying editing (Figure S3 and Figure 5D), but little is known about the regulation of *tadA* expression and activity *in vivo*. No major role of *tadA* expression was found for the regulation of editome dynamics under the stressed studied here (Figure 4B-D), but changes in *tadA* expression could become relevant under conditions that remain to be defined. Moreover, the activity of TadA could be further fine-tuned by post-translational modifications or biochemical conditions, as was recently shown for the pH dependency of ADAR catalytic activity (92). In addition to the regulatory potential of TadA, the accessibility of the modification site and the presence of a tRNA anticodon arm-like structure may constitute an important hub for editing regulation. At least in eukaryotes, structural remodelling in response to temperature (compare Figure 3A) or to cross-talking RNA modifications was shown to affect editing efficiency (93–95), and similar effects can easily be envisioned for small RNAs, RNA-binding proteins or the translating ribosome. Lastly, the dual-target specificity of TadA is another unexplored player in the modulation of editing. Although tRNA abundances remained stable in response to oxidative stress (Figure S5B), sudden demands on tRNA abundance and quality could concentrate TadA activity on tRNA rather than mRNA, thereby contributing to global changes in mRNA editing.

In conclusion, our work not only provides a detailed picture of A-to-I editing at the transcriptome scale in a bacterial species with expanded editing capacity, but also demonstrates that stress-dependent changes in mRNA editing are caused by altered transcript turnover. Our findings provide a first glimpse into the network modulating A-to-I editing in bacteria and may thus pave the way for a deeper understanding of the implications of editing on bacterial physiology and virulence.

## DATA AVAILABILITY

All Next Generation sequencing data have been deposited in the European Nucleotide Archive (ENA) under accession PRJEB64678. The RNA-seq data analysed for Figure 2E are available under GEO accessions GSE84641 and GSE40198, and under BioProject accession PRJNA193607. The source code of the editing identification pipeline is available at https://github.com/MPUSP/RNA_editing_analysis_pipeline (DOI: 10.5281/zenodo.8334693).

## AUTHOR CONTRIBUTIONS

Thomas F. Wulff: Conceptualisation, Methodology, Investigation, Validation, Formal analysis, Writing—original draft. Karin Hahnke: Investigation, Validation. Anne-Laure Lécrivain: Methodology, Writing—review & editing. Katja Schmidt: Investigation. Rina Ahmed-Begrich: Formal analysis, Software, Writing—review & editing. Knut Finstermeier: Conceptualisation, Formal analysis, Software, Writing—review & editing. Emmanuelle Charpentier: Conceptualisation, Supervision, Writing—review & editing.

## ACKNOWLEDGEMENTS

The authors gratefully acknowledge Anaïs Le Rhun for conceptual input to this study and the Sequencing Facility of the Max Planck Institute for Molecular Genetics for the preparation and sequencing of NGS libraries. The authors thank the members of the Max Planck Unit for the Science of Pathogens for constructive discussions and critical reading of the manuscript.

