## Supplementary Material for "Dynamics of diversified A-to-I editing in *Streptococcus pyogenes* is governed by changes in mRNA stability"

### SUPPLEMENTARY METHODS

#### Construction of genomic mutants

*General approach*. For anhydrotetracycline (AHT)-inducible gene expression, we used the previously described Tn*10*-derived $P_{tet}$ cassette, harbouring the transcriptional repressor *tetR* with two divergent promoters and three operator sites (hereafter referred to simply as $P_{tet}$), together with *cat86* as antibiotic marker (1, 2), and integrated the cassette into the *S. pyogenes* genome using suicide vectors. The *cat86*-$P_{tet}$ cassette was obtained from pEC536 (2) by restriction digestion with PstI/SmaI and ligated into PstI/SmaI-digested pEC801 to obtain pEC808. An internal NdeI restriction site in *tetR* was synonymously mutated by site-directed mutagenesis of pEC808 using OLEC3322/3323 to obtain pEC812 with *tetR*(A582C).

*Deletion of* tadA. The up- and downstream flanking regions of *tadA* were amplified from SF370 genomic DNA using the primers listed in Table S2. The flanking regions were then PCR-ligated to the lox71-$P_{ermAM/B}$-*ermAM/B*-lox66 cassette and integrated into pEC801 using Gibson Assembly® Master Mix generating pEC2899.

*Inducible expression of RNase J1*. The up- and downstream flanking regions of *rnjA* were amplified from SF370 genomic DNA using the primers listed in Table S2. The flanking regions were then sequentially integrated into pEC812 by restriction-ligation using PstI (upstream) and NdeI (downstream) generating pEC852.

*Inducible expression of editing target gene operons*. The *cat86*-$P_{tet}$ cassette was PCR-amplified from pEC808 and assembled into pEC801 with a terminator upstream of *cat86* and flanked by two Bsp1407I restriction sites using Gibson Assembly® Master Mix resulting in pEC2901. Regions upstream and downstream of the operon transcriptional start site (TSS) were then amplified by Phusion PCR from SF370 wildtype genomic DNA using the primers listed in Table S2 and introduced into Bsp1407I-digested pEC2901 using Gibson Assembly® Master Mix generating pEC2964, pEC2965 and pEC2966.

*Inducible expression of TadA*. The $TT_{tadA}$-*cat86*-$P_{tet}$ cassette was integrated between *SPy_0208* and *tadA*, which together constitute a bicistronic operon. As a control, the $TT_{tadA}$-*cat86*-$P_{tet}$ cassette was integrated directly downstream of the *tadA* coding sequence. The terminator region and the up- and downstream flanking regions were PCR-ligated to the *cat86*-$P_{tet}$

cassette using the primers listed in Table S2. The purified amplicon was introduced in the PCR-linearised suicide vector pEC801 using Gibson Assembly® Master Mix, resulting in pEC2913 and pEC2914. Due to promoter leakage, the ribosome binding site was adapted through site-directed mutagenesis using OLEC13498/13499 or OLEC13498/13500, generating pEC3000 from pEC2913 and pEC3001 from pEC2914, respectively. *tadA* expression was further optimised by site-directed mutagenesis of pEC3001 using OLEC13766/13767, generating pEC3021.

*Electroporation, selection and validation of strains.* Plasmids were linearised by digestion with Cfr42I and introduced into electro-competent *S. pyogenes* SF370 by electroporation (3). Strains were selected for two passages on TSA plates with 3% blood and 6 µg/mL chloramphenicol, cultured in THY without antibiotics, and flash-frozen in liquid nitrogen in THY with 20% glycerol. All strains were validated by Sanger sequencing using gene-specific and *cat86*-$P_{tet}$ cassette-targeting primers as well as primers targeting the virulence regulators *mga*, *covRS* and *ropB* (see Table S2), which are common mutational hotspots affecting *S. pyogenes* virulence (4). A schematic overview of the modified loci is shown in Figure S1.

### Construction of plasmids

*Templates for tRNA* in vitro *transcription.* tRNA template and T7 promoter sequences were assembled from six oligonucleotides in a modular approach using the HindIII and BamHI restriction sites in pUC19 as previously described (5). For *S. pyogenes* tRNAs, the CCA terminus with an overlapping MvaI restriction site was artificially added to the 3′ end of the tRNA gene. In brief, oligonucleotides were 5′ phosphorylated, annealed and ligated into BamHI-/HindIII-digested pUC19. Mutations in the T7 promoter of pEC2405 and pEC2406 were repaired by site-directed mutagenesis using primers OLEC9115/9117 for pEC2405 and OLEC9115/9116 for pEC2406, respectively.

*TadA protein purification.* The *S. pyogenes tadA* gene (NC_002737.2:187,764..188,279) was PCR-amplified from *S. pyogenes* SF370 genomic DNA using primers OLEC8780/8782 and the *E. coli tadA* gene was amplified from *E. coli* NEB® 5-alpha genomic DNA using OLEC8956/8958. pET-21a(+) and PCR amplicons were digested with FastDigest NdeI and FastDigest HindIII, ligated and introduced by transformation into *E. coli*, generating plasmids pEC2360 (*S. pyogenes tadA*) and pEC2389 (*E. coli tadA*).

*Ectopic expression of* tadA *in S. pyogenes.* The constitutive promoter $P_{gyrA}$ from *S. agalactiae* was amplified from pEC455 using OLEC11695/11697. *tadA* was amplified from genomic DNA using OLEC11698/11541, and a C-terminal $His_6$ tag was added by PCR-mediated ligation to the assembled oligos OLEC11059/11060 using the primers OLEC11692/11698. Amplicons were assembled in OLEC11689/11694-amplified pEC2173 by Gibson assembly, generating pEC2813. The empty vector control was built by Gibson assembly of OLEC11695/11696-amplified $P_{gyrA}$ from pEC455 into PCR-amplified pEC2173 generating pEC2812.

*ermBL-based reporter assay.* A schematic overview of the reporter construct is shown in Figure S2. The *ermB* leader peptide (*ermBL*)-containing 5′ UTR of Tn*917 ermB*, 30 nt of the *ermB* CDS, the *ftsK*-derived A-to-I editing site and a $Gly_3Ser$ linker were amplified using OLEC14017/14018 from the assembled oligos OLEC14013 to OLEC14016. The firefly luciferase *ffluc* was amplified from pEC2173 using OLEC14009/14010, and the terminator TT3 was assembled using OLEC14011/14012. All amplicons were ligated by PCR using OLEC13810/14017 and integrated into PCR-amplified pEC2812 using OLEC10676/14008 using Gibson Assembly® Master Mix generating pEC3045. The *ermBL*(M1*) mutant reporter pEC3046 was generated by site-directed mutagenesis using OLEC14019/14020. The mutant reporter pEC3047 with a deletion of *ermBL* and the intergenic region between *ermBL* and *ermB* was constructed with OLEC14135/14136-amplified pEC3045 and assembled oligos OLEC14163/14164 using Gibson Assembly® Master Mix. The start codon mutant of pEC3047 (*i.e.*, pEC3069) was generated by site-directed mutagenesis using OLEC14216/14217.

**Puromycin incorporation assay**

Strain SF370 was grown to mid-logarithmic growth phase, exposed to 1 mM $H_2O_2$ or water as a control for 20 min, and treated with 10 µg/mL puromycin (Sigma-Aldrich) for further 10 min. Proteins were isolated by bead-beating and quantified using Bio-Rad Protein Assay. Equal protein amounts were separated on Any kD Mini-PROTEAN TGX Precast Protein Gels (Bio-Rad), transferred onto 0.45 µm PROTRAN nitrocellulose membranes in 1x Towbin buffer with 20%

methanol and stained with 0.2% Ponceau S in 3% acetic acid as a loading control. After destaining, membranes were blocked in 5% skim milk in 1x TBS with 0.1% Tween-20, incubated first with $\alpha$-puromycin (1:2,000; MABE342, Sigma-Aldrich) and second with HRP-linked ECL Mouse IgG (1:5,000; GE Healthcare), and developed using SuperSignal West Pico Chemiluminescent Substrate (Thermo Scientific).

**Identification of A-to-I editing events by RNA sequencing**

We developed a pipeline to identify A-to-I editing sites based on a recent publication (6). Since libraries of genomic DNA did not contain Unique Molecular Identifiers (UMIs), UMI extraction was performed using UMI Tools (v1.0.1) for RNA samples only. Adapter sequences were removed using Cutadapt (v2.10) (7), and reads were then mapped to the respective reference genome (NC_002737.2 for *S. pyogenes* SF370, NZ_CP008776.1 for *S. pyogenes* 5448) and the PhiX genome using BWA-MEM (v.0.7.17) (8). We sorted and indexed the obtained BAM files using Samtools (v1.9) (9). UMI-based PCR deduplication was performed using UMI Tools (v1.0.1) and reference demultiplexing was performed using Samtools. For paired-end reads, overlapping reads were merged and read duplication artefacts created by the mapping algorithm were removed. We masked homopolymer stretches longer than 4 bp from the reference genome and identified single nucleotide polymorphisms (SNPs). Specifically, we extracted and filtered unique reads for each genomic position based on site-specific nucleotides with a Phred quality score of at least 30 and a minimum distance of at least 4 nt from the end of the reads. We further filtered sites with a minimum coverage of 20, at least two supporting reads per strand direction, and a minimum frequency of 0.01. In RNA datasets, we filtered SNPs for the observation of only two different nucleotides, with one being the reference genome. We removed SNPs in the RNA and DNA datasets positioned at masked sites. Next, we compared the SNPs and their positions in the RNA with the DNA dataset and removed them if they were also observed in the DNA dataset. For the $H_2O_2$ stress experiment, we identified SNPs by comparing only RNA sequences to previously sequenced genomic DNA from strain SF370 (from the dataset for initial identification of editing sites across growth phases). A site was further considered if it was observed in two independent replicates. We then filtered for A-to-G transitions and reported them.

We manually curated reported A-to-I editing sites. To improve comparability with the publication on A-to-I editing in *E. coli* (6) and to retrieve high-confidence sites, we additionally filtered positions for a minimum frequency of 3% in two independent replicates. We excluded sites in ribosomal RNA to avoid biases due to rRNA depletion and because of the presence of known modifications (*e.g.*, $m^6$A1518/1519 in 16S rRNA). We found false-positive hits for position 22 in transfer RNAs. Due to N1 methylation of adenosine-22, guanosine was preferentially introduced during reverse transcription in the context of the corresponding local sequence (10), so we excluded sites at tRNA position 22. We checked editing positions in intergenic regions (IGRs) against the orientation of the transcript's coding sequence (CDS) and removed inconsistent sites. We also excluded identified sites adjacent to the 3′ end of tRNA genes due to the post-transcriptional addition of the CCA terminus. In case of position 450,537 in the 5′ UTR of *SPy_0558*, RT-PCR revealed the absence of a portion of the 5′ UTR in RNA but not gDNA samples, leading to misaligned reads and false-positive site identification (data not shown). Lastly, we manually checked sites for mapping artefacts or unusual processing events using the Integrative Genomics Viewer (11). We confirmed by BLAST analysis that the editing sites identified were not in misalignment-prone paralogous sequences resulting from gene duplication events (31 bp window around editing position; data not shown).

We identified two positions located in poorly annotated regions of the SF370 reference genome. First, position 819,206 is annotated in the CDS of *speI* (*SPy_1007*) at codon position 9, but according to previous reports – given the signal peptide sequence and translation initiation at an alternative GTG start codon – the position lies within codon 43 (updated *speI* annotation: NC_002737.2:819,078..819,857) (12). Second, position 962,967 is located upstream of the annotated *gid* CDS (*SPy_1173*), but analysis of *gid* annotations in different strains suggests translation initiation from an alternative GTG start codon further upstream, and the identified editing position is thus located within codon 6 (updated *gid* annotation: NC_002737.2:962,952..964,346).

### SUPPLEMENTARY FIGURES

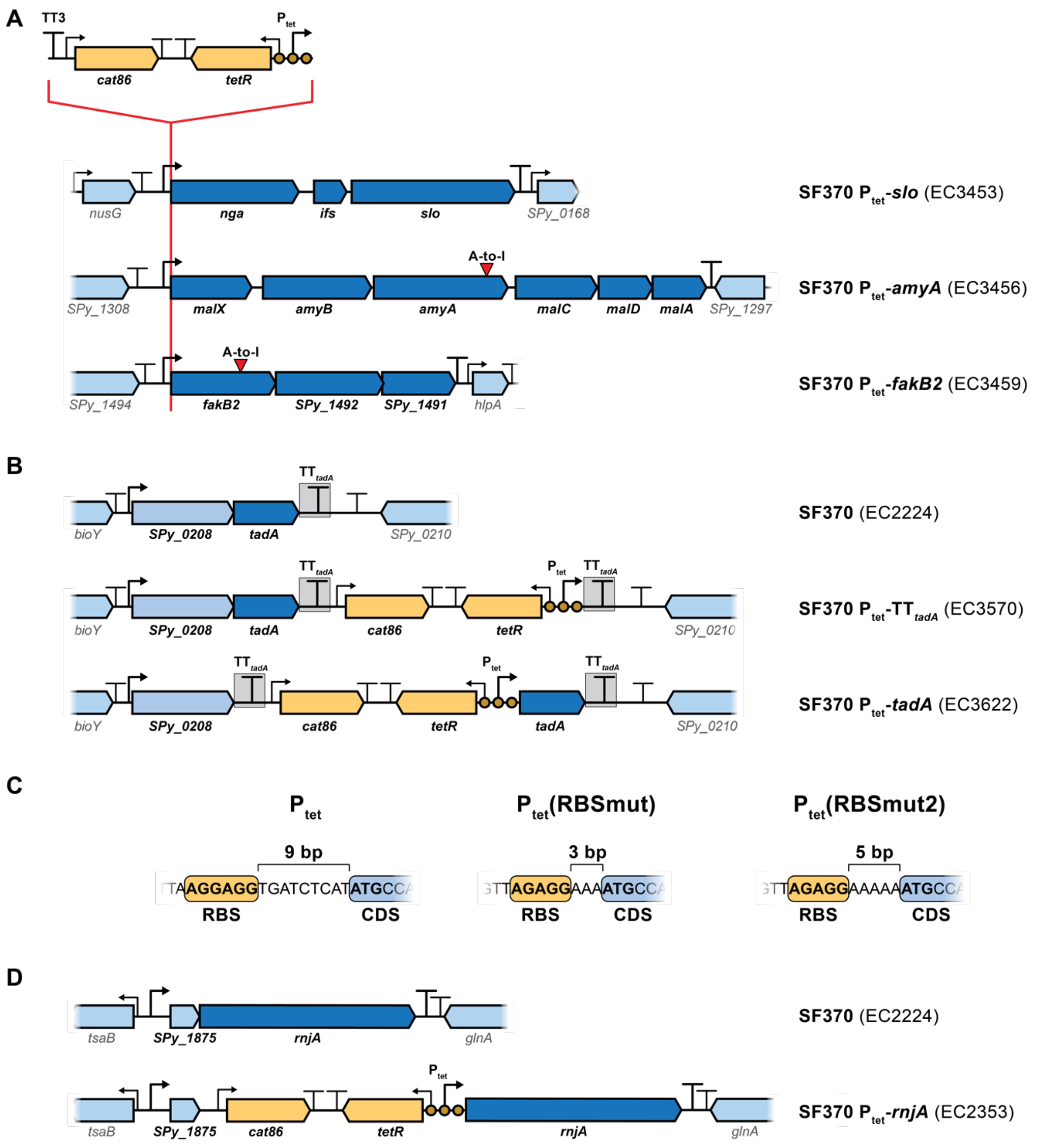

**Figure S1.** Schematic maps of gene loci for P$_{tet}$-harboring strains. (A) To test the influence of gene expression on A-to-I editing levels, the inducible P$_{tet}$ promoter cassette was inserted at the transcriptional start site of two selected editing target operons (*malX-amyBA-malCDA* for strain EC3456 and *fakB2-SPy_1492-SPy_1491* for strain EC3459) and a control operon (*nga-ifs-slo* for strain EC3453). Genes of the affected operons are shown in dark blue, while the surrounding genes are shown in light blue. A-to-I editing sites are marked by red arrows. The

$P_{tet}$ cassette including the *cat86* resistance marker is shown in yellow. Promoters and terminators are indicated. (B) The wildtype locus of the bicistronic *SPy_0208-tadA* operon in strain SF370 (EC2224) is shown in the upper panel with *tadA* highlighted in dark blue and the *tadA* terminator ($TT_{tadA}$) framed by a grey box. For the control strain EC3570, the $P_{tet}$ promoter cassette was inserted downstream of $TT_{tadA}$, and $TT_{tadA}$ was additionally used as terminator directly downstream of the $P_{tet}$ promoter (middle panel). For the inducible $P_{tet}$-*tadA* strain EC3622, the $P_{tet}$ cassette was inserted between *SPy_0208* and *tadA*, and $TT_{tadA}$ was additionally used as terminator for *SPy_0208* (lower panel). (C) Different ribosome binding site (RBS) variants used together with the $P_{tet}$ promoter are shown with the RBS highlighted in yellow and the coding sequence in blue. The distance between RBS and start codon (in bold) is depicted. (D) The wildtype locus of the bicistronic *SPy_1875-rnjA* operon in strain SF370 (EC2224) is shown in the upper panel with *rnjA* highlighted in dark blue. For the inducible $P_{tet}$-*rnjA* strain EC2353, the $P_{tet}$ cassette was inserted between *SPy_1875* and *rnjA* as previously reported (2).

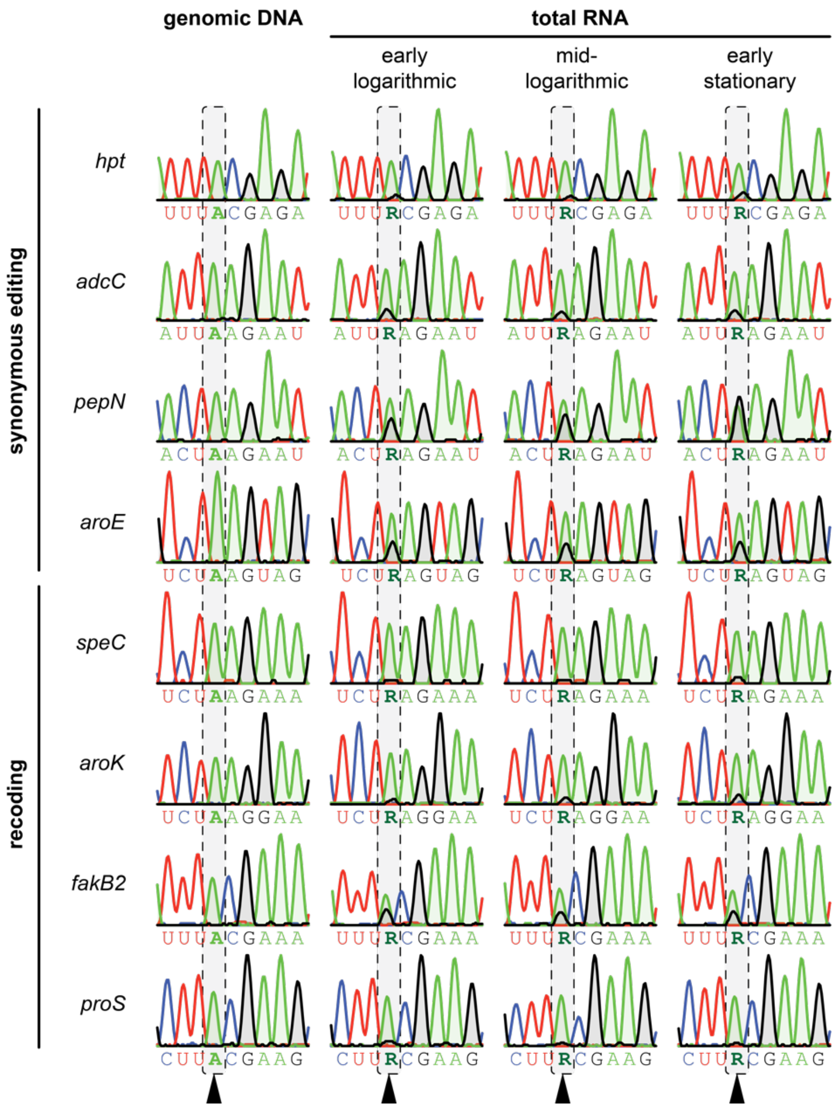

**Figure S2.** Validation of selected A-to-I editing sites in *S. pyogenes* by Sanger sequencing. Total RNA was extracted from early logarithmic, mid-logarithmic and early stationary growth phase. Selected A-to-I editing sites identified by NGS were validated by Sanger sequencing in biologically independent samples. Chromatograms of four selected synonymous editing events are shown in the upper panels and those of four recoding events in the lower panels. Chromatograms for amplicons derived from genomic DNA of SF370 are shown as controls (left). The corresponding editing positions are indicated with black arrows. Nucleotide sequences are shown below each chromatogram. Sequencing traces of adenosine and guanosine are shaded in green and grey, respectively, for better visualisation.

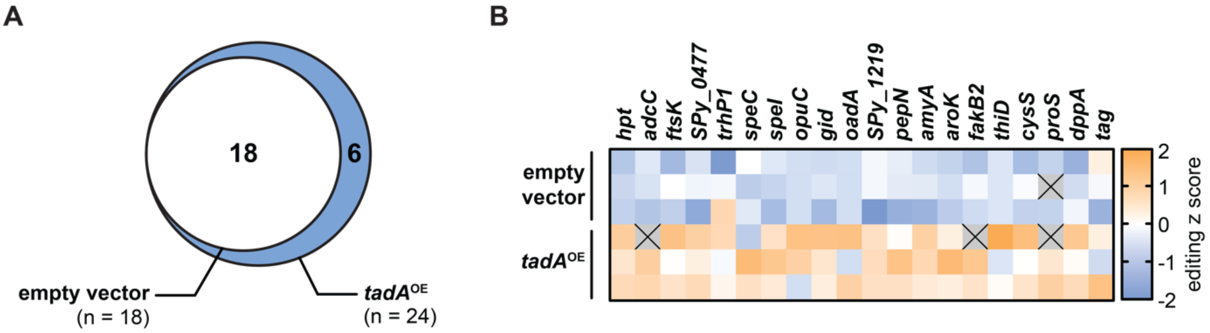

**Figure S3.** Validation of A-to-I editing sites by *tadA* overexpression. A *tadA* overexpression plasmid (pEC2813) and the empty vector control (pEC2812) were introduced by transformation into *S. pyogenes* SF370, and cultures were grown to mid-logarithmic phase in C medium in the presence of kanamycin. Total RNA was extracted and NGS was performed to examine editing levels. (A) Venn diagram of the identified editing sites in the empty vector control and upon overexpression of *tadA*. (B) Editing z scores for each replicate are shown in the heatmap. For replicates depicted as grey tiles with a black cross, editing was not determined due to filtering steps.

| | synonymous editing | | | | | | | | recoding | | | | | | | | | | | | | | | |
|---|---|---|---|---|---|---|---|---|---|---|---|---|---|---|---|---|---|---|---|---|---|---|---|---|
| | *hpt* | *adcC* | *SPy_0558* | *trhP1* | *ppc* | *SPy_1219* | *pepN* | *aroE* | *glpT* | *ftsK* | *SPy_0477* | *SPy_1094* | *opuC* | *amyA* | *aroK* | *fakB2* | *arcD* | *lacA.1* | *dinP* | *thiD* | *cysS* | *proS* | *pflA* | *tag* |
| ***Lla*** | 94 | 47 | 0 | 47 | 0 | 32 | 46 | 49 | 30 | 46 | 70 | 47 | 45 | 0 | 47 | 0 | 0 | 23 | 47 | 46 | 47 | 47 | 0 | 0 |
| ***Sag*** | 50 | 51 | 0 | 51 | 51 | 53 | 51 | 51 | 49 | 51 | 51 | 42 | 51 | 0 | 51 | 51 | 0 | 58 | 51 | 51 | 51 | 51 | 51 | 51 |
| ***San*** | 7 | 7 | 0 | 7 | 7 | 0 | 7 | 7 | 0 | 7 | 7 | 7 | 3 | 0 | 7 | 7 | 7 | 1 | 7 | 7 | 7 | 7 | 7 | 6 |
| ***Sdy*** | 21 | 21 | 0 | 21 | 21 | 21 | 20 | 21 | 15 | 21 | 8 | 21 | 21 | 0 | 21 | 21 | 21 | 31 | 21 | 21 | 21 | 21 | 23 | 22 |
| ***Seq*** | 15 | 15 | 0 | 15 | 15 | 0 | 14 | 15 | 0 | 13 | 15 | 15 | 15 | 15 | 15 | 14 | 15 | 15 | 15 | 15 | 15 | 15 | 15 | 15 |
| ***Sin*** | 6 | 6 | 0 | 6 | 6 | 2 | 6 | 6 | 6 | 6 | 6 | 6 | 6 | 6 | 6 | 6 | 6 | 0 | 6 | 6 | 6 | 6 | 6 | 6 |
| ***Smi*** | 5 | 5 | 0 | 5 | 5 | 0 | 5 | 5 | 0 | 5 | 2 | 0 | 5 | 0 | 5 | 5 | 0 | 5 | 5 | 5 | 5 | 5 | 0 | 5 |
| ***Smu*** | 15 | 15 | 0 | 15 | 14 | 1 | 15 | 15 | 0 | 15 | 15 | 15 | 15 | 0 | 15 | 14 | 0 | 15 | 15 | 15 | 15 | 16 | 15 | 15 |
| ***Spn*** | 53 | 53 | 0 | 52 | 53 | 0 | 53 | 53 | 0 | 53 | 0 | 0 | 53 | 0 | 53 | 53 | 52 | 53 | 53 | 53 | 53 | 53 | 53 | 53 |
| ***Spy*** | 114 | 114 | 49 | 115 | 115 | 115 | 115 | 211 | 115 | 115 | 114 | 115 | 115 | 41 | 115 | 115 | 114 | 230 | 115 | 115 | 115 | 115 | 114 | 115 |
| ***Ssa*** | 11 | 11 | 0 | 11 | 11 | 0 | 11 | 11 | 0 | 11 | 0 | 11 | 11 | 0 | 11 | 11 | 0 | 0 | 11 | 12 | 11 | 11 | 0 | 12 |
| ***Ssu*** | 34 | 34 | 0 | 34 | 34 | 34 | 0 | 34 | 0 | 34 | 0 | 34 | 33 | 3 | 35 | 34 | 33 | 37 | 34 | 50 | 33 | 34 | 32 | 34 |
| ***Sth*** | 35 | 35 | 0 | 35 | 35 | 0 | 35 | 35 | 0 | 34 | 0 | 35 | 0 | 0 | 35 | 35 | 0 | 0 | 35 | 36 | 35 | 36 | 0 | 35 |

**Figure S4.** Conservation of A-to-I editing sites in *Streptococcaceae* members. The number of identified homologs of *S. pyogenes* SF370 editing target genes in other *Streptococcaceae* species is shown. Species: *Lactococcus lactis* (*Lla*), *Streptococcus agalactiae* (*Sag*), *S. anginosus* (*San*), *S. dysgalactiae* (*Sdy*), *S. equi* (*Seq*), *S. iniae* (*Sin*), *S. mitis* (*Smi*), *S. mutans* (*Smu*), *S. pneumoniae* (*Spn*), *S. pyogenes* (*Spy*), *S. salivarius* (*Ssa*), *S. suis* (*Ssu*), and *S. thermophilus* (*Sth*).

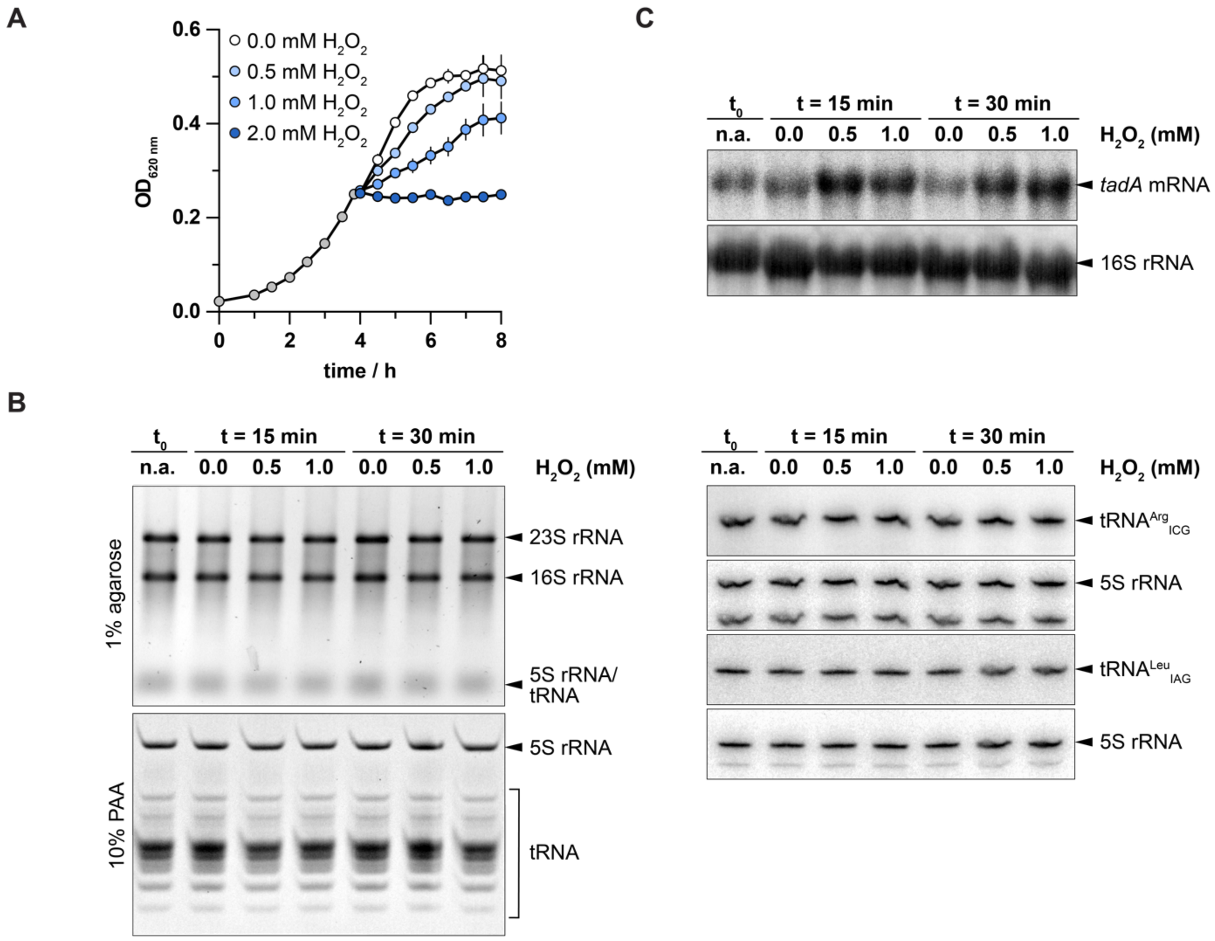

**Figure S5.** Cellular response of *S. pyogenes* SF370 to $H_2O_2$ exposure. (A) *S. pyogenes* was grown to mid-logarithmic growth phase in C medium (grey circles) and challenged with different concentrations of $H_2O_2$ (0.5, 1.0 and 2.0 mM; circles in shades of blue) or $H_2O$ as control (0.0 mM $H_2O_2$, white circles). $OD_{620}$ was measured every 30 min to follow growth. (B) Culture aliquots were taken before and after challenge with $H_2O_2$ for the indicated time and concentration, and total RNA was extracted, separated by denaturing agarose (top left panel) or polyacrylamide gel electrophoresis (bottom left panel) and stained using SYBR™ Gold Nucleic Acid Gel Stain. In addition, the abundances of the two TadA target tRNAs were determined by denaturing polyacrylamide gel electrophoresis and consecutive Northern blotting using 5S rRNA as a loading control (right panel). (C) *tadA* mRNA abundance after $H_2O_2$ exposure was determined by denaturing agarose gel electrophoresis and subsequent Northern blotting using 16S rRNA as a loading control. rRNA, tRNA and mRNA are labelled on the right. Experiments were performed in biological triplicates. n.a. (not applicable)

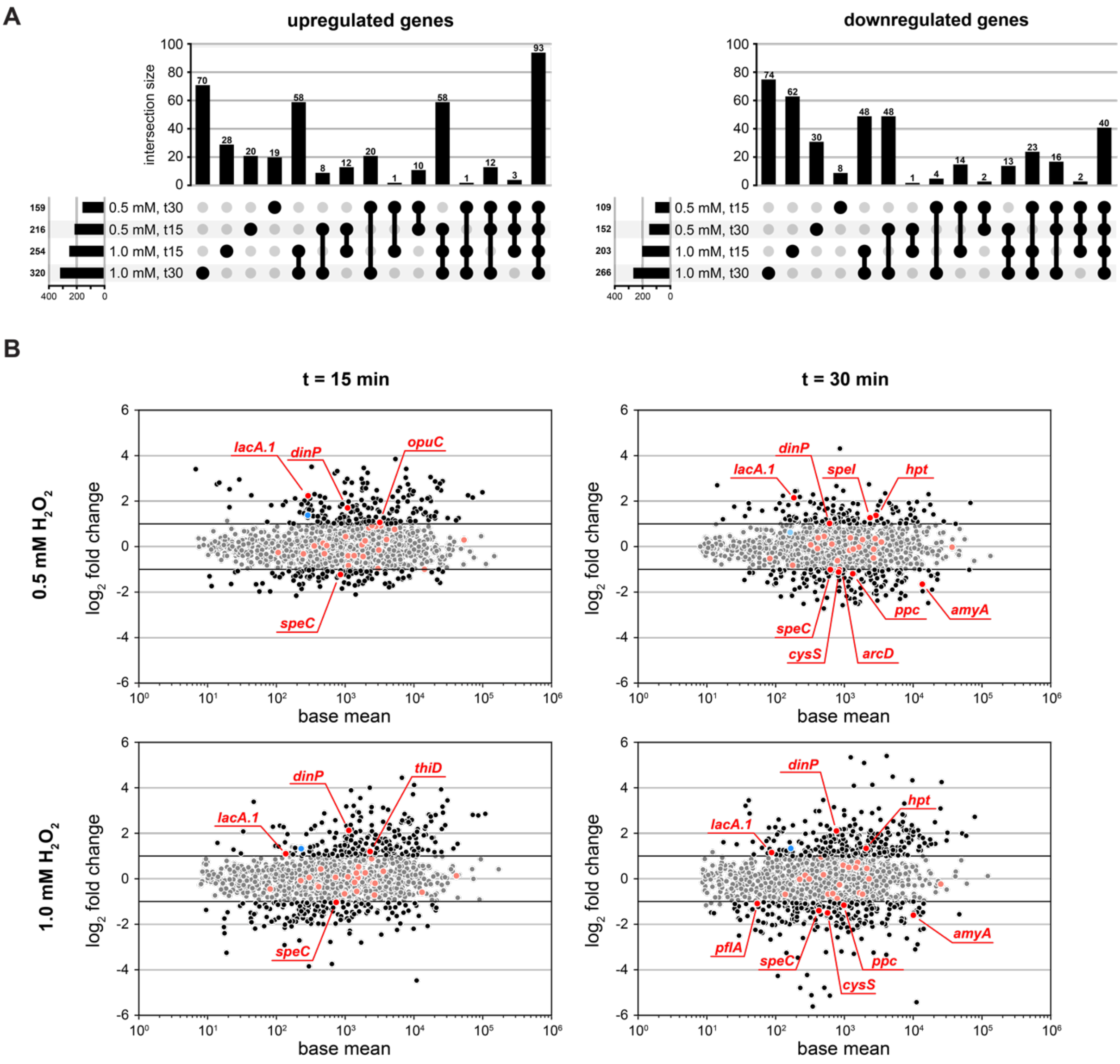

**Figure S6.** Differential expression analysis of *S. pyogenes* SF370 exposed to $H_2O_2$. Differential expression analysis was performed relative to the untreated control at the respective time point with a minimum $log_2$ fold change of 1 or -1 and an adjusted p-value no more than 0.05. (A) The overlaps of differentially expressed genes (including small RNAs) upon challenge of *S. pyogenes* SF370 with different concentrations of $H_2O_2$ for 15 min or 30 min are illustrated by UpSet plots (left: upregulated genes; right: downregulated genes). (B) MA plots are shown for each stress condition with editing target genes (as in Table 1) highlighted in red and *tadA* in blue. Pale circles indicate a non-significant change in expression. Small RNAs were omitted for better visualisation.

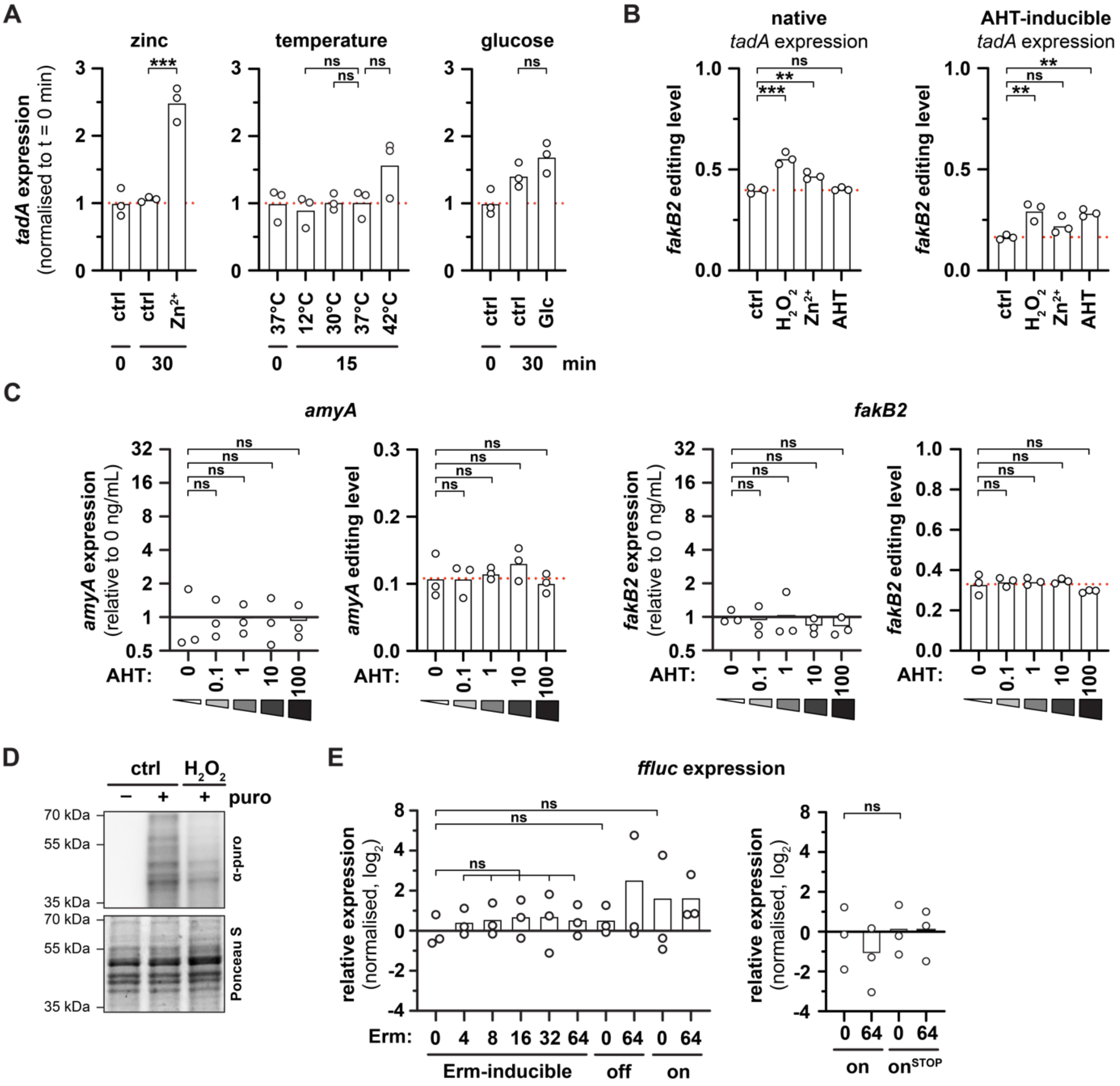

**Figure S7.** The dynamics of stress-dependent mRNA editing is not governed by *tadA* expression, mRNA expression or mRNA translation. (A) Effect of different stimuli on *tadA* expression. *S. pyogenes* SF370 was grown to mid-logarithmic growth phase and exposed for the indicated time to different stimuli. *tadA* expression was then examined by qRT-PCR with *gyrA*, *rpoB* and *era* as reference genes and normalised to the mean expression at t = 0 min. Cells were exposed to 0.5 mM $ZnSO_4$ (left), different temperatures (middle) or 0.5% (w/v) glucose (right). Untreated cultures served as control ("ctrl"). Statistical analysis was performed using one-way ANOVA and Dunnett's post-hoc test (temperature) or unpaired t test (zinc and glucose). (B) Editing of *fakB2* in response to $H_2O_2$, zinc and AHT exposure in the native (left)

and AHT-inducible (right) *tadA* expression strains (compare Figure 4C-D). (C) Effect of AHT on mRNA abundance and mRNA editing of *amyA* (left panels) and *fakB2* (right panels). The AHT-inducible $P_{tet}$ promoter cassette was integrated upstream of the *nga-ifs-slo* operon as control, and the strain was grown and analysed as described for Figure 4E. (D) Changes in translation rate in response to $H_2O_2$. *S. pyogenes* was exposed at mid-logarithmic phase to 1.0 mM $H_2O_2$ for 20 min and treated with puromycin ('puro') for a further 10 min. The rate of translation was examined by Western blotting against puromycin. Results from one exemplary replicate are shown. Statistical analysis was performed using one-way ANOVA and Dunnett's post-hoc test. (E) *ffluc* expression in *ermB* regulatory region-dependent luminescence reporter assay. Strains were grown and treated as described in Figure 4F, and *ffluc* expression was examined by qRT-PCR with *gyrA* and *rpoB* as reference genes, normalised to the mean expression at 0 ng/mL erythromycin (Erm) in the wildtype reporter and $\log_2$ transformed. (A-E) Statistical analysis was performed as described in the main figure, unless otherwise stated. *** $P < 0.001$; ** $P < 0.01$; * $P < 0.05$; ns, not significant.

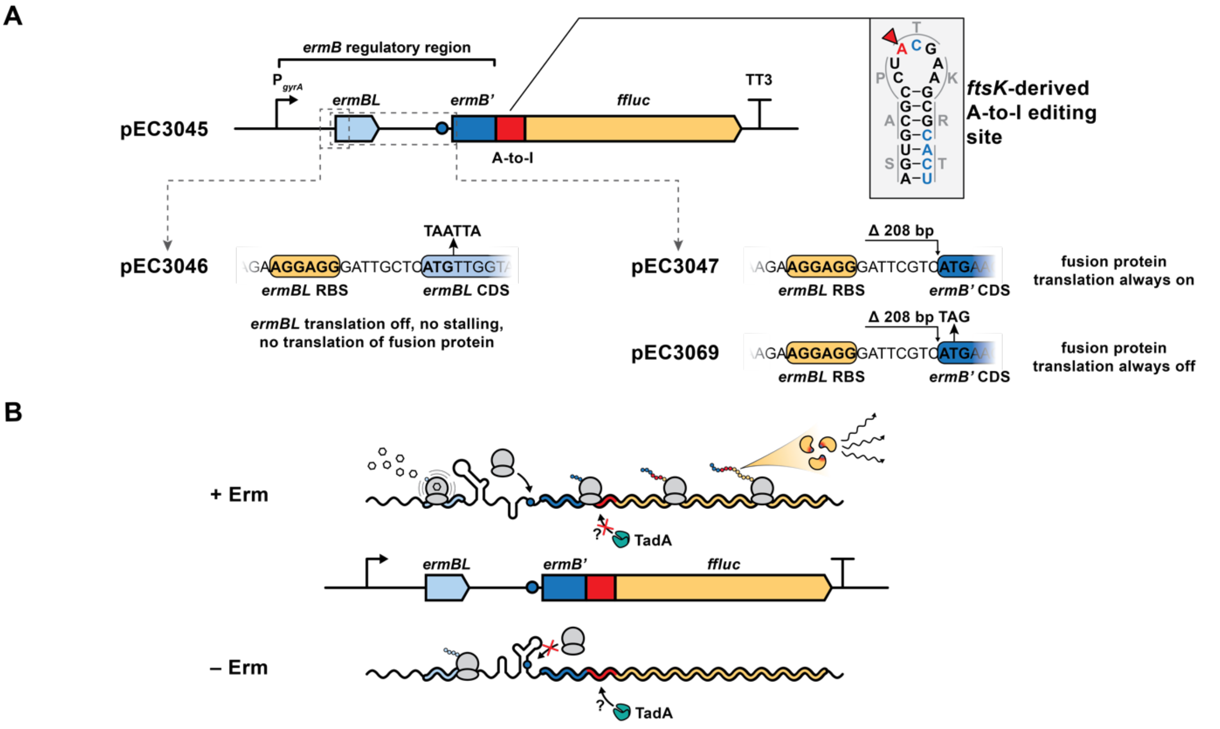

**Figure S8.** Schematic overview of the *ermBL*-based reporter assay. (A) In the wildtype reporter plasmid pEC3045, the constitutive promoter $P_{gyrA}$ drives the transcription of the reporter mRNA, consisting of the *ermB* regulatory region (including the leader peptide *ermBL* in light blue and the first 30 nt of the *ermB* coding sequence in dark blue) with the *ermB* fragment fused in-frame to an *fstK*-like A-to-I editing site (in red) and the firefly luciferase *ffluc* (in orange). The *ftsK*-like editing site was carefully adjusted (blue nucleotides) to strengthen the secondary structure and achieve high editing levels (at the adenosine highlighted in red) (right side). Two mutant versions were designed to (i) shut down translation irrespective of the presence of erythromycin (pEC3046; mutation of start codon of *ermBL*) or to (ii) allow constitutive translation of the fusion protein (pEC3047; deletion of 208 bp comprising *ermBL* and the *ermBL*-*ermB*' intergenic region). As an additional control for pEC3047, the start codon of the constitutively translated fusion protein was mutated (pEC3069). (B) For the wildtype reporter, translation of the fusion protein is inhibited in the absence of erythromycin (– Erm) and the editing site is assumed to be accessible to TadA (in turquoise). In contrast, the presence of erythromycin (+ Erm) results in ribosome stalling in the leader peptide *ermBL* and increased translation of the fusion protein, thereby possibly masking the editing site from TadA.

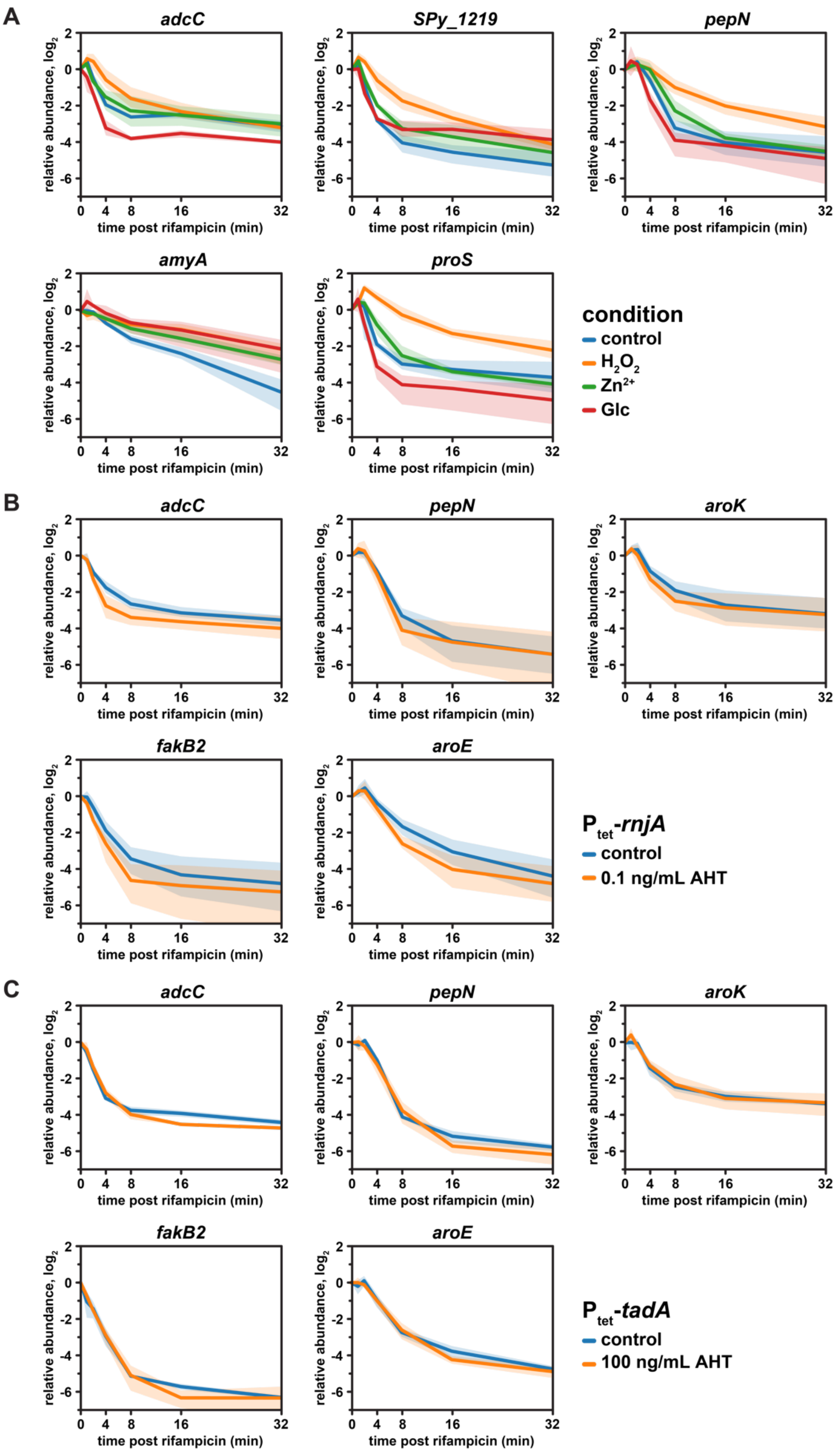

**Figure S9.** Changes in mRNA turnover for selected editing target genes. (A) *S. pyogenes* SF370 was grown to mid-logarithmic phase and exposed to 1.0 mM $H_2O_2$, 0.5 mM $ZnSO_4$ or 0.5%

glucose for 30 min. (B, C) *S. pyogenes* $P_{tet}$-*rnjA* (B) and $P_{tet}$-*tadA* (C) were grown in the presence or absence of AHT to mid-logarithmic growth phase. (A – C) Rifampicin was added at 250 µg/mL and samples were taken right before and after 1 min, 2 min, 4 min, 8 min, 16 min, and 32 min. mRNA abundances were measured by qRT-PCR, normalised first to 16S rRNA levels and second to t = 0 min for each gene. Mean mRNA abundances are shown for each gene as a function of time after rifampicin addition, with standard deviation represented by shaded areas.

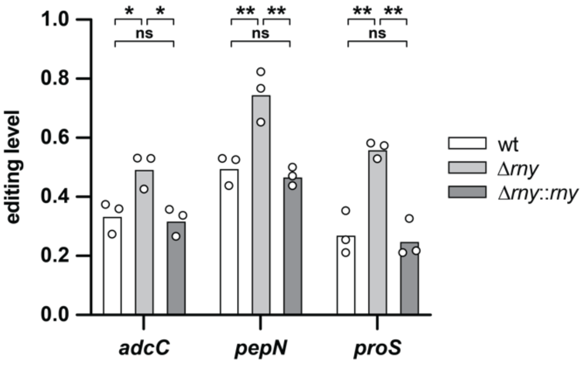

**Figure S10.** Effect of *rny* deletion on A-to-I editing. *S. pyogenes* SF370 wild type, Δ*rny* and Δ*rny*::*rny* were grown to early stationary phase in THY. A-to-I editing levels were analysed by Sanger sequencing for *pepN* and two target genes known to be encoded in RNase Y-processed operons (*adcC* and *proS*) (13). Statistical analysis was performed using one-way ANOVA and Tukey's post-hoc test. ** P < 0.01; * P < 0.05; ns, not significant.

### SUPPLEMENTARY TABLES

**Table S1.** Strains used in this study.

| Strain name | Strain code | Relevant characteristics | Source |
|---|---|---|---|
| ***Streptococcus pyogenes*** | | | |
| **Wild types** | | | |
| SF370 | EC2224 | M1 serotype | ATCC 700294 |
| 5448 | EC2615 | M1T1 serotype | (14) |
| 5448AP | EC2617 | M1T1 serotype, *covS* | (14) |
| **Isogenic mutants** | | | |
| SF370 Δ*rny* | EC2246 | SF370 Δ*rny*::lox72 | (3) |
| SF370 Δ*rny*::*rny* | EC2298 | EC2246 Δlox72::*rny*-TT3-lox72 | (15) |
| SF370 $P_{tet}$-*rnjA* | EC2353 | SF370 *SPy_1875*::*cat86*-$P_{tet}$* | This study |
| SF370 $P_{tet}$-*slo* | EC3453 | SF370 $P_{nga}$::TT3-*cat86*-$P_{tet}$ | This study |
| SF370 $P_{tet}$-*amyA* | EC3456 | SF370 $P_{malX}$::TT3-*cat86*-$P_{tet}$ | This study |
| SF370 $P_{tet}$-*fakB2* | EC3459 | SF370 $P_{fakB2}$::TT3-*cat86*-$P_{tet}$ | This study |
| SF370 $P_{tet}$-$TT_{tadA}$ | EC3570 | SF370 $TT_{tadA}$::*cat86*-$P_{tet}$(RBSmut)-$TT_{tadA}$ | This study |
| SF370 $P_{tet}$-*tadA* | EC3622 | SF370 *SPy_0208*::$TT_{tadA}$-*cat86*-$P_{tet}$(RBSmut2) | This study |
| ***Escherichia coli*** | | | |
| TOP10 | | Host for cloning | Invitrogen |
| NEB 5-alpha | | Host for cloning | NEB |
| NiCo21(DE3) | | Host for protein production | NEB |

Promoters are denoted as P with the related gene name. The AHT-inducible $P_{tet}$ cassette harbours the repressor *tetR* and two divergent promoters with three operator sites, and is separated from the chloramphenicol resistance gene *cat86* by two T4 terminators as previously described (1, 2). A mutant version of $P_{tet}$ with *tetR*(A582C) is shown as $P_{tet}$*. Mutations of the ribosomal binding site as shown in Figure S1 are described as $P_{tet}$(RBSmut) and $P_{tet}$(RBSmut2). Transcriptional terminators are denoted as TT, with TT3 as the phage T3 terminator.

**Table S2.** Oligonucleotides used in this study.

| Purpose | Code | Sequence 5′-3′[a] | F/R[b] | Usage[c] |
|---|---|---|---|---|
| **Sanger sequencing of A34-to-I34 editing of transfer RNAs** | | | | |
| *E. coli argQ* / $tRNA^{Arg}_{ACG}$ | OLEC9821 | *GCATCCGTAGCTCAGCTGG* | F | RT-PCR |
| | OLEC9822 | *TGGTGCATCCGGGAGGATTC* | R | RT |
| | OLEC9823 | TAATACGACTCACTATAGGG*TGGTGCATCCGGGAGGATTC* | R | RT-PCR |
| *SPy_t37* / $tRNA^{Arg}_{ACG}$ | OLEC8776 | *GCACCCTTAGCTCAACTGG* | F | RT-PCR |
| | OLEC8777 | *TGGTGCACCCTAGAGGAG* | R | RT |
| | OLEC9171 | TAATACGACTCACTATAGGG*TGGTGCACCCTAGAGGAG* | R | RT-PCR |
| *SPy_*t52 / $tRNA^{Leu}_{AAG}$ | OLEC8778 | *GCGGGGATGGCGGAATTG* | F | RT-PCR |
| | OLEC8779 | *TGGTGCGGAGAGTGGGAC* | R | RT |
| | OLEC9172 | TAATACGACTCACTATAGGG*TGGTGCGGAGAGTGGGAC* | R | RT-PCR |
| all tRNAs | OLEC1889 | *TAATACGACTCACTATAGGG* | F | SEQ |
| **Sanger sequencing of A-to-I editing of messenger RNAs** | | | | |
| *SPy_0014 / hpt* | OLEC10617 | *CATGGCGGTACATCTAGCAG* | F | RT-PCR, SEQ |
| | OLEC10618 | *CGTAATCTAAACCAAAGCCTACG* | R | RT-PCR |
| *SPy_0093 / adcC* | OLEC11128 | *GAATTTGTCACCATGACCGGTG* | F | RT-PCR |
| | OLEC11129 | *GCTTGCACATGCTCTTCATCG* | R | RT-PCR, SEQ |
| *SPy_0711 / speC* | OLEC10605 | *CAGTCATACTGATTTCTACTATTTCACC* | F | RT-PCR |
| | OLEC10606 | *GGCCTCATAAGACATTTCGG* | R | RT-PCR, SEQ |
| *SPy_1219* | OLEC10623 | *GGATTGGGTACTTAATGTTGC* | F | RT-PCR, SEQ |
| | OLEC10624 | *ATGAGCTAGTGCTTCAATAGC* | R | RT-PCR |
| *SPy_1239 / pepN* | OLEC10626 | *CATGTATCGATGAACCACAAGC* | F | RT-PCR, SEQ |
| | OLEC10627 | *CTTTCCATGAAGTTCTCCTAGC* | R | RT-PCR |
| *SPy_1302 / amyA* | OLEC10614 | *CAAACCAGCAATAGTTACCAAGC* | F | RT-PCR, SEQ |
| | OLEC10615 | *GGTAGATGAGTATCAAAGAACCAG* | R | RT-PCR |
| *SPy_1351 / aroK* | OLEC11118 | *GCAAATGATAATAGTATCATCG* | F | RT-PCR, SEQ |
| | OLEC11119 | *CCATACGTTGTTGATAAAATTC* | R | RT-PCR |
| *SPy_1493 / fakB2* | OLEC11120 | *GGAGGTCGTATTGGTCGTG* | F | RT-PCR |
| | OLEC11121 | *GAGTGGTTGTAATAAGCTGCG* | R | RT-PCR, SEQ |
| *SPy_1584 / aroE* | OLEC11130 | *CTTGGCTGACGAAGGAGTGTC* | F | RT-PCR |
| | OLEC11131 | *GGCTTCATTCCTACACTTGTAGC* | R | RT-PCR, SEQ |
| *SPy_1962 / proS* | OLEC11126 | *CTTACAAGCAATTGCCCTTAAAC* | F | RT-PCR |
| | OLEC11127 | *CGCAATAGACTTGTCAAGAACC* | R | RT-PCR, SEQ |
| *ermB′*-A2I-*ffluc* | OLEC14185 | *CATGAACAAAAATATAAAATATTCTCAAAACAGTGC* | F | RT-PCR |
| | OLEC14186 | *GCATCTGTAAAAGCAATTGTTCC* | R | RT-PCR, SEQ |
| **Quantitative RT-PCR** | | | | |
| *SPy_0093 / adcC* | OLEC14246 | *ACCAAAGGCTGGACGAGTTA* | F | qRT-PCR |
| | OLEC14247 | *CGTAAACGGTGGATGGAAAACC* | R | qRT-PCR |
| *SPy_0098 / rpoB* | OLEC13490 | *CTGTCTTTGACGGGGCTTCA* | F | qRT-PCR |
| | OLEC13491 | *ACCGGTGCGACCATCATAAA* | R | qRT-PCR |
| *SPy_0167 / slo* | OLEC13505 | *ATGAAGCTCCGCCACTCTTT* | F | qRT-PCR |
| | OLEC13506 | *TGCACTAAAGGCCGCTTCAA* | R | qRT-PCR |
| *SPy_0209 / tadA* | OLEC13939 | *TCCCCATTGGCTGTGTCATT* | F | qRT-PCR |

| Purpose | Code | Sequence 5′-3′[a] | F/R[b] | Usage[c] |
|---|---|---|---|---|
| **Sanger sequencing of A34-to-I34 editing of transfer RNAs** | | | | |
| *E. coli argQ* / $tRNA^{Arg}_{ACG}$ | OLEC9821 | *GCATCCGTAGCTCAGCTGG* | F | RT-PCR |
| | OLEC9822 | *TGGTGCATCCGGGAGGATTC* | R | RT |
| | OLEC9823 | TAATACGACTCACTATAGGG*TGGTGCATCCGGGAGGATTC* | R | RT-PCR |
| *SPy_t37* / $tRNA^{Arg}_{ACG}$ | OLEC8776 | *GCACCCTTAGCTCAACTGG* | F | RT-PCR |
| | OLEC8777 | *TGGTGCACCCTAGAGGAG* | R | RT |
| | OLEC9171 | TAATACGACTCACTATAGGG*TGGTGCACCCTAGAGGAG* | R | RT-PCR |
| *SPy_*t52 / $tRNA^{Leu}_{AAG}$ | OLEC8778 | *GCGGGGATGGCGGAATTG* | F | RT-PCR |
| | OLEC8779 | *TGGTGCGGAGAGTGGGAC* | R | RT |
| | OLEC9172 | TAATACGACTCACTATAGGG*TGGTGCGGAGAGTGGGAC* | R | RT-PCR |
| all tRNAs | OLEC1889 | *TAATACGACTCACTATAGGG* | F | SEQ |
| | OLEC13940 | *GCCATCATTTCAGCGTGCAT* | R | qRT-PCR |
| *SPy_0476 / era* | OLEC13488 | *TGACGCAACAAGAGGTTCCA* | F | qRT-PCR |
| | OLEC13489 | *GCTATCGCGCTCCACCATAA* | R | qRT-PCR |
| *SPy_1152 / gyrA* | OLEC13482 | *CCGACTGGTGCCCTTGTTAT* | F | qRT-PCR |
| | OLEC13483 | *CGTTCCCGACCTGTTTGAGT* | R | qRT-PCR |
| *SPy_1219* | OLEC14254 | *TCGTTCTCCTGGCCCTTATT* | F | qRT-PCR |
| | OLEC14255 | *TTCCCCCTGTAGCCATAACA* | R | qRT-PCR |
| *SPy_1239 / pepN* | OLEC14256 | *GCGAGGAGACTGGTCTTTGG* | F | qRT-PCR |
| | OLEC14257 | *GAAGTTCTCCTAGCGCAAACG* | R | qRT-PCR |
| *SPy_1302 / amyA* | OLEC13507 | *AGACAGTACCAGGGGAACAAC* | F | qRT-PCR |
| | OLEC13508 | *ACTTGGCAATGGTCTGAGTGT* | R | qRT-PCR |
| *SPy_1351 / aroK* | OLEC14475 | *TCTTTTGAAACTTTGTACCAGCGT* | F | qRT-PCR |
| | OLEC14476 | *TCCCTCATAAAATACCATACGTTGT* | R | qRT-PCR |
| *SPy_1493 / fakB2* | OLEC13509 | *CGCTTGTCAAAGGTCGAGGTA* | F | qRT-PCR |
| | OLEC13510 | *CAGGCTAGCTTCACCAGCAT* | R | qRT-PCR |
| *SPy_1584 / aroE* | OLEC14473 | *CGCCAAAGAGGTTCGTTTGT* | F | qRT-PCR |
| | OLEC14474 | *GTCCTTGTTAGCTGGTTCAGTT* | R | qRT-PCR |
| *SPy_1805 / secA* | OLEC13486 | *AGGGCGTCAAGTGGAATCTG* | F | qRT-PCR |
| | OLEC13487 | *CGCTGTTCACGCATCACATC* | R | qRT-PCR |
| *SPy_1876 / rnjA* | OLEC14481 | *CTTTGGCGTGAAGCAACTGT* | F | qRT-PCR |
| | OLEC14482 | *CGACCGGTTCTGGAATGGAA* | R | qRT-PCR |
| *SPy_1962 / proS* | OLEC14264 | *AGACGGCTATAGTTTCCATCACA* | F | qRT-PCR |
| | OLEC14265 | *CGCCATCTCCAATAATCCCTTTG* | R | qRT-PCR |
| *ffluc* | OLEC14238 | *AGAGATACGCCCTGGTTCCT* | F | qRT-PCR |
| | OLEC14239 | *TGCCAACCGAACGGACATTT* | R | qRT-PCR |
| 16S rRNA | OLEC14228 | *TGAGTGCAGAAGGGGAGAGT* | F | qRT-PCR |
| | OLEC14229 | *GAGCCTCAGCGTCAGTTACA* | R | qRT-PCR |
| **Probes for Northern blotting** | | | | |
| *SPy_t16* / $tRNA^{Ser}_{UGA}$ | OLEC10579 | *GCACGCTTTTACACGCCTGACG* | R | NB |
| *SPy_t37* / $tRNA^{Arg}_{ACG}$ | OLEC9362 | *GAACCTCTAACCGCCTGATT* | R | NB |

| Purpose | Code | Sequence 5′-3′[a] | F/R[b] | Usage[c] |
|---|---|---|---|---|
| **Sanger sequencing of A34-to-I34 editing of transfer RNAs** | | | | |
| *E. coli argQ* / tRNA$^{Arg}_{ACG}$ | OLEC9821 | *GCATCCGTAGCTCAGCTGG* | F | RT-PCR |
| | OLEC9822 | *TGGTGCATCCGGGAGGATTC* | R | RT |
| | OLEC9823 | TAATACGACTCACTATAGGG*TGGTGCATCCGGGAGGATTC* | R | RT-PCR |
| *SPy_t37* / tRNA$^{Arg}_{ACG}$ | OLEC8776 | *GCACCCTTAGCTCAACTGG* | F | RT-PCR |
| | OLEC8777 | *TGGTGCACCCTAGAGGAG* | R | RT |
| | OLEC9171 | TAATACGACTCACTATAGGG*TGGTGCACCCTAGAGGAG* | R | RT-PCR |
| *SPy*_t52 / tRNA$^{Leu}_{AAG}$ | OLEC8778 | *GCGGGGATGGCGGAATTG* | F | RT-PCR |
| | OLEC8779 | *TGGTGCGGAGAGTGGGAC* | R | RT |
| | OLEC9172 | TAATACGACTCACTATAGGG*TGGTGCGGAGAGTGGGAC* | R | RT-PCR |
| all tRNAs | OLEC1889 | *TAATACGACTCACTATAGGG* | F | SEQ |
| *SPy*_t52 / tRNA$^{Leu}_{AAG}$ | OLEC9363 | *CACACGACCTAAAGCGGTCAC* | R | NB |
| 5S rRNA | OLEC288 | *CTAAGCGACTACCTTATCTCA* | R | NB |
| **Templates for *in vitro* transcriptions** | | | | |
| pEC2322 | OLEC8611 | AGCT**TAATACGACTCACTATAG***CGGGGATGGCGGAATTGGC* | F | OA |
| | OLEC8612 | *AGACGCGCAGGACTAAGGATCCTGTGACCGCTTTAGGTCG* | F | OA |
| | OLEC8613 | *TGTGGGTTCAAGTCCCACTCTCCGCACCAGG* | F | OA |
| | OLEC8614 | *GCGTCTGCCAATTCCGCCATCCCCG***CTATAGTGAGTCGTATTA** | R | OA |
| | OLEC8615 | *CCCACACGACCTAAAGCGGTCACAGGATCCTTAGTCCTGC* | R | OA |
| | OLEC8616 | GATC*CCTGGTGCGGAGAGTGGGACTTGAA* | R | OA |
| pEC2405 | OLEC8926 | AGCTTAATACGACGACTCACTATA*GCACCCTTAG* | F | OA |
| | OLEC8927 | *CTCAACTGGATAGAGTACCTGACTACGAATCAGGCGGT* | F | OA |
| | OLEC8928 | *TAGAGGTTCGACTCCTCTAGGGTGCACCAG* | F | OA |
| | OLEC8929 | GAT*CCTGGTGCACCCTAGAGGAGTCGA* | R | OA |
| | OLEC8930 | *ACCTCTAACCGCCTGATTCGTAGTCAGGTACTCTATCCA* | R | OA |
| | OLEC8931 | *GTTGAGCTAAGGGTGC*TATAGTGAGTCGTCGTATTA | R | OA |
| | OLEC9115 | **CTATAGTGAGTCGTATTA**AGCTTGGCGTAATCATGGTCATAGC | R | SDM-PCR |
| | OLEC9117 | CCAAGCT**TAATACGACTCACTATAG***CACCCTTAGCTCAACTG* | F | SDM-PCR |
| pEC2406 | OLEC8920 | AGCTTAATACGACGACTCACTATA*GCATCCGTAGCTCAGC* | F | OA |
| | OLEC8921 | *TGGATAGAGTACTCGGCTACGAACCGAGCGGTC* | F | OA |
| | OLEC8922 | *GGAGGTTCGAATCCTCCCGGATGCACCAG* | F | OA |
| | OLEC8923 | GAT*CCTGGTGCATCCGGGAGGATTCGAACCTCCGACCGC* | R | OA |
| | OLEC8924 | *TCGGTTCGTAGCCGAGTACTCTATCCAGCTGAG* | R | OA |

| Purpose | Code | Sequence 5′-3′[a] | F/R[b] | Usage[c] |
|---|---|---|---|---|
| **Sanger sequencing of A34-to-I34 editing of transfer RNAs** | | | | |
| *E. coli argQ* / $\mathrm{tRNA}^{\mathrm{Arg}}_{\mathrm{ACG}}$ | OLEC9821 | *GCATCCGTAGCTCAGCTGG* | F | RT-PCR |
| | OLEC9822 | *TGGTGCATCCGGGAGGATTC* | R | RT |
| | OLEC9823 | TAATACGACTCACTATAGGG*TGGTGCATCCGGGAGGATTC* | R | RT-PCR |
| *SPy_t37* / $\mathrm{tRNA}^{\mathrm{Arg}}_{\mathrm{ACG}}$ | OLEC8776 | *GCACCCTTAGCTCAACTGG* | F | RT-PCR |
| | OLEC8777 | *TGGTGCACCCTAGAGGAG* | R | RT |
| | OLEC9171 | TAATACGACTCACTATAGGG*TGGTGCACCCTAGAGGAG* | R | RT-PCR |
| *SPy*_t52 / $\mathrm{tRNA}^{\mathrm{Leu}}_{\mathrm{AAG}}$ | OLEC8778 | *GCGGGGATGGCGGAATTG* | F | RT-PCR |
| | OLEC8779 | *TGGTGCGGAGAGTGGGAC* | R | RT |
| | OLEC9172 | TAATACGACTCACTATAGGG*TGGTGCGGAGAGTGGGAC* | R | RT-PCR |
| all tRNAs | OLEC1889 | *TAATACGACTCACTATAGGG* | F | SEQ |
| | OLEC8925 | *CTACGGATGC*TATAGTGAGTCGTCGTATTA | R | OA |
| | OLEC9115 | **CTATAGTGAGTCGTATTA**AGCTTGGCGTAATCATGGTCATAGC | R | SDM-PCR |
| | OLEC9116 | CCAAGCT**TAATACGACTCACTATAG***CATCCGTAGCTCAGC* | F | SDM-PCR |
| **TadA protein production** | | | | |
| pEC2360 | OLEC8780 | AGCACAT*ATGCCATATAGTTTAGAAGAGC* | F | PCR |
| | OLEC8782 | ACGTAAGCTT*GTCAAAGGGATCTGACTG* | R | PCR |
| pEC2389 | OLEC8956 | GATCCAT*ATGTCTGAAGTCGAATTTAGCCACG* | F | PCR |
| | OLEC8958 | AGCGAAGCTT*ATCCGTCGAGGATTGCGC* | R | PCR |
| **Ectopic expression of *tadA* in *S. pyogenes*** | | | | |
| pEC2173 linearisation | OLEC11689 | AAGGCCCACTTTTGTGGGCCTTTTTT*GGATCGGCCGCTCTAGAGTC* | F | PCR |
| | OLEC11694 | *GAGCTCCAATTCGCCCTATAGTGAG* | R | PCR |
| pEC2812 | OLEC11695 | CTATAGGGCGAATTGGAGCTC*GCCTATCATTTTCAATGAAAGAAGTC* | F | PCR |
| | OLEC11696 | AAGGCCCACAAAAGTGGGCCTTTTTTT*TAAAAAATGCCCCTTTTCTTCAC* | R | PCR |
| pEC2813 | OLEC11695 | CTATAGGGCGAATTGGAGCTC*GCCTATCATTTTCAATGAAAGAAGTC* | F | PCR |
| | OLEC11697 | CTAAACTATATGGCATT*AAAAAATGCCCCTTTTCTTCACTG* | R | PCR |
| | OLEC11692 | AAGGCCCACAAAAGTGGGCCTTTTTT*CTAATGGTGATGATGGTGATGAGAACCTCC* | R | PCR |
| | OLEC11698 | GGGCATTTTTTA*ATGCCATATAGTTTAGAAGAGCAAACTTATTTC* | F | PCR |
| | OLEC11541 | *GTCAAAGGGATCTGACTGTTC* | R | PCR |
| | OLEC11059 | CAAGGAACAGTCAGATCCCTTTGACGGTGGAGGTTCTCATCACCATCATCACCATTAGAAAAAAGGCCCACTTTTGTGGGCCTTTTTTTAACGCTGATAGTGCTAGTGTAG | F | OA |
| | OLEC11060 | CTACACTAGCACTATCAGCGTTAAAAAAAGGCCCACAAAAGTGGGCCTTTTTTCTAATGGTGATGATGGTGA | R | OA |

| **Purpose** | **Code** | **Sequence 5′-3′[a]** | **F/R[b]** | **Usage[c]** |
|---|---|---|---|---|
| **Sanger sequencing of A34-to-I34 editing of transfer RNAs** | | | | |
| *E. coli argQ* / tRNA$^{Arg}_{ACG}$ | OLEC9821 | *GCATCCGTAGCTCAGCTGG* | F | RT-PCR |
| | OLEC9822 | *TGGTGCATCCGGGAGGATTC* | R | RT |
| | OLEC9823 | TAATACGACTCACTATAGGG*TGGTGCATCCGGGAGGATTC* | R | RT-PCR |
| *SPy_t37* / tRNA$^{Arg}_{ACG}$ | OLEC8776 | *GCACCCTTAGCTCAACTGG* | F | RT-PCR |
| | OLEC8777 | *TGGTGCACCCTAGAGGAG* | R | RT |
| | OLEC9171 | TAATACGACTCACTATAGGG*TGGTGCACCCTAGAGGAG* | R | RT-PCR |
| *SPy*_t52 / tRNA$^{Leu}_{AAG}$ | OLEC8778 | *GCGGGGATGGCGGAATTG* | F | RT-PCR |
| | OLEC8779 | *TGGTGCGGAGAGTGGGAC* | R | RT |
| | OLEC9172 | TAATACGACTCACTATAGGG*TGGTGCGGAGAGTGGGAC* | R | RT-PCR |
| all tRNAs | OLEC1889 | *TAATACGACTCACTATAGGG* | F | SEQ |
| | | TGAGAACCTCCACCGTCAAAGGGATCTGACTGTTCCTTG | | |
| ***ermBL*-based editing reporter assay** | | | | |
| *ermB*(5′UTR)-*ermB′*:A2I (for pEC3045) | OLEC14013 | CCTAAATTATGGTACAATGTAAGAGGAAGTTAAATTAGATGCTAAAAATTTGTAATTAAGAAGGAGGGATTCGTCATGTTGGTATTCCAAATGCGTAATGTAGATAAAACAT | F | OA |
| | OLEC14014 | AGTCATAAGATTAGTCACTGGTAGGAATTAATCTAACGTATTTATTTATCTGCGTAATCACTGTTTTTAGTCTGTTTCAAAACAGTAGATGTTTTATCTACATTACGCATTTGGAATA | R | OA |
| | OLEC14015 | AATTCCTACCAGTGACTAATCTTATGACTTTTTAAACAGATAACTAAAATTACAAACAAATCGTTTAACTTCTGTATTTATTTATAGATGTAATCACTTCAGGAGTGATTACATGAAC | F | OA |
| | OLEC14016 | CTTTCTTTATGTTTTTGGCGTCTTCAGAACCTCCACCAGTACGCTTCGTAGGCGCACTGTTTTGAGAATATTTTATATTTTTGTTCATGTAATCACTCCTGAAGTGATTACA | R | OA |
| | OLEC14017 | *CCTAAATTATGGTACAATGTAAGAGGAAGT* | F | PCR |
| | OLEC14018 | *CTTTCTTTATGTTTTTGGCGTCTTCAG* | R | PCR |
| *ffluc* (for pEC3045) | OLEC14009 | *GAAGACGCCAAAAACATAAAGAAAGG* | F | PCR |
| | OLEC14010 | *TTACAATTTGGACTTTCCGCCC* | R | PCR |
| TT3 (for pEC3045) | OLEC14011 | GAAGGGCGGAAAGTCCAAATTGTAATAGAAAAAAGGCCCACTTTTGTGGGCCTTTTTTGGATCGGCCGCTCTAGAGTCGTGTT | F | OA |
| | OLEC14012 | AACACGACTCTAGAGCGGCCGATCCAAAAAAGGCCCACAAAAGTGGGCCTTTTTTCTATTACAATTTGGACTTTCCGCCCTTC | R | OA |
| *ermB*(5′UTR)-*ermB′*:A2I:*ffluc*-TT3 (for pEC3045) | OLEC14017 | *CCTAAATTATGGTACAATGTAAGAGGAAGT* | F | PCR |
| | OLEC13810 | *CGACTCTAGAGCGGCCGATC* | R | PCR |
| | OLEC10676 | *GGATCGGCCGCTCTAGAG* | F | PCR |

| **Purpose** | **Code** | **Sequence 5′-3′[a]** | **F/R[b]** | **Usage[c]** |
|---|---|---|---|---|
| **Sanger sequencing of A34-to-I34 editing of transfer RNAs** | | | | |
| *E. coli argQ* / $tRNA^{Arg}_{ACG}$ | OLEC9821 | *GCATCCGTAGCTCAGCTGG* | F | RT-PCR |
| | OLEC9822 | *TGGTGCATCCGGGAGGATTC* | R | RT |
| | OLEC9823 | TAATACGACTCACTATAGGG*TGGTGCATCCGGGAGGATTC* | R | RT-PCR |
| *SPy_t37* / $tRNA^{Arg}_{ACG}$ | OLEC8776 | *GCACCCTTAGCTCAACTGG* | F | RT-PCR |
| | OLEC8777 | *TGGTGCACCCTAGAGGAG* | R | RT |
| | OLEC9171 | TAATACGACTCACTATAGGG*TGGTGCACCCTAGAGGAG* | R | RT-PCR |
| *SPy*_t52 / $tRNA^{Leu}_{AAG}$ | OLEC8778 | *GCGGGGATGGCGGAATTG* | F | RT-PCR |
| | OLEC8779 | *TGGTGCGGAGAGTGGGAC* | R | RT |
| | OLEC9172 | TAATACGACTCACTATAGGG*TGGTGCGGAGAGTGGGAC* | R | RT-PCR |
| all tRNAs | OLEC1889 | *TAATACGACTCACTATAGGG* | F | SEQ |
| pEC2173 backbone with P$_{gyrA}$ | OLEC14008 | *CTCTTACATTGTACCATAATTTAGGTAAAATTGC* | R | PCR |
| pEC3046 | OLEC14019 | *GGATTCGTCTAATTAGTATTCCAAATGCGTAATGTAGATAAAACATCTACTG* | F | SDM-PCR |
| | OLEC14020 | *TTGGAATACTAATTAGACGAATCCCTCCTTCTTAATTACAAATTTTTAGC* | R | SDM-PCR |
| pEC3047 | OLEC14135 | *CAAAAATATAAAATATTCTCAAAACAGTGC* | F | PCR |
| | OLEC14136 | *CTGAAGTGATTACATCTATAAATAAATACAG* | R | PCR |
| | OLEC14163 | *AGGGATTCGTCATGAACAAAAATATAAAATATTCTCAAAACAGTGCG* | F | SDM-PCR |
| | OLEC14164 | *TATTTTTGTTCATGACGAATCCCTCCTTCTTAATTACAAATTTTTAG* | R | SDM-PCR |
| pEC3069 | OLEC14216 | *AGGGATTCGTCTAGAACAAAAATATAAAATATTCTCAAAACAGTGCG* | F | SDM-PCR |
| | OLEC14217 | *TATTTTTGTTCTAGACGAATCCCTCCTTCTTAATTACAAATTTTTAG* | R | SDM-PCR |
| **Suicide vector template for deletion of *tadA* from genome (pEC2899)** | | | | |
| pEC801 linearisation | OLEC9943 | *CTGCAGGCATGCAAGCTTGCG* | F | PCR |
| | OLEC9944 | *TCTAGAGGATCCCCGGGTACCGAG* | R | PCR |
| Upstream fragment | OLEC12752 | ACCCGGGGATCCTCTAGAC*TGTGGCAGCCTAAAGTATTGC* | F | PCR |
| | OLEC12753 | TGTATGCTATACGAACGGTA*TCAAGAAAGACTCCCCAAGTTCC* | R | PCR |
| Downstream fragment | OLEC12754 | GCATACATTATACGAACGGTA*AGTTTTAAGGAGTTTGTTATCACATGC* | F | PCR |
| | OLEC12755 | AAGCTTGCATGCCTGCAG*TCAGATTGAGCACATCTAATGCTTG* | R | PCR |
| lox71-P$_{ermAM/B}$-*ermAM/B*-lox66 | OLEC1943 | *TACCGTTCGTATAGCATACATTATACGAAGTTATCCGTAGCGGTTTTCAAAATTTGCAACC* | F | PCR |
| | OLEC1932 | *TACCGTTCGTATAATGTATGCTATACGAAGTTATTTATTTCCTCCCGTTAAATAATAGATAACTATTAAA* | R | PCR |
| PCR ligation | OLEC11417 | *ACCCGGGGATCCTCTAGA* | F | LM-PCR |
| | OLEC11418 | *AAGCTTGCATGCCTGCAG* | R | LM-PCR |

| **Purpose** | **Code** | **Sequence 5′-3′[a]** | **F/R[b]** | **Usage[c]** |
|---|---|---|---|---|
| **Sanger sequencing of A34-to-I34 editing of transfer RNAs** | | | | |
| *E. coli argQ* / $tRNA^{Arg}_{ACG}$ | OLEC9821 | *GCATCCGTAGCTCAGCTGG* | F | RT-PCR |
| | OLEC9822 | *TGGTGCATCCGGGAGGATTC* | R | RT |
| | OLEC9823 | TAATACGACTCACTATAGGG*TGGTGCATCCGGGAGGATTC* | R | RT-PCR |
| *SPy_t37* / $tRNA^{Arg}_{ACG}$ | OLEC8776 | *GCACCCTTAGCTCAACTGG* | F | RT-PCR |
| | OLEC8777 | *TGGTGCACCCTAGAGGAG* | R | RT |
| | OLEC9171 | TAATACGACTCACTATAGGG*TGGTGCACCCTAGAGGAG* | R | RT-PCR |
| *SPy*_t52 / $tRNA^{Leu}_{AAG}$ | OLEC8778 | *GCGGGGATGGCGGAATTG* | F | RT-PCR |
| | OLEC8779 | *TGGTGCGGAGAGTGGGAC* | R | RT |
| | OLEC9172 | TAATACGACTCACTATAGGG*TGGTGCGGAGAGTGGGAC* | R | RT-PCR |
| all tRNAs | OLEC1889 | *TAATACGACTCACTATAGGG* | F | SEQ |
| Validation of locus | OLEC12750 | *GGATGCAGTGATTATTAGCC* | F | SEQ |
| | OLEC12751 | *GATATACTAACACAATTGAAGGC* | R | SEQ |
| **Suicide vector template for genomic integration of AHT-inducible promoter (pEC2901)** | | | | |
| pEC801 linearisation | OLEC9943 | *CTGCAGGCATGCAAGCTTGCG* | F | PCR |
| | OLEC9944 | *TCTAGAGGATCCCCGGGTACCGAG* | R | PCR |
| *cat86*-$P_{tet}$ | OLEC12291 | *GTCAATATTTTTTTTAGTTTTTCATGAACTCG* | R | PCR |
| | OLEC12294 | *CGAAAATTGGATAAAGTGGGATATTTTT* | F | PCR |
| TT3 insertion upstream of *cat86* | OLEC12295 | GGTACCCGGGGATCCTCTAGATGTACAAAAAAGGCCCACAAAAGTGGGCCTTTTTTCGAAAATTGGATAAAGTGGGATAT | F | OA |
| | OLEC12296 | ATATCCCACTTTATCCAATTTTCGAAAAAAGGCCCACTTTTGTGGGCCTTTTTTGTACATCTAGAGGATCCCCGGGTACC | R | OA |
| $P_{tet}$ | OLEC12292 | CTCGAGTTCATGAAAAACTAAAAAAAATATTGACACTCTATCATTGATAGAGTATAATTAAAATAAGACTCTATCATTGATAGAGTGTACAGTCGACCTGCAGGCATGCAAGCTTGCG | F | OA |
| | OLEC12293 | CGCAAGCTTGCATGCCTGCAGGTCGACTGTACACTCTATCAATGATAGAGTCTTATTTTAATTATACTCTATCAATGATAGAGTGTCAATATTTTTTTTAGTTTTTCATGAACTCGAG | R | OA |
| Sequencing of *cat86*-$P_{tet}$ cassette | OLEC12291 | *GTCAATATTTTTTTTAGTTTTTCATGAACTCG* | R | SEQ |
| | OLEC12294 | *CGAAAATTGGATAAAGTGGGATATTTTT* | F | SEQ |
| **EC2353 (SF370 $P_{tet}$-*rnjA*) via pEC812 and pEC852** | | | | |
| *tetR*(A582C) (pEC812) | OLEC3322 | *CCGCAGATGATCAATTCAAGG* | F | SDM-PCR |
| | OLEC3323 | *CCTTGAATTGATCATCTGCGG* | R | SDM-PCR |
| Upstream fragment (pEC852) | OLEC3524 | AAAAACTGCAG*TAGTGAAGCATGGGCTTGTG* | F | PCR |
| | OLEC3525 | AAAAACTGCAG*CTTAGAACTCCGTTAATTCAAAAACACC* | R | PCR |
| Downstream fragment (pEC852) | OLEC3526 | AAAACATATG*ACAAATATCAGTTTAAAACCTAATGAAGTTG* | F | PCR |
| | OLEC3527 | AAAAACATATG*CTGCCATAGACTCACCTTGACTACC* | R | PCR |
| **EC3453 (SF370 $P_{tet}$-*slo*) via pEC2964** | | | | |

| Purpose | Code | Sequence 5′-3′[a] | F/R[b] | Usage[c] |
|---|---|---|---|---|
| **Sanger sequencing of A34-to-I34 editing of transfer RNAs** | | | | |
| *E. coli argQ* / $tRNA^{Arg}_{ACG}$ | OLEC9821 | *GCATCCGTAGCTCAGCTGG* | F | RT-PCR |
| | OLEC9822 | *TGGTGCATCCGGGAGGATTC* | R | RT |
| | OLEC9823 | TAATACGACTCACTATAGGG*TGGTGCATCCGGGAGGATTC* | R | RT-PCR |
| *SPy_t37* / $tRNA^{Arg}_{ACG}$ | OLEC8776 | *GCACCCTTAGCTCAACTGG* | F | RT-PCR |
| | OLEC8777 | *TGGTGCACCCTAGAGGAG* | R | RT |
| | OLEC9171 | TAATACGACTCACTATAGGG*TGGTGCACCCTAGAGGAG* | R | RT-PCR |
| *SPy*_t52 / $tRNA^{Leu}_{AAG}$ | OLEC8778 | *GCGGGGATGGCGGAATTG* | F | RT-PCR |
| | OLEC8779 | *TGGTGCGGAGAGTGGGAC* | R | RT |
| | OLEC9172 | TAATACGACTCACTATAGGG*TGGTGCGGAGAGTGGGAC* | R | RT-PCR |
| all tRNAs | OLEC1889 | *TAATACGACTCACTATAGGG* | F | SEQ |
| Upstream fragment (pEC2964) | OLEC12908 | TACCCGGGGATCCTCTAGAT*CCGTGCTATCTTGTTGTCTATGG* | F | PCR |
| | OLEC12909 | CACTTTTGTGGGCCTTTTTT*AAGTAAACTATACATTAGCAAATAGTTTTTGTC* | R | PCR |
| Downstream fragment (pEC2964) | OLEC12910 | AATAAGACTCTATCATTGATAGAGT*AAAAATAATATAAGGTGGTTTACATGAGAAAC* | F | PCR |
| | OLEC12911 | TGCATGCCTGCAGGTCGACT*GATGGACCTCTGTTACCTCAATATC* | R | PCR |
| Validation of locus | OLEC12912 | *GGTTGAAATGGTCATGACAGATG* | F | PCR, SEQ |
| | OLEC12913 | *TCAGCATACTTGTCGGCAG* | R | PCR, SEQ |
| **EC3456 (SF370 $P_{tet}$-*amyA*) via pEC2965** | | | | |
| Upstream fragment (pEC2965) | OLEC12896 | TACCCGGGGATCCTCTAGAT*GACAACGAATTGCTAATCAGCTG* | F | PCR |
| | OLEC12897 | CACTTTTGTGGGCCTTTTTT*AAGTTAAGTTTATCGTAATCGCTTTCATTTG* | R | PCR |
| Downstream fragment (pEC2965) | OLEC12898 | AATAAGACTCTATCATTGATAGAGT*AAACGTTTAAGTTTCGAAACTATTACTG* | F | PCR |
| | OLEC12899 | TGCATGCCTGCAGGTCGACT*CAAAGTTTCAATTACGTCTGGTG* | R | PCR |
| Validation of locus | OLEC12900 | *CTTACCTAGATGCTCGTCG* | F | PCR, SEQ |
| | OLEC12901 | *CCGGCAATTAAACCATAACC* | R | PCR, SEQ |
| **EC3459 (SF370 $P_{tet}$-*fakB2*) via pEC2966** | | | | |
| Upstream fragment (pEC2966) | OLEC12891 | TACCCGGGGATCCTCTAGAT*GAGGACATTTAGATAACAAACTGGACG* | F | PCR |
| | OLEC12892 | CACTTTTGTGGGCCTTTTTT*CTCTTTATTCTAACACTCCTTACTAAATAAAG*c | R | PCR |
| Downstream fragment (pEC2966) | OLEC12893 | AATAAGACTCTATCATTGATAGAGT*GTGAGAAACTAGTTTAGAATGGAAAAATAC* | F | PCR |
| | OLEC12894 | TGCATGCCTGCAGGTCGACT*CATTAAGTAGCGAGCTCAAAACAC* | R | PCR |
| Validation of locus | OLEC12895 | *GAAGATACAATTGAGCCAGAGC* | F | PCR, SEQ |
| | OLEC11121 | *GAGTGGTTGTAATAAGCTGCG* | R | PCR, SEQ |
| **EC3570** (**SF370 $P_{tet}$-$TT_{tadA}$**) **via pEC3000** | | | | |

| **Purpose** | **Code** | **Sequence 5′-3′[a]** | **F/R[b]** | **Usage[c]** |
|---|---|---|---|---|
| **Sanger sequencing of A34-to-I34 editing of transfer RNAs** | | | | |
| *E. coli argQ* / tRNA$^{Arg}_{ACG}$ | OLEC9821 | *GCATCCGTAGCTCAGCTGG* | F | RT-PCR |
| | OLEC9822 | *TGGTGCATCCGGGAGGATTC* | R | RT |
| | OLEC9823 | TAATACGACTCACTATAGGG*TGGTGCATCCGGGAGGATTC* | R | RT-PCR |
| *SPy_t37* / tRNA$^{Arg}_{ACG}$ | OLEC8776 | *GCACCCTTAGCTCAACTGG* | F | RT-PCR |
| | OLEC8777 | *TGGTGCACCCTAGAGGAG* | R | RT |
| | OLEC9171 | TAATACGACTCACTATAGGG*TGGTGCACCCTAGAGGAG* | R | RT-PCR |
| *SPy*_t52 / tRNA$^{Leu}_{AAG}$ | OLEC8778 | *GCGGGGATGGCGGAATTG* | F | RT-PCR |
| | OLEC8779 | *TGGTGCGGAGAGTGGGAC* | R | RT |
| | OLEC9172 | TAATACGACTCACTATAGGG*TGGTGCGGAGAGTGGGAC* | R | RT-PCR |
| all tRNAs | OLEC1889 | *TAATACGACTCACTATAGGG* | F | SEQ |
| *cat86*-P$_{tet}$ | OLEC12294 | *CGAAAATTGGATAAAGTGGGATATTTTT* | F | PCR, SEQ |
| | OLEC12869 | *ATGAGATCACCTCCTTAACTAGAC* | R | PCR, SEQ |
| Upstream fragment (pEC2913) | OLEC12872 | ACCCGGGGATCCTCTAGAC*GTGAAGAGAGTAACCAAGCC* | F | PCR |
| | OLEC12868 | AATATCCCACTTTATCCAATTTTCG*GTTTTCATCTCGATTGGGTCTGA* | R | PCR |
| Downstream fragment (pEC2913) | OLEC12873 | CTAGTTAAGGAGGTGATCTCAT*AGTTTTAAGGAGTTTGTTATCACATGC* | F | PCR |
| | OLEC12755 | AAGCTTGCATGCCTGCAG*TCAGATTGAGCACATCTAATGCTTG* | R | PCR |
| PCR ligation (up, cassette, down) | OLEC11417 | ACCCGGGGATCCTCTAGA | F | LM-PCR |
| | OLEC11418 | AAGCTTGCATGCCTGCAG | R | LM-PCR |
| RBS mutagenesis (pEC3000) | OLEC13498 | TTTCCTC*TAACTAGACTCGAAGATCTATTCGAG* | R | SDM-PCR |
| | OLEC13499 | *TCGAGTCTAGTTA*GAGGAAA*AGTTTTAAGGAGTTTGTTATCACATGC* | F | SDM-PCR |
| **EC3622 (SF370 P$_{tet}$-*tadA*-TT$_{tadA}$) via pEC3021** | | | | |
| *cat86*-P$_{tet}$ | OLEC12294 | *CGAAAATTGGATAAAGTGGGATATTTTT* | F | PCR, SEQ |
| | OLEC12869 | *ATGAGATCACCTCCTTAACTAGAC* | R | PCR, SEQ |
| Upstream fragment (pEC2914) | OLEC12752 | ACCCGGGGATCCTCTAGAC*TGTGGCAGCCTAAAGTATTGC* | F | PCR |
| | OLEC12866 | CAAACTCCTTAAAACT*CAAGAAAGACTCCCCAAGTTCC* | R | PCR |
| TT$_{tadA}$ | OLEC12867 | GAGTCTTTCTTGA*GTTTTAAGGAGTTTGTTATCACATGC* | F | PCR |
| | OLEC12868 | AATATCCCACTTTATCCAATTTTCG*GTTTTCATCTCGATTGGGTCTGA* | R | PCR |
| Downstream fragment (pEC2914) | OLEC12870 | CTAGTTAAGGAGGTGATCTCAT*ATGCCATATAGTTTAGAAGAGCAAAC* | F | PCR |
| | OLEC12871 | AAGCTTGCATGCCTGCAG*CTAGTCAAAGGGATCTGACTGTTC* | R | PCR |
| PCR ligation (up, cassette, down) | OLEC11417 | ACCCGGGGATCCTCTAGA | F | LM-PCR |
| | OLEC11418 | AAGCTTGCATGCCTGCAG | R | LM-PCR |

| Purpose | Code | Sequence 5′-3′[a] | F/R[b] | Usage[c] |
|---|---|---|---|---|
| **Sanger sequencing of A34-to-I34 editing of transfer RNAs** | | | | |
| *E. coli argQ* / tRNA$^{Arg}_{ACG}$ | OLEC9821 | *GCATCCGTAGCTCAGCTGG* | F | RT-PCR |
| | OLEC9822 | *TGGTGCATCCGGGAGGATTC* | R | RT |
| | OLEC9823 | TAATACGACTCACTATAGGG*TGGTGCATCCGGGAGGATTC* | R | RT-PCR |
| *SPy_t37* / tRNA$^{Arg}_{ACG}$ | OLEC8776 | *GCACCCTTAGCTCAACTGG* | F | RT-PCR |
| | OLEC8777 | *TGGTGCACCCTAGAGGAG* | R | RT |
| | OLEC9171 | TAATACGACTCACTATAGGG*TGGTGCACCCTAGAGGAG* | R | RT-PCR |
| *SPy*_t52 / tRNA$^{Leu}_{AAG}$ | OLEC8778 | *GCGGGGATGGCGGAATTG* | F | RT-PCR |
| | OLEC8779 | *TGGTGCGGAGAGTGGGAC* | R | RT |
| | OLEC9172 | TAATACGACTCACTATAGGG*TGGTGCGGAGAGTGGGAC* | R | RT-PCR |
| all tRNAs | OLEC1889 | *TAATACGACTCACTATAGGG* | F | SEQ |
| RBS mutagenesis (pEC3001) | OLEC13498 | TTTCCTC*TAACTAGACTCGAAGATCTATTCGAG* | R | SDM-PCR |
| | OLEC13500 | *TCGAGTCTAGTTA*GAGGAAAA*TGCCATATAGTTTAGAAGAGCAAAC* | F | SDM-PCR |
| RBS mutagenesis (pEC3021) | OLEC13766 | T*TTTTCCTCTAACTAGACTCGAAGATCTATTCGAGC* | R | SDM-PCR |
| | OLEC13767 | *TCGAGTCTAGTTAGAGGAAAAAATGCCATATAGTTTAGAAGAGCAAAC* | F | SDM-PCR |
| **Verification of virulence regulator gene integrity in mutant strains** | | | | |
| *covRS* | OLEC4856 | *TCGCTAGAAGACTATTTGACCAT* | F | PCR, SEQ |
| | OLEC4867 | *AAGACATCGCGATTGACAGT* | R | PCR, SEQ |
| | OLEC3609 | *GGCTATGTTCAAGTCTTTCATG* | F | SEQ |
| | OLEC3610 | *CCAAATAACTCAACAACTAGTAGC* | R | SEQ |
| *mga* | oliRN172 | *AGTTGACTAACCAATTGATCTACGCCTTTT* | F | PCR, SEQ |
| | OLEC290 | *TTAACCTCTGTTTGATTCGC* | R | PCR, SEQ |
| *ropB* | OLEC4854 | *AGCGACTATCATCCGAAACAT* | F | PCR, SEQ |
| | OLEC4855 | *GCCCTGGAGCTGTTGAGATA* | R | PCR, SEQ |

[a] *italic*, sequence annealing to the template; underlined, restriction site; **bold**, T7 promoter.

[b] F, forward primer; R, reverse primer.

[c] LM-PCR, PCR-mediated ligation; NB, probe for Northern blot; SEQ, sequencing; SDM-PCR, PCR-mediated site-directed mutagenesis; RT-PCR, reverse transcription-PCR; OA, oligo assembly

**Table S3.** Plasmids used in this study.

| Plasmids | Relevant characteristics | Source |
|---|---|---|
| **Vector backbones for *S. pyogenes*** | | |
| pLZ12Km2-P*23R*:TA:*ffluc* (pEC2173) | pSH71, *aphIII*, ω-ε-ζ TA cassette, $P_{23R}$-*ffluc* | Addgene #88900 (16) |
| **Deletion of *tadA*** | | |
| pEC801 | pRO1600/ColE1, *bla* | SEVA (17) |
| pEC2899 | pEC801Ω*tadA*(up)-lox71-$P_{ermAM/B}$-*ermAM/B*-lox66-*tadA*(down) | This study |
| **Integration of AHT-inducible promoter into the genome of *S. pyogenes*** | | |
| pEC536 | ColE1-*repDEG*, *cat86*, $P_{tet}$ | pEU8517 (2) |
| pEC808 | pEC801Ω*cat86*-$P_{tet}$ | This study |
| pEC812 | pEC801Ω*cat86*-$P_{tet}$* | This study |
| pEC852 | pEC801Ω*rnjA*(up)-*cat86*-$P_{tet}$*-*rnjA*(1..925) | This study |
| pEC2901 | pEC801ΩTT3-*cat86*-$P_{tet}$ | This study |
| pEC2964 | pEC801Ω$P_{nga}$(up)-TT3-*cat86*-$P_{tet}$-$P_{nga}$(down) | This study |
| pEC2965 | pEC801Ω$P_{malX}$(up)-TT3-*cat86*-$P_{tet}$-$P_{malX}$(down) | This study |
| pEC2966 | pEC801Ω$P_{fakB2}$(up)-TT3-*cat86*-$P_{tet}$-$P_{fakB2}$(down) | This study |
| pEC2913 | pEC801Ω*tadA*-TT3-*cat86*-$P_{tet}$-*tadA*(down) | This study |
| pEC2914 | pEC801Ω*tadA*(up)-TT3-*cat86*-$P_{tet}$-*tadA* | This study |
| pEC3000 | pEC801Ω*tadA*-TT3-*cat86*-$P_{tet}$(RBSmut)-*tadA*(down) | This study |
| pEC3001 | pEC801Ω*tadA*(up)-TT3-*cat86*-$P_{tet}$(RBSmut)-*tadA* | This study |
| pEC3021 | pEC801Ω*tadA*(up)-TT3-*cat86*-$P_{tet}$(RBSmut2)-*tadA* | This study |
| **TadA protein production** | | |
| pET-21a(+) | pBR322, *bla*, *lacI*, $P_{T7}$(lacO) | Novagen |
| pEC2360 | pET-21a(+)Ω*tadA*(*Spy*) | This study |
| pEC2389 | pET-21a(+)Ω*tadA*(*Eco*) | This study |
| ***In vitro* transcription** | | |
| pUC19 | pMB1, *bla*, $P_{lac}$(lacO)-*lacZ*$\alpha$ | NEB |
| pEC2322 | pUC19Ω$P_{T7}$-tRNA-Leu-AAG(*Spy*) | This study |
| pEC2405 | pUC19Ω$P_{T7}$-tRNA-Arg-ACG(*Spy*) | This study |
| pEC2406 | pUC19Ω$P_{T7}$-tRNA-Arg-ACG(*Eco*) | This study |
| **Ectopic expression of *tadA*** | | |
| pEC2812 | pLZ12Km2-TA:$P_{gyrA}$(*Sag*)-TT3 | This study |
| pEC2813 | pLZ12Km2-TA:$P_{gyrA}$(*Sag*)-*tadA*(*Spy*):$His_6$-TT3 | This study |

| | | |
|---|---|---|
| ***ermBL* reporter assay** | | |
| pEC3045 | pLZ12Km2-TA:$P_{gyrA}$(*Sag*)-*ermBL*-*ermB*':A2I:*ffluc*-TT3 | This study |
| pEC3046 | pLZ12Km2-TA:$P_{gyrA}$(*Sag*)-*ermBL*(M1*)-*ermB*':A2I:*ffluc*-TT3 | This study |
| pEC3047 | pLZ12Km2-TA:$P_{gyrA}$(*Sag*)-*ermB*':A2I:*ffluc*-TT3 | This study |
| pEC3069 | pLZ12Km2-TA:$P_{gyrA}$(*Sag*)-*ermB*'(M1*):A2I:*ffluc*-TT3 | This study |

Promoters are denoted as P with the related gene name. The erythromycin resistance cassette ($P_{ermAM/B}$-*ermAM/B*) is flanked by lox71 and lox66 sites for Cre recombinase-based excision. The AHT-inducible $P_{tet}$ cassette harbours the repressor *tetR* and two divergent promoters with three operator sites, and is separated from the chloramphenicol resistance gene *cat86* by two T4 terminators as previously described (1, 2). A mutant version of $P_{tet}$ with *tetR*(A582C) is shown as $P_{tet}$*. Mutations of the ribosomal binding site as shown in Figure S1 are described as $P_{tet}$(RBSmut). Transcriptional terminators are denoted as TT, with TT3 as the phage T3 terminator. Species are indicated in brackets after the respective genes if applicable (*Eco*: *E. coli*; *Sag*: *S. agalactiae*; *Spy*: *S. pyogenes*).

**Table S10.** Differential expression of selected oxidative stress signature genes upon exposure to $H_2O_2$.

| | | | **0.5 mM $H_2O_2$** | | **1.0 mM $H_2O_2$** | |
|---|---|---|---|---|---|---|
| **Locus tag** | **Gene** | **Function** | **15 min** | **30 min** | **15 min** | **30 min** |
| **Peroxide detoxification** | | | | | | |
| *SPy_1406* | *sodA* | superoxide dismutase | 2.39 | 1.92 | 2.86 | 2.72 |
| *SPy_2079* | *ahpC* | alkyl hydroperoxide reductase C | 1.48 | 1.27 | 2.07 | 1.58 |
| *SPy_2080* | *ahpF* | alkyl hydroperoxide reductase F | 1.33 | 1.41 | 1.80 | 1.57 |
| **DNA repair and protein quality control** | | | | | | |
| *SPy_0185* | *polA* | DNA damage-induced DNA polymerase I | 1.18 | n.s. | 1.93 | 1.42 |
| *SPy_2216* | *htrA* | membrane-anchored quality control protease | n.s. | n.s. | 1.03 | 1.39 |
| **Metal ion homeostasis** | | | | | | |
| *SPy_0453* | *mtsA* | metal ABC transporter subunit | n.s. | n.s. | –1.15 | –1.28 |
| *SPy_1434* | *zntA* | heavy metal transporter ATPase | –1.38 | –2.10 | n.s. | –1.72 |
| *SPy_1531* | *dpr* | ferritin-like iron-binding protein | 2.58 | 1.59 | 3.17 | 3.09 |
| *SPy_1798* | *shr* | heme-binding protein | n.s. | n.s. | –1.26 | –1.60 |

Gene expression for each time point and $H_2O_2$ concentration was compared relative to the mock-treated control. Genes were considered differentially expressed with an absolute $\log_2$ fold change of at least 1 and an adjusted p value below 0.05. n.s.: not significant. Compare Table S9 for full differential expression analysis.

**SUPPLEMENTARY TABLE LEGENDS**

**Table S4.** A-to-I editing in the transcriptome of *S. pyogenes SF370*. Identified A-to-I editing sites were grouped according to their potential effect on the coding sequence (synonymous *vs.* non-synonymous). For each genomic position, locus tag, name (if available), function of the affected gene and the encoded amino acid are shown with editing levels (in %) for each replicate and growth phase. '—' indicates that editing levels could not be determined.

**Table S5.** A-to-I editing in the transcriptome of *S. pyogenes* SF370 upon ectopic *tadA* overexpression. Experiments were performed as described for Figure S3 and data are presented as for Table S4.

**Table S6.** A-to-I editing in the transcriptomes of *S. pyogenes* 5448 and 5448AP. A-to-I editing sites in the transcriptomes of *S. pyogenes* 5448 and 5448AP were identified in duplicates for three growth phases. Data are presented as in Table S4 with the corresponding locus tag in strains 5448 and SF370 for better comparison.

**Table S7.** A-to-I editing in the transcriptome of *S. pyogenes* SF370 in three different culture media. *S. pyogenes* SF370 was grown to mid-logarithmic growth phase in chemically defined medium (CDM), THY and C medium in triplicate, and data are presented as in Table S4.

**Table S8.** A-to-I editing in the transcriptome of *S. pyogenes* SF370 in response to hydrogen peroxide. *S. pyogenes* SF370 was grown to mid-logarithmic growth phase in C medium and exposed to 0.5 mM and 1.0 mM $H_2O_2$ with water as mock control (0.0 mM) for 15 min and 30 min. Data are presented as in Table S4.

**Table S9.** Genes differentially expressed in the transcriptome of *S. pyogenes* SF370 in response to hydrogen peroxide. Experiment was performed as described for Table S8, and differential expression analysis was performed with mock-treated samples as controls for each time point and $H_2O_2$ concentration. $Log_2$ fold changes and adjusted p-values are indicated for all significant hits (adjusted p-value < 0.05 and an absolute $log_2$ fold change of at least 1). n.s.: not significant.

**Table S11.** A-to-I editing in the transcriptome of *S. pyogenes SF370* $P_{tet}$-*rnjA* in the presence and absence of AHT. *S. pyogenes SF370* $P_{tet}$-*rnjA* was grown in THY supplemented with ("induced") or without ("uninduced") 0.1 ng/mL AHT to mid-logarithmic growth phase. Data are presented as in Table S4.
